# Long-read error correction: a survey and qualitative comparison

**DOI:** 10.1101/2020.03.06.977975

**Authors:** Pierre Morisse, Thierry Lecroq, Arnaud Lefebvre

## Abstract

Third generation sequencing technologies Pacific Biosciences and Oxford Nanopore Technologies were respectively made available in 2011 and 2014. In contrast with second generation sequencing technologies such as Illumina, these new technologies allow the sequencing of long reads of tens to hundreds of kbp. These so called long reads are particularly promising, and are especially expected to solve various problems such as contig and haplotype assembly or scaffolding, for instance. However, these reads are also much more error prone than second generation reads, and display error rates reaching 10 to 30%, according to the sequencing technology and to the version of the chemistry. Moreover, these errors are mainly composed of insertions and deletions, whereas most errors are substitutions in Illumina reads. As a result, long reads require efficient error correction, and a plethora of error correction tools, directly targeted at these reads, were developed in the past ten years. These methods can adopt a hybrid approach, using complementary short reads to perform correction, or a self-correction approach, only making use of the information contained in the long reads sequences. Both these approaches make use of various strategies such as multiple sequence alignment, de Bruijn graphs, Hidden Markov Models, or even combine different strategies. In this paper, we describe a complete survey of long-read error correction, reviewing all the different methodologies and tools existing up to date, for both hybrid and self-correction. Moreover, the long reads characteristics, such as sequencing depth, length, error rate, or even sequencing technology, have huge impacts on how well a given tool or strategy performs, and can thus drastically reduce the correction quality. We thus also present an in-depth benchmark of available long-read error correction tools, on a wide variety of datasets, composed of both simulated and real data, with various error rates, coverages, and read lengths, ranging from small bacterial to large mammal genomes.

## 1 Introduction

Since their inception in 2011 and 2014, third generation sequencing technologies Pacific Biosciences (PacBio) and Oxford Nanopore Technologies (ONT) became widely used and allowed the sequencing of massive amount of data. These technologies distinguish themselves from second generation sequencing technologies, such as Illumina, by the fact that they allow to produce much longer reads, reaching length of tens of kbp on average, and up to 1 million bps [31]. Thanks to their length, these so called long reads are expected to solve various problems, such as contig and haplotype assembly of large and complex organisms, scaffolding, or even structural variant calling, for instance. These reads are however extremely noisy, and display error rates of 10 to 30%, while second generation short reads usually reach error rates of around 1%. Moreover, long reads error are mainly composed of insertions and deletions, whereas short reads mainly contain substitutions. As a result, in addition to a higher error rate, the error profiles of the long reads are also much more complex than the error profiles of the short reads. In addition, ONT reads also suffer from bias in homopolymer regions, and thus tend to contain systematic errors in such regions, when they reach more than 6 bps. As a consequence, error correction is often used as a first step in projects dealing with long reads. Since the error profiles and error rates of the long reads are much different than those of the short reads, this necessity led to new algorithmic developments, specifically targeted at these long reads.

Two major ways of approaching long-read correction were thus developed. The first one, hybrid correction, makes use of additional short read data to perform correction. The second one, self-correction, on the contrary, attempts to correct long reads solely based on the information contained in their sequences. Both these approaches rely on various strategies, such as multiple sequence alignment, de Bruijn graphs, or Hidden Markov Models, for instance. As a result, a plethora of long-read correction methods were developed since 2012, and thirty different tools are available as of today.

### 1.1 Contribution

In this paper, we propose a complete survey of long-read error correction tools. Previous papers already covered this topic. On the one hand, [24] and [70] focused on hybrid correction. The former proposed a benchmark of ten tools on five datasets, whereas the latter studied performances of graph-based and alignment-based methods. On the other hand, [50] covered both hybrid and self-correction, but focused on RNA-sequencing data. Here, we focus on DNA-sequencing, and draw-up a summary of every single approach described in the literature, both for hybrid and self-correction. In addition, we also provide a list of all the available methods, and describe the strategies they rely on. We thus propose the most complete survey on long-read correction up to date.

Additionally, long reads characteristics, such as the sequencing depth, the length, the error rate, and the sequencing technology, can impact how well a given tool or strategy performs. As a result, a given tool performing the best on a given dataset does not mean this same tool will perform as well on other datasets, especially if their characteristics fluctuate from one another. Therefore, we also present an in-depth benchmark of all available long-read correction tools, on a total of 19 different datasets, displaying diverse characteristics. In particular, we assess both simulated and real data, and rely on datasets having varying read lengths, error rates, and sequencing depths, ranging from smaller bacterial to large mammal genomes. Compared to previous papers, we thus propose the most extensive benchmark up to date.

## 2 State-of-the-art

As mentioned in Section 1, the literature describes two main approaches to tackle long-read error correction. On the one hand, hybrid correction makes use of complementary, high quality, short reads to perform correction. On the other hand, self-correction attempts to correct the long reads solely using the information contained in their sequences.

One of the major interests of hybrid correction is that error correction is mainly guided by the short read data. As a result, the sequencing depth of the long reads has no impact on this strategy whatsoever. This way, datasets composed of a very low coverage of long reads can still be efficiently corrected using a hybrid approach, as long as the sequencing depth of the short reads remains sufficient, *i.e.* around 50x.

Contrariwise, self-correction is purely based on the information contained in the long reads. As a result, deeper long read coverages are usually required, and self-correction can thus prove to be inefficient when dealing with datasets displaying low coverages. The required sequencing depth to allow for an efficient self-correction is however reasonable, as it has been shown that from a coverage of 30x, self-correction methods are able to provide satisfying results [66].

In this section, we present the state-of-the-art of available long-read error correction methods. More particularly, we describe the various methodologies adopted by the different tools, and list the tools relying on each methodology, both for hybrid and self-correction. Details about performances, both in terms of resource consumption and quality of the results, are however not discussed here. Experimental results of a subset of available correction tools, on various datasets displaying diverse characteristics, are presented in Section 3. A summary of available hybrid correction tools is given Table 1. A summary of available self-correction tools is given Table 2.

**Table 1:** List of available hybrid correction tools. Complete reads correspond to reads which are neither trimmed nor split. The case column indicates whether the method reports corrected bases in a different case from the uncorrected bases.

| Method | Approach | Output reads |  |  | Case | Release | Validated on |
| --- | --- | --- | --- | --- | --- | --- | --- |
|  |  | Complete | Trimmed | Split |  |  |  |
| PBcR | Short reads alignment | ✗ | ✗ | ✓ | ✗ | 2012 | PacBio |
| LSC | Short reads alignment | ✗ | ✓ | ✗ | ✗ | 2012 | PacBio |
| ECTools | Contigs alignment | ✗ | ✗ | ✓ | ✗ | 2014 | PacBio |
| LoRDEC | De Bruijn graphs | ✓ | ✗ | ✗ | ✓ | 2014 | PacBio |
| Proovread | Short reads alignment | ✓ | ✗ | ✓ | ✗ | 2014 | PacBio |
| Nanocorr | Short reads alignment | ✓ | ✗ | ✗ | ✗ | 2015 | ONT |
| NaS | Short reads alignment | ✓ | ✗ | ✗ | ✗ | 2015 | ONT |
| CoLoRMap | Short reads alignment | ✓ | ✗ | ✗ | ✓ | 2016 | PacBio |
| Jabba | De Bruijn graphs | ✗ | ✗ | ✓ | ✗ | 2016 | PacBio |
| LSCplus | Short reads alignment | ✓ | ✓ | ✗ | ✗ | 2016 | PacBio |
| HALC | Contigs alignment | ✓ | ✓ | ✓ | ✓ | 2017 | PacBio |
| HECIL | Short reads alignment | ✓ | ✗ | ✗ | ✗ | 2018 | PacBio |
| Hercules | Hidden Markov Models | ✓ | ✗ | ✗ | ✗ | 2018 | PacBio |
| FMLRC | De Bruijn graphs | ✓ | ✗ | ✗ | ✗ | 2018 | PacBio |
| HG-CoLoR | Short reads alignment<br>+ de Bruijn graphs | ✓ | ✓ | ✓ | ✓ | 2018 | PacBio + ONT |
| MiRCA | Contigs alignment | ✗ | ✓ | ✗ | ✗ | 2018 | ONT |
| ParLECH | De Bruijn graphs | ✓ | ✗ | ✗ | ✗ | 2019 | PacBio |

**Table 2:** List of available self-correction tools. Complete reads correspond to reads which are neither trimmed nor split. The case column indicates whether the method reports corrected bases in a different case from the uncorrected bases.

| Method | Approach | Output reads |  |  | Case | Release | Validated on |
| --- | --- | --- | --- | --- | --- | --- | --- |
|  |  | Complete | Trimmed | Split |  |  |  |
| PBcR-BLASR | Multiple sequence alignment | ✗ | ✗ | ✓ | ✗ | 2013 | PacBio |
| PBDAGCon | Multiple sequence alignment | ✓ | ✗ | ✗ | ✗ | 2013 | PacBio |
| Sprai | Multiple sequence alignment | ✓ | ✗ | ✗ | ✗ | 2014 | PacBio |
| PBcR-MHAP<br>(PBDAGCon) | Multiple sequence alignment | ✓ | ✗ | ✗ | ✗ | 2015 | PacBio |
| PBcR-MHAP<br>(FalconSense) | Multiple sequence alignment | ✗ | ✗ | ✓ | ✗ | 2015 | PacBio |
| FalconSense | Multiple sequence alignment | ✗ | ✗ | ✓ | ✗ | 2016 | PacBio |
| LoRMA | De Bruijn graphs | ✗ | ✗ | ✓ | ✗ | 2016 | PacBio |
| Sparc | Multiple sequence alignment | ✓ | ✗ | ✗ | ✗ | 2016 | PacBio + ONT |
| Canu | Multiple sequence alignment | ✗ | ✗ | ✓ | ✗ | 2017 | PacBio + ONT |
| Daccord | De Bruijn graphs | ✗ | ✗ | ✓ | ✗ | 2017 | PacBio + ONT |
| MECAT | Multiple sequence alignment | ✗ | ✗ | ✓ | ✗ | 2017 | PacBio + ONT |
| CONSENT | Multiple sequence alignment<br>+ de Bruijn graphs | ✗ | ✓ | ✗ | ✓ | 2019 | PacBio + ONT |
| FLAS | Multiple sequence alignment | ✗ | ✓ | ✓ | ✗ | 2019 | PacBio |

### 2.1 Hybrid correction

Hybrid correction was the first approach to be described in the literature. This strategy is based on a set of long reads and a set of short reads, both sequenced for the same individual. It aims to use the high quality information contained in the short reads to enhance the quality of the long reads. As first long read sequencing experiments displayed high error rates (*>* 15% on average), most methods relied on this additional use of short reads data. Four different hybrid correction approaches thus exist:

1. Alignment of short reads to the long reads. Once the short reads are aligned, the long reads can indeed be corrected by computing a consensus sequence from the subset of short reads associated to each long read. PBcR / PacBioToCA [37], LSC [4], Proovread [27], Nanocorr [25], LSCplus [29], CoLoRMap [28], and HECIL [16] are all based on this approach.
2. Alignment of long reads and contigs obtained from short reads assembly. In the same fashion, long reads can also be corrected with the help of the contig they align to, by computing consensus sequences from these contigs. ECTools [45], HALC [6], and MiRCA [32] adopt this methodology.
3. Use of de Bruijn graphs, built from the short reads’ *k*-mers. Once built, the long reads can indeed be anchored to the graph. It can then be traversed, in order to find paths allowing to link together anchored regions of the long reads, and thus correct unanchored regions. LoRDEC [63], Jabba [56], FMLRC [71], and ParLECH [17] rely on this strategy.
4. Use of Hidden Markov Models. These can indeed be used in order to represent the long reads. The models can then be trained with the help of short reads, in order to extract consensus sequences, representing the corrected long reads. Hercules [22] is based on this approach.

Other methods, such as NaS [51] and HG-CoLoR [57], combine different of the aforementioned approaches in order to counterbalance their advantages and their drawbacks. We describe each approach more into details, and summarize the core algorithm of each tool, in the following subsections.

#### 2.1.1 Short reads alignment

This approach was the first long-read error correction approach described in the literature. It consists of two distinct steps. First, the short reads are aligned to the long reads. This step allows to cover each long reads with a subset of related short reads. This subset can then be used to compute a high quality consensus sequence, which can, in turn, be used as the correction of the original long read. The different methods adopting this approach mainly vary by the alignment methods they use, and also by algorithmic choices made during the consensus sequences computation.

#### PBcR / PacBioToCA (2012)

PBcR can compute alignment with the help of the aligner developed and included in the tool, with BLASR [12], or with Bowtie [42]. In order to reduce computational times, and avoid unsignificant alignments, alignments are only computed between short reads and long reads that share a sufficient number of *k*-mers. Moreover, only the short reads that align throughout their whole length are considered for the correction of the long reads they are associated to. Each short read can thus be aligned to multiple long reads. However, on a same, given long read, a short read is only considered once, for the correction of the region to which it aligns with the highest identity.

Moreover, to avoid covering repeated regions of different long reads with the same subset of short reads, the short reads are associated to the repeats they are more likely to belong to, according to the identity of the alignments. Each short read is thus only authorized to participate to the correction of the *N* long reads to which it aligns with the highest identity, where *N* roughly represents the sequencing depth of the long reads. Thus, the different repeats are only covered by the short reads representing them the best.

Finally, a consensus sequence is generated from the subset of short reads covering each long read, with the help of the consensus module included in the assembler AMOS [62]. This module computes the consensus sequence in multiple steps, in a iterative way. At each step, all the short reads are aligned to the current consensus (being the long read itself during the first step), in order to generate a multiple sequence alignment (MSA). This MSA is then used to compute a new consensus sequence, with a majority vote at each position. Iterations stop when the new consensus sequence is identical to the previous step’s consensus sequence.

PBcR thus produces split corrected long reads. Indeed, extremities of the long reads that are not covered by alignments are not included in the consensus sequence computation, and are thus not reported in the corrected versions of the long reads. Moreover, if gaps of coverage are spotted along a long read sequence, the corrected version of this read is split, and each covered region is reported independently, in order to avoid introducing incorrect bases in the corrected long reads.

#### LSC (2012)

LSC starts by compressing homopolymers regions, both in the short reads and in the long reads. To this aim, homopolymers are replaced by a single occurrence of the repeated nucleotide. For instance, the sequence CTTAGGA is compressed as CTAGA. This compression step is used to facilitate the alignments computation. Moreover, in order to be able to perform the symmetric operation, and decompress the reads, an index is also computed. This index is composed of two arrays for each compressed read, associating the position of the homopolymers to the length of these homopolymers in the original read. Once the reads are compressed, the long reads are concatenated, in order to obtain larger sequences. During this concatenation step, consecutive reads are separated by a block of *n* nucleotides *N*, where *n* represents the average length of the short reads.

Short reads are then aligned to these new sequences, obtained after concatenation. By default, alignments are computed with Novoalign^1^, although LSC is also compatible with other, less resource-consuming aligners, such as BWA [49] or SeqAlto [59]. After this alignment step, a subset of short reads if associated to each long read.

Each long is then corrected by computing local consensus sequences from the short reads, on regions where alignments define substitutions, deletions or insertions between the long read and the short reads. Moreover, compressed homopolymers regions, whether they are located on the long read or on the short reads, are also locally corrected in the same fashion. For each aforementioned region, the sequences of all the short reads covering the region are temporarily decompressed. A local consensus sequence is thus computed with a majority vote, in order to correct this region in the long read. The bases of the long read that do not correspond to any of these regions are thus left unmodified by the correction step. Once all the regions requiring correction have been processed, compressed homopolymers regions of the long reads, unmodified during the correction step, are decompressed and left unmodified in the sequences of the corrected long reads.

LSC thus produces trimmed corrected long reads, defined, for each read, as the sequence of the uncompressed long read, from the leftmost base to the rightmost base, covered by at least one short read. However, bases of the long reads that could not be corrected are not reported in a different case from the corrected bases. As a result, the corrected long reads cannot further be split after correction.

#### Proovread (2014)

Proovread computes alignments with the help of SHRiMP2 [18]. Although this tool is used as the default aligner, all the aligners reporting their results in SAM format, like BWA, Bowtie or Bowtie2 [41] are compatible with Proovread and can easily be used. A validation step is then applied, in order to filter out low quality alignments. To this aim, Proovread uses scores that are normalized according to the length of the alignments, in a local context. For this purpose, the long reads are represented as a series of consecutive, small windows of fixed length *L* (*L* = 20 by default). Each alignment is thus assigned to one of these windows, according to the position of its center.

Once the alignments have been allocated to the different windows, only the alignments having the highest scores in each window are considered for the consensus sequence computation, during the next step. To compute this consensus sequence, a matrix composed of one column per nucleotide is used for each long read. This matrix is filled with the bases from the short reads, according to the information contained in the alignments selected during the previous step. The consensus sequence is then computed with a majority vote on each column of the matrix. If no base coming from a short read is available for a given column, the base of the original long read is conserved. Moreover, a score mimicking the Phred quality score is assigned to each base of the corrected long read. This score is defined from the support of the base that was chosen as the consensus, in the corresponding column of the matrix.

In order to reduce time and memory consumption of the alignment step, which can be extremely costly when the required level of sensitivity is high, Proovread proposes an iterative procedure for alignment and correction, where the alignment sensitivity is increased at each step. This strategy is composed of three cycles of pre-correction and of a finishing step. Thus, during the pre-correction cycles, only a subset of 20, 30, and 50% of the original set of reads is used, respectively for the first, second, and third cycle. During the first cycle, the short reads are quickly aligned, with a moderate level of sensitivity. Long reads are then pre-corrected with the help of the aligned short reads, as described in the previous paragraph. Moreover, regions of the long reads that are sufficiently covered by the short reads are masked, and are thus not further considered in the following alignment and correction cycles. Two additional pre-correction cycles are then applied, using larger subsets of the short reads, and increasing levels of sensitivity. On average, the first two cycles are enough to mask more than 80% of the long reads sequencing. Finally, for the finishing step, the masks are removed from the pre-corrected long reads, and the complete set of short reads is aligned with a high sensitivity, in order to perform a last correction step, and thus polish previous pre-corrections.

Proovread thus produces corrected long reads both in their trimmed version and in their split version. The splitting step is indeed not dependent on the correction step, and relies on the pseudo-Phred quality scores assigned to each bases of the corrected long reads. Low quality regions of the corrected long reads can thus easily be identified and removes, in order to only retain high quality regions.

#### Nanocorr (2015)

Nanocorr computes alignments with the help of BLAST [2]. A set of alignments, composed of both correct alignments, spanning almost through the whole length of the short reads, and incorrect of partial alignments of the short reads is thus obtained. A first filtering step is then applied to these alignments, in order to remove alignments that are too short, and fully included in another, larger alignment. A dynamic programming algorithm, based on the longest increasing subsequences problem [65] is then applied, in order to select the optimal subset of short reads alignments covering each long read.

From this subset, a consensus sequence is computed, with the help of a directed acyclic graph (DAG), using the consensus module PBDAGCon, described Section 2.2.1.

Nanocorr thus produces untrimmed corrected long reads. Moreover, bases of the long reads that could not be corrected are not reported in a different case from the corrected bases, for the reasons mentioned in the description of PBDAGCon, Section 2.2.1. As a result, the corrected long reads cannot further be trimmed or split after correction.

#### LSCplus (2016)

LSCplus is based on the same principle as LSC, which we previously presented. It however brings a few modifications in terms of implementation and optimization during the alignment and correction steps. First, as in LSC, the homopolymers regions, both in the short reads and in the long reads, are compressed and replaced by a single occurrence of the repeated nucleotide. The index allowing to retrieve the original size of the homopolymers is however altered compared to that of LSC. Indeed, in LSCplus, this index is composed of a single array for each compressed read, registering, for each position (whether it is compressed or not), the number of bases in the original read. Moreover, unlike LSC, the long reads are not concatenated in larger sequences before performing the alignment step. The short reads are thus directly aligned to the long reads, with the help of Bowtie2.

Each long read can then be corrected, using the subset of short reads associated to it. A consensus sequence is thus computed, with a majority vote, at each position of the compressed long read. Unlike LSC, which decompresses regions requiring correction before computing the consensus, the most frequent nucleotide at each position is determined before decompression, in LSCplus. Once the most frequent nucleotide has been chosen, LSCplus performs decompression, and the size of the homopolymer is determined from the information contained in the indexes of the short reads that align at this position. The sequence thus obtained is concatenated at the end of the global consensus sequence, which is iteratively built, as the compressed long read bases are processed. Bases corresponding to uncovered positions of the long reads are decompressed without further modifications, and concatenated at the end of the global consensus sequence as well.

Unlike LSC, LSCplus thus produces corrected long reads both in their trimmed and in their untrimmed versions. As for LSC, the sequence of a trimmed long read is defined as the sequence of the uncompressed long read, from the leftmost base to the rightmost base, covered by at least one short read. Once again, as for LSC, bases of the long reads that could not be corrected are not reported in a different case from the corrected bases. As a result, the corrected long reads cannot further be split after correction.

#### CoLoRMap (2016)

CoLoRMap computes alignments with the help of BWA-MEM [46]. An alignment graph is then built for each long read, from the subset of short reads that align to it. This graph is directed and weighted, and each vertex is defined from the alignment of a short read to the long read. An edge is defined between two vertices *u* and *v* if the short reads associated to these two vertices share a prefix / suffix overlap, over a sufficient length, and with at most one error. At last, the weight of an edge between two vertices *u* and *v* is defined as the edit distance between the suffix of the read associated to *v* that is not included in the overlap, and the original long read.

Each connected component of this graph is then processed independently. For each of these connected components, a *source* vertex, corresponding to the leftmost aligned short read of the component, and a *target* vertex, corresponding to the rightmost aligned short read, are defined. Dijkstra’s algorithm [19] is then used, in order to find the shortest path of the graph allowing to link the source and the target. This path, through the short reads associated to the traversed vertices, dictates a correction for the region of the long read which is covered by the alignments contained in this connected component.

To correct long reads regions that were not covered by any short read during the alignment step, CoLoRMap makes use of the information carried by the paired reads that can be sequenced from second generation sequencing technologies. *One-End Anchors (OEA)* are thus defined, in order to correct these uncovered regions. These OEA are short reads than could not be aligned, but whose mate could be aligned to a region flanking the uncovered region to correct. The set of OEA associated to a given uncovered region of a long read can thus be defined by observing the alignments located in the flanking, corrected regions.

For this second correction step, short reads are aligned to the corrected long reads, obtained at the end of the previous step, once again with the help of BWA. The short reads can indeed easily be aligned to the corrected regions of the long reads, due to the low divergence between their sequences and the corrected regions of the long reads. For each region of a given long read that could not be corrected during the previous step, CoLoRMap defines a set of OEA, by observing the alignments located in the flanking regions. The reads composing this set of OEA are then assembled, with the help of Minia [13], in order to obtain a set of contigs, which is then, in turn, used to correct the uncovered region.

CoLoRMap thus produces untrimmed corrected long reads. However, the bases of the long reads that could not be corrected are reported in a different case from the corrected bases. Thus, the corrected long reads can easily be trimmed or split after correction, in order to get rid of their uncorrected regions.

#### HECIL (2018)

HECIL computes alignments with the help of BWA-MEM. From these alignments, erroneous positions, denoting disagreements (insertions, deletions or substitutions) between a given long read and the short reads that align to it are marked. For each of this positions, the set of aligned short reads is processed, and two correction steps are applied.

First, in case of strong consensus (*i.e.* at least 90%) of the short reads at a given position, the nucleotide of the long read is replaced by the consensus nucleotide from the short reads. This first step, inspired by other methods relying on majority votes, allows for a quick first correction step. Moreover, it also allows to avoid spurious correction at certain positions, performed from high-frequency, but low-quality short reads. The corrections that are realized during this first step also allow to greatly reduce the number of positions to process during the next step.

In a second time, for the remaining erroneous positions, a score is associated to each aligned short read, by combining the quality of the short read, obtained through its Phred score, to the similarity score of the alignment between the short read and the long read. The main motivation behind this combined score is that the incorporation of short reads quality into the problem of long-read correction had not been tackled by any state-of-the-art method at that time. For each erroneous position, the base from the short read which maximizes the combined score is chosen as the correction of the long read’s base. In case of a tie between multiple short reads, the base from the short read having the highest quality is chosen as the correction.

Moreover, HECIL also defines an iterative learning methodology. More precisely, the two previously described correction steps are applied iteratively. At each iteration, the previously defined combined scores are used in order to define a subset of high-confidence positions to correct. Only the positions contained in this subset are thus corrected during this given iteration. These positions are then fixed and marked, and are thus no longer modified during subsequent iterations. By only correcting the best subset of position for each iteration, HECIL aims to perform better correction during the following iterations, since these iterations will benefit from the higher quality of the long reads, increased by the previous iterations. Thus, at each iteration, the short reads are realigned to the updated long reads, and the two correction steps are performed on a new subset of high-confidence positions. The iterations are stopped when the number of unique *k*-mers of the long reads varies less than a given threshold between two consecutive iterations, or once a user-defined maximum number of iterations is reached.

HECIL thus produces untrimmed corrected long reads. Indeed, local corrections are performed on the long reads, on positions denoting disagreements with the aligned short reads. Thus, the uncovered extremities of the long reads cannot be modified or corrected. Moreover, the covered bases of the long read that could not be corrected are not reported in a different case from the corrected bases. As a result, the corrected long reads cannot further be split after correction.

#### 2.1.2 Contigs alignement

Given their length, short reads can be difficult to align to repeated regions, or to extremely noisy regions of the long reads. The contigs alignment approach aims to address this issue by first assembling the short reads. Indeed, the contigs obtained after assembling the short reads are much longer than the original short reads. As a result, they can cover the repeated or highly erroneous regions of the long reads much more efficiently, by using the context of the adjacent regions during alignment. In the same fashion as the short reads alignment strategy described in Section 2.1.1, the contigs aligned with the long reads can then be used to compute high quality consensus sequences, and thus correct the long reads they are associated to. Once again, the different methods adopting this strategy vary by the alignment methods they use, and by algorithmic choices made during consensus computation.

ECTools requires a set of short reads assembled into unitigs as a basis for its error correction process. This means that the short reads assembly has to be performed before launching ECTools, and that it is thus not automatically generated during the error correction process. The assembly can thus be produced by any tool, although the authors recommend using Celera [60], as it guarantees that each short read is included in a unitig, by creating unitigs composed of a single short read, if necessary. This allows ECTools to ensure all the information contained in the short reads will be transmitted to the set of unitigs, and that no information will be lost during the assembly process.

The unitigs are aligned to the long reads with the help of Nucmer, which is available in the MUMmer [40] suite of tools. The subset of unitigs that covers the best each long read is then chosen and used for its correction. This optimal subset is computed with the help of an algorithm based on the longest increasing subsequence problem, which is also available in the MUMmer suite of tools. Once computed, this subset of unitigs is used to correct the long read. The bases of the long read which are different from those of the selected unitigs are identified, and the long read is corrected according to the bases of the unitigs.

ECTools thus produces split corrected long reads. Uncovered extremities of the long reads are first removed from the sequences of the corrected long reads. Then, internal, uncovered regions, are considered as uncorrected, and are labeled with an error rate of 15%. In the same fashion, covered regions are considered as corrected, and are labeled with an error rate of 1%. The average error rate of each corrected long read is then computed on its whole length, according to these labels, and is then compared to a minimal quality threshold. If the average quality of a given long read is below that threshold, the read is then split, by removing the longest uncorrected region from its sequence. This process is then recursively applied to each obtained fragment, until they are all above the minimum quality threshold.

#### HALC (2017)

In the same way as ECTools, HALC also requires a set of assembled short reads as a basis for its error correction process. Once again, assembly of the short reads thus has to be performed before launching HALC, and is not automatically generated during the error correction process. Unlike ECTools, HALC however makes use of the contigs generated by the assembly, instead of only using the unitigs. The long reads are thus aligned to the contigs, with the help of BLASR [12], to initiate the error correction process.

An undirected graph is then built from the alignments between the long reads and the short reads. This allows to represent the alternative alignments, in particular in repeated regions, by different paths. Vertices of the graph are thus defined from the regions of the contigs to which a long read aligns. An edge is present between two vertices if at least a pair of adjacent regions of a given long read aligned to the two regions of the contigs corresponding to these vertices.

This graph is also weighted, and the weight of each edge is computed according to two rules. First, the weight of an edge (*u, v*) is set to 0 if the regions corresponding to vertices *u* and *v* are adjacent in one of the contigs. Contrariwise, for a given edge (*u, v*) for which the regions corresponding to vertices *u* and *v* are not adjacent in any of the contigs, the weight is set to *max C*_0_ *C,* 0. *C*_0_ represents the long read coverage on the contigs, whereas *C* represents the number of adjacent regions of the long reads that align to the regions corresponding to vertices *u* and *v*.

Once the graph is built, each long read is processed and corrected independently. To this aim, paths representing alternative alignments are searched through the graph. The path displaying the lowest cumulative weight is then computed via a dynamic programming algorithm, and the regions of the contigs corresponding to the traversed vertices are used to correct the long read.

However, with this approach, adjacent regions of the long reads that are corrected with the help of non-adjacent contigs regions have a high chance of being repeated regions. Thus, they also have a high chance of having been corrected with similar repeats rather than the true genome regions they actually belong to. Such regions are recorded, and their correction is then polished with the help of LoRDEC, another hybrid correction tool presented in Section 2.1.3.

HALC thus produces untrimmed corrected long reads. However, the last polishing step, being based on LoRDEC, allows to report the corrected bases in a different case from the uncorrected bases. As a result, the corrected long reads can easily be trimmed or split after correction, in order to get rid of their uncorrected regions. In particular, untrimmed, trimmed, and split corrected long reads are all reported by HALC.

Unlike previously presented methods, MiRCA starts by applying a filtering step to the short reads. Indeed, although they display a much higher quality than the long reads, short reads also contain sequencing errors that can lead to imprecisions during the assembly step and during the alignment step of the long reads to the obtained contigs. Thus, in MiRCA, short reads containing too many errors are removed, and are not considered during the following steps. This filtering step relies on a frequency analysis of the short reads’ *k*-mers. Short reads containing too many weak *k*-mers (*i.e. k*-mers appearing less frequently than a given threshold) are thus filtered out.

High-quality short reads are then assembled with the help of SPAdes [5], an assembler based on a de Bruijn graph, that allows to ensure that all the information contained in the short reads is preserved in the obtained set of contigs. These contigs are then aligned to the long reads with the help of BLAST+ [11].

The long reads can then be corrected from the obtained alignments. In the case a unique contig aligns to a given region of a long read, this region is simply replaced by the sequence of the aligned contig. In the case where multiple contigs are aligned to a same given region of a long reads, these alignments define a multiple sequence alignment, and a majority vote is used to correct the region of the long read. If the majority vote does not allow to define a consensus nucleotide at a given position, the long read nucleotide at this position is kept as the correction.

MiRCA thus produces trimmed corrected long reads. Indeed, extremities of the long reads to which no contig aligned cannot be processed by the algorithm, and are thus removed from the sequences of the corrected long reads. However, the internal regions of the long reads that were not covered by a single contig, and that could not be corrected, are not reported in a different case from the corrected bases. As a result, corrected long reads cannot further be split after correction.

#### 2.1.3 De Bruijn graphs

Another alternative to the alignment of short reads to the long reads is the direct use of a de Bruijn graph, built from the short reads *k*-mers. This approach aims to avoid the explicit step of short reads assembly altogether, contrary to methods mentioned in Section 2.1.2, and instead directly use the graph to correct the long reads. The graph is first built from the *k*-mers of the short reads. The long reads can then be anchored to the graph according to their *k*-mers. Finally, the graph can be traversed, in order to find paths, and link anchored regions of the long reads together, and thus correct erroneous, unanchored, regions. Methods adopting this approach vary by the way they represent the graph, but also by the way they anchor the long reads to the graph, and by the way they correct unanchored regions.

#### LoRDEC (2014)

LoRDEC starts by building a de Bruijn graph from the solid *k*-mers (*i.e. k*-mers that appear more frequently than a given threshold) of the short reads. This filtering step of the weak *k*-mers allows to avoid the introduction of errors during the correction of the long reads, as it limits the presence of tips and bubbles in the graph. The solid *k*-mers of the long reads are then used as anchor points on the graph. The graph is then traversed, from and towards the vertices corresponding to these anchors, in order to find paths allowing to corrected regions of the long reads that are only composed of weak *k*-mers. Thus, if a given long read does not contained any solid *k*-mer, it cannot be corrected.

Given the high error rates of the long reads, a small *k*-mer size has to be chosen for the graph construction, in order to allow the correction of as much long reads as possible. Thus, it is recommended to build the graph for values of *k* between 19 and 23. Two different correction approaches are then applied, according to the localization of the region to correct. Internal regions and extremities of the long reads are indeed corrected differently.

In the first case, solid *k*-mers located before and after the region to correct are used as anchor points on the graph. The solid *k*-mers located on the left of the region to correct define *sources*, whereas the solid *k*-mers located on the left define *targets*. The graph is thus traversed, in order to find a path between a source and a target *k*-mer. Once found, such a path dictates a correction for the erroneous region of the long read. In order to find an optimal path, at each branching path, the edge minimizing the edit distance to the erroneous region of the long read is chosen. However, a limit on the maximal number of branching paths that can be explored is set, in order to avoid extensive and costly traversals of the graph. Several pairs of source and target *k*-mers are thus considered, in order to limit the cases where no path can be found to correct a weak *k*-mers region. As a result, the graph is traversed multiple times for the correction of a given region. Several of these traversals can thus find a path between two anchor *k*-mers, and dictate a correction for the erroneous region of the long read.

In this case, the path dictating the sequence minimizing the edit distance to the erroneous region of the long read is chosen as the correction.

In the second case, one of the source or target *k*-mer is missing, and the search thus cannot be stopped when a path between two *k*-mers is found. In this case, the graph is only traversed from a solid *k*-mer close to the erroneous region. The traversal of the graph stops when the length of the erroneous extremity of the long read is reached, when the edit distance between the current path and the erroneous region is too high, or when two edges can be followed out of the current vertex. Once the graph traversal is over, the obtained sequence is realigned to the extremity of the long read. This allows to determine the prefix (resp. the suffix) of the graph sequence that aligns the best to the extremity of the long read. The aligned prefix (resp. suffix) of the long read extremity is thus corrected with the prefix (resp. suffix) of the graph sequence. This alignment step allows to ensure divergent sequences, caused by inappropriate traversals of the graph, are not used as correction.

Two correction passes are thus applied for each long read, first from the left to the right, and then from the right to the left. Indeed, the long reads are corrected on the fly by the graph traversals. These corrections thus allow to generate new solid *k*-mers that can be used as anchor points by the second correction pass. Moreover, repeats and sequencing errors that are present in the short reads can lead to the traversal of different subgraphs, according to the extremity that is chosen as the starting point of the path search.

LoRDEC thus produces untrimmed corrected long reads. However, bases corresponding to solid *k*-mers are reported in a different case from bases corresponding to weak *k*-mers. As a result, the corrected long reads can easily be trimmed or split after correction, in order to get rid of their uncorrected regions. Two additional tools allowing to perform these operations are provided along with LoRDEC.

#### Jabba (2016)

Unlike previously presented methods, Jabba starts by correcting the short reads, in order to limit the impact of their sequencing errors on the quality of the de Bruijn graph, and thus on the quality of the correction. To this aim, the authors recommend using Karect [1], although any correction tool can be used. The de Bruijn graph is then built from the *k*-mers of the corrected short reads. To further reduce the impact of the errors contained in the graph on the correction of the long reads, another correction procedure is applied to the graph itself. More precisely, the tips are eliminated, and the bubbles are removed from the graph, by selecting, for each of them, the highest supported path. The construction and correction of the graph is performed with the help of Brownie^2^.

Moreover, unlike LoRDEC, Jabba does not rely on the study of *k*-mers that appear both in the graph and in the long reads to find anchor points, but rather uses Maximal Exact Matches (MEM) between the long reads and the vertices of the graph. Thereby, since the graph also displays a high quality, due to the two correction processes, large values of *k* can be used for its construction. Usually, the graph is built with *k* = 75. Using such values of *k* allows to reduce the complexity of the graph, by resolving short repeats, of size smaller than *k*, directly inside the graph, which, in turns, allows to facilitate the alignment of the long reads to the graph, during the following step.

A seed-and-extend strategy is applied to align the long reads to the graph. First, the MEMs, which represent the seeds, are computed with the help of essaMEM [69]. This method is based on an enhanced suffix array, built for the sequence obtained by concatenating all the vertices of the graph, as well as their reverse-complements. Once the seeds of a given long read are computed, they are placed of the graph. To this aim, several passes are performed on the long read. For each iteration, all the regions of the long reads that have not yet been aligned are considered. For each of these regions, the longest seeds are used, in order to determine the vertices to which this region can be aligned. The quality of the alignments between each vertex and the long read region are then verified. Alignments that are fully included in longer alignments (called *containments*), and alignments covering a fraction of the vertex that is below a given threshold are then filtered, and are thus not considered during the following step.

Once the alignments between the regions of the long read and the vertices of the graph are computed for the current iteration, the alignments between the vertices are chained, by following the paths of the graph. Each alignment on a vertex is thus extended, by considering all the possible paths of the graph starting from this vertex. If the vertex only has one outgoing edge, the alignment is extended along that edge. If multiple outgoing edges are available from this vertex, the length of the target vertex of each edge is studied. The extension taking place between two regions of the long read, a maximal distance between the two edges can be defined, and edges that are too long are thus not considered. In the case where no outgoing edge can be followed from a given vertex, the corresponding region of the long read is reprocessed and realigned during the next iteration.

Once all the iterations have been performed, several alignments usually remain possible for each long reads. The alignment covering the largest proportion of the long read sequence is thus selected, and used for the correction. For the correction of the extremities of the long read, the alignment is extended along unique paths of the graph. If the long read extends outside of these unique paths, its extremities are trimmed. Indeed, correcting these extremities is extremely time consuming, since they need to be aligned to all the possible paths, as no seed can be used to guide the alignment. This problem is thus not tackled by Jabba, as the gain in quality is too weak compared to the computational cost.

Jabba thus produces split corrected long reads. Indeed, as mentioned in the previous paragraph, extremities of the reads are not corrected outside of the longest unique path of the graph to which they align. Moreover, in cases where, even after multiple iterations, no edge can be followed to extend the alignment of a given vertex, a direct path between the first and the last seed of the long read does not exist. In this case, Jabba splits the long read, and independently corrects and outputs the different regions of this read that could be aligned and chained along the graph.

#### FMLRC (2018)

FMLRC relies on a principle that is similar to that of LoRDEC. Indeed, it also builds a de Bruijn graph from the *k*-mers of the shorts reads, and uses the solid *k*-mers of the long reads as anchor points on this graph, in order to find paths allowing to correct the regions of the long reads only composed of weak *k*-mers. One of the main differences between FMLRC and LoRDEC is that the de Bruijn graph is implicitly represented in FMLRC, with the help of an FM-index built from the sequence obtained by concatenating the set of short reads. As a result, the edges between the different vertices are found by querying the FM-index. The size of *k* thus does not have to be known at construction time, but is rather directly chosen at execution time. This way, the FM-index allows to represent all the de Bruijn graphs, and not simply a single, fixed order, graph. Such a graph is called a *variable-order* de Bruijn graph. However, unlike LoRDEC, the weak *k*-mers of the long reads are not filtered, and all the *k*-mers of the short reads are thus present in the graph.

The correction of the long reads is then realized in the same way as LoRDEC. For regions composed of weak *k*-mers bordered by two regions composed of solid *k*-mers, source and target *k*-mers are defined on each bordered regions, and used as anchor points on the graph. A path is then searched through the graph, from the source toward the target, and from the target toward the source. Indeed, the search can traverse different subgraphs, according to its starting point. As in LoRDEC, if multiple paths are found, the one minimizing the edit distance to the erroneous region of the long read is chosen as the correction. For erroneous regions located at the extremities of the long reads, the solid *k*-mers that is the closest to the erroneous region is chosen as the anchor point. The graph is thus traversed, and the traversal stops when the length of the path reaches the length of the erroneous region to correct. Several paths can thus be obtained, and the one minimizing the edit distance to the erroneous region of the long read is, once again, chosen as the correction.

However, unlike LoRDEC, FMLRC performs two passes of correction for each long read. A first pass is performed with a small value of *k* (*k* = 20 by default), and a second pass in then performed, with a larger value of *k* (*k* = 59 by default). The first pass usually manages to correct most errors, and the second pass allows to polish the correction, especially in repeated regions, which are easier to resolve with the help of a higher order de Bruijn graph. Moreover, unlike LoRDEC, the solidity threshold of the long reads’ *k*-mers is dynamically adjusted for each long read, according to the chosen *k*-mer size and to the *k*-mers frequencies of this long read. The limit on the maximal number of branching paths that can be explored is also set according to the *k*-mer size, in order to reduce the number of branches exploration during the first pass. Indeed, the traversal of the first graph can be extremely costly, due to the important number of branching paths in repeated regions, when using a small *k*-mer size. These regions are rather resolved during the second correction pass, where a greater number of branches exploration is allowed, and where the *k*-mer size is higher.

FMLRC thus produces untrimmed corrected long read. However, unlike LoRDEC, bases corresponding to solid *k*-mers are not reported in a different case from bases corresponding to weak *k*-mers. As a result, the corrected long reads cannot further be trimmed or split after correction.

#### ParLECH (2019)

Unlike other de Bruijn graph based methods, ParLECH relies on two distinct correction steps, depending on the type of errors. A first step is thus dedicated to the correction of indels, and another step focuses on the correction of substitutions, which can be introduced by the short reads during the correction of indels.

For the correction of indels, ParLECH first builds the de Bruijn graph from the *k*-mers of the short reads. The graph is stored in a key-value hashmap, with the help of HazelCast^3^, where the *k*-mers define the keys, and where predecessors and successors of these *k*-mers define the values. Moreover, a number of occurrences is associated to each key. In the same fashion as in previously presented methods, regions solely composed of weak *k*-mers are corrected by traversing the graph, using flanking, solid *k*-mers as anchor points. If multiple paths allow to link together a given pair of anchors, the path maximizing the minimum *k*-mer coverage between two vertices is chosen as the correction.

For the correction of substitutions, ParLECH first divides the long reads into shorter fragments, approximatively of the size of the short reads. Indeed, *k*-mers in smaller subregions usually have similar abundances. This allows ParLECH to divide the long reads into sequences of high and low coverage fragments. If a fragment belongs to a low coverage region of the genome, most of its *k*-mers are expected to have a low coverage. On the contrary, most of its *k*-mers are expected to have a high coverage. This methodology allows ParLECH to better distinguish low coverage genomic *k*-mers, and high coverage erroneous *k*-mers. Once the long reads have been fragmented, the Pearson’s skew coefficient of the *k*-mer coverage of each fragment is computed, and used as a threshold to classify fragments as correct or erroneous. If the coefficient of a fragment falls within a given interval, the fragment is considered as correct. Otherwise, it is considered as erroneous. Moreover, fragments with mostly low-coverage *k*-mers are ignored.

After classification, the erroneous fragments are divided into subsets of high and low coverage. To this aim, if the median *k*-mer coverage of a fragment is greater than the median coverage of the whole set of *k*-mers, the fragment is considered as high coverage. Otherwise, it is considered as low coverage. To correct substitutions, ParLECH makes use of a majority vote algorithm similar to that of Quake [33]. However, unlike Quake, ParLECH makes use of different thresholds for high and low coverage regions, in order to improve accuracy. For each erroneous base detected during the previous step, ParLECH attempts to replace it with all the other possible nucleotides, and computes the coverage of all the *k*-mers with this new base. The erroneous base is thus replaced by the base such that all *k*-mers containing this base exceed or equal the threshold for this region. ParLECH thus produces untrimmed corrected long reads. Moreover, bases associated to subgraphs that could be trained are not reported in a different case than the bases to which no short read aligned. As a result, the corrected long reads cannot further be trimmed or split after correction.

#### 2.1.4 Hidden Markov Models

Hidden Markov Models, used for short read error correction, were also adopted for the error correction of long reads. To this aim, models are first initialized in order to represent the original long reads. A subset of short reads is then assigned to each long read, by alignment. Each subset of short reads can then be used to train the model it is associated to. Finally, the trained models can be used to compute consensus sequences, and thus correct the long reads they represent.

#### Hercules (2018)

Like LSC and LSCplus, Hercules starts by compressing homopolymers regions of the long reads and of the short reads. The compressed short reads are then filtered, and those that are shorter than a given threshold are removed. Such short reads could indeed ambiguously align to the long reads. The remaining short reads are then aligned to the long reads. Bowtie2 is used by default, although any aligner can be used, as long as it outputs alignments in SAM format.

After computing the alignment, Hercules decompresses both the long reads and the short reads, and recomputes the beginning positions of the alignments, according to the uncompressed sequences. However, unlike other methods based on the alignment of short reads to the long reads, Hercules then only uses the beginning positions of the alignments, and does not make use of any other information reported by the aligner. Thus, the choice of the bases used to correct the long reads is not performed by the aligner.

Each long read is then represented with the help of a profile Hidden Markov Model (pHMM). These pHMMs have three different types of states, allowing to represent deletions, insertions, or matches / mismatches. The aim of Hercules is to make use of the pHMM to produce the consensus sequence of the long read, which is defined as the most probable sequence, among all the sequences that can be produced by the model. However, the consensus that can be produced by a classical pHMM is only based on match states. As a result, in order to take into account insertion and deletion states during consensus computation, the pHMM are modified in Hercules. Loops on insertion states are removed, and replaced by multiple, distinct, insertion states. Deletion states are also removed, and replaced by deletion transitions. Once the model of a given long read is built, emission and transmission probabilities of each state are initialized according to the error profile of the sequencing technology.

The model is then updated according to the information contained in the short reads. To this aim, each alignment of a short read to a long read is considered independently. First, the subgraph corresponding to the region of the long read which is covered by this short read is extracted. Then, this subgraph is trained, with the help of the information contained in the selected short read, with the help of the Forward-Backward algorithm [8]. Emission and transmission probabilities of each vertex are thus updated by the algorithm, independently and exclusively in each subgraph, using the probabilities of the initial model. In the case a vertex is considered and updated in multiple subgraphs, its final probabilities, in the trained model, are defined by computing the average probability of each subgraph it appears in. Uncovered regions of the long reads cannot yield subgraphs. Emission and transmission probabilities of the initial models are thus kept, for the corresponding vertices.

Once all the alignments have been considered, and once all the subgraphs have been trained, the model is updated according to these subgraphs. Viterbi algorithm [68] is then used, in order to decode the consensus sequences of the updated model. This algorithm considers the pHMM associated to each long read, and searches for a path, from the initial state to the final state, which produces the highest emission and transmission probabilities, via a dynamic programming approach.

Hercules thus produces untrimmed corrected long reads. Indeed, the consensus sequence of a long read is extracted from the pHMM by searching for a path from the initial to the final state, representing respectively the first and the last base of this read. The entire length of the long reads is thus preserved after correction. Moreover, bases associated to subgraphs that could be trained are not reported in a different case than the bases to which no short read aligned. As a result, the corrected long reads cannot further be trimmed or split after correction.

#### 2.1.5 Combination of strategies

Other methods combine different of the aforementioned strategies, in order to balance their advantages and drawbacks. For instance, NaS combines a first step of short reads alignment to a second step of short reads recruitment and assembly, in order to correct the long reads. HG-CoLoR also relies on a first step relying on short reads alignment, but then makes use of a variable order de Bruijn graph, in order to correct regions of the long reads that were not covered by the original alignments. We describe these two tools below.

#### NaS (2015)

Unlike previously presented methods, which either perform local correction on specific long reads regions, or compute consensus sequences from short reads alignments, NaS generates corrected long reads by assembling subsets of related short reads. These sequences, called synthetic long reads, are computed as follows.

First, the alignments between the short reads and the long reads are computed either with the help of BLAT [34] or LAST [35], according to the chosen correction mode (fast or sensitive, respectively). The short reads thus aligned to the long reads define seeds. For each long read, a seed-and-extend strategy is then applied. The seeds are compared to the set of short reads, in order to recruit new similar short reads, that allow to cover the regions of the long reads that were not covered by the initial alignments. This recruitment step is performed with the help of Commet [52], which considers two short reads are similar if they share a sufficient number of non-overlapping *k*-mers.

After recruiting new short reads, the subset of short reads formed by these new reads and by the seeds is assembled with the help of Newbler [55]. A unique contig is usually built, although in repeated regions, a few short reads originating from the wrong copy of the repeat can be erroneously recruited, and thus yield additional contigs that should not be associated to the original long read. To address this issue, and produce a single contig for each long read, the contig graph is explicitly built. This graph is a directed and weighted graph, for which the contigs defined the vertices. An edge is present between two vertices if the associated contigs overlap. The weights are however associated to the vertices, and not to the edges. Thus, the weight of each vertex is defined as the coverage of the contig by the seed short reads. Once the graph is built, Floyd-Warshall algorithm [23, 72] is used in order to extract the shortest path. The final contig is thus obtained by combining the contigs associated to the vertices traversed by this path.

Finally, the short reads are realigned to the obtained contig, in order to verify its consistency. The contig is then defined as the correction of the original long read, if it is sufficiently covered by short reads.

Unlike previous methods, that usually either trim or split the corrected long reads, NaS rather tends to extend the reads, and thus produce corrected long reads that are longer than the original reads they are associated to. This is caused by the fact that the recruitment process often includes short reads that are located outside the extremities of the original long read to the subset of short reads used for the assembly. Such a behavior can thus assemble corrected sequences that span outside of the original long reads.

#### HG-CoLoR (2018)

HG-CoLoR combines together a short read alignment and a de Bruijn graph approach into a seed-and-extend strategy. As a first step, it starts by correcting the short reads, in order to get rid of as much sequencing errors as possible. Indeed, since these short reads will be aligned to the long reads, and will also be used to build the graph, they need to reach a high level of quality to avoid the introduction of errors into the corrected long reads. This short-read error correction step is performed using QuorUM [53].

HG-CoLoR then aligns the corrected short reads to the long reads with the help of BLASR. A set of seeds is thus defined for each long read, and covered regions of the long reads can directly be corrected with help of these seeds. After the alignment step, a variable order de Bruijn graph is built from the solid *k*-mers of the corrected short reads. Unlike FMLRC, this graph is built with the help of PgSA [39]. Moreover, HG-CoLoR allows to explore every order of the graph, between a minimum order *k* and a maximum order *K*, instead of limiting the graph explorations to two different orders. To correct uncovered regions of the long reads, the prefixes and suffixes of the seeds that aligned to covered regions are used as sources and targets on the graph, which is traversed to perform correction.

The graph is always explored starting from its highest order. The order of the graph is thus only locally decreased if no edge can be followed out of the current node, or if all its edges, for the current order, have already been explored and did not allow to link the source and the target. At branching paths, for a given order *k*, HG-CoLoR performs a greedy selection and explores the edge leading to the *k*-mer having the highest number of occurrences. Thus, when a path between two *k*-mers is found, it is considered as optimal, due to that greedy selection strategy, and due to the fact that the order of the graph is only locally decreased. As a result, it is thus directly chosen as the correction of the uncovered region of the long read. As in other methods, a maximal number of branches explorations is also set, in order to avoid resource consuming traversals of the graphs. Moreover, if a given pair of seeds cannot be link after reaching that limit, HG-CoLoR can attempt to skip a certain number of seeds, and thus try link together the same source to different targets. A maximal number of skips is also set, and if no source-target pair could be linked, the uncovered region of the long read remains uncorrected.

Finally, as seeds do not always cover the extremities of the long reads, HG-CoLoR keeps on traversing the graph after linking together all the seeds, in order to correct the extremities of the long reads. The graph is thus traversed, until the whole length of the processed long read has been corrected, or until a branching path is reached. If the extremities of the original long read could not be reached, they are reported unmodified in the corrected long read.

HG-CoLoR thus produces untrimmed corrected long reads. However, the corrected bases, whether they come from seeds or from graph traversals, are reported in a different case from the uncorrected bases. As a result, the corrected long reads can easily be trimmed or split after correction, in order to get rid of their uncorrected regions. In particular, untrimmed, trimmed, and split corrected long reads are all reported by HG-CoLoR.

### 2.2 Self-correction

Self-correction aims to avoid the use of short reads data altogether, and to correct long reads solely based on the information contained in their sequences. Third generation sequencing technologies indeed evolve fast, and now allow the sequencing of long reads reaching error rates of 10-12%. As a result, correction is still required to properly deal with errors, but self-correction has recently undergone important developments. Two different self-correction approaches thus exist:

1. Multiple sequence alignment. This approach is similar to the hybrid approaches relying on short reads or contigs alignments, described in Section 2.1.1 and in Section 2.1.2. Indeed, once aligned against each other, the long reads can be corrected by computing a consensus sequence for each of them, in the same fashion as in the hybrid approaches. PBDAGCon (the correction module used in the HGAP assembler) [15], PBcR-BLASR [36], Sprai^4^ [26], PBcR-MHAP [9], FalconSense (the correction module used in the assembler Falcon) [14], Sparc [74], the correction module used in the assembler Canu [38], MECAT [73] and FLAS [7] rely on this approach.
2. Use of de Bruijn graphs. In a similar manner as the hybrid correction approach using de Bruijn graphs, described in Section 2.1.3, once the graph is built, it can be used to anchor the reads. The graph can then be traversed, in order to find paths allowing to link together anchored regions of the long reads, and thus correct unanchored, erroneous regions. LoRMA [64] and Daccord [67] are based on this approach.

As for hybrid correction, other methods, such as CONSENT [58], combine these two approaches, in order to counterbalance their advantages and their drawbacks. We describe each approach more into details, and list the related tools, in the following subsections.

#### 2.2.1 Multiple sequence alignment

This approach is highly similar to the short reads alignment approach for hybrid correction, described in Section 2.1.1, and to the contigs alignment approach described in Section 2.1.2. It is thus composed of a first step of overlaps computation between the long reads, and of a second step of consensus computation from the overlaps. The overlaps computation can be performed either via a mapping strategy, which only provides the positions of the similar regions of the long reads, or via alignment, which provides the positions of the similar regions, as well as their actual base to base correspondence in terms of matches, mismatches, insertions and deletions. For the consensus computation step, a DAG is usually built in order to summarize the alignments, and extract a consensus sequence. Methods adopting this strategy thus vary by the overlapping strategy they rely on, but also by algorithmic choices made during the consensus computation step.

#### PBDAGCon (2013)

PBDAGCon is a consensus module, included in the assembler HGAP. First, the alignments between the reads are computed with the help of BLASR. For each read, a DAG is then built, from the set of similar reads determined during the previous step. The vertices of this graph are labeled according to the bases of the reads, with one base per vertex. An edge is present between two vertices if there exists a read containing the two bases corresponding to these vertices consecutively (*i.e.* there is an edge between *u* and *v* if *uv* is a factor of any read). This way, every read can be represented by a unique path in this graph. Each vertex is thus supported by a given set of reads, and each edge is also supported by a given set of reads. The graph can thus be weighted, by defining the weight of an edge as the number of reads that support it, *i.e.* the number of reads that contain the factor dictated by the two vertices.

The graph is built iteratively. First, it is initialized with the sequence of the read to be corrected. All the edges are thus initialized with weight 1. The graph is then updated, by adding the other reads sequentially, with the help of the information contained in the alignments. This way, if an alignment corresponds to vertices and edges that are already present in the graph, the weight of the corresponding edges are simply incremented. Otherwise, new vertices and edges are created if necessary, and the weight of the edges are set to 1.

Once all the reads have been added to the graph, the consensus sequence of the original read can be computed. To this aim, a score is assigned to each vertex, according to the weight of its outgoing edges. The path that maximizes the cumulative score of the traversed vertices is then searched through the graph, with the help of a dynamic programming algorithm. The starting point of this path is defined as the vertex corresponding the first base of the original read, and the ending point is defined as the vertex corresponding to its last base.

PBDAGCon thus produces untrimmed corrected long reads. Indeed, the consensus sequence of a read is computed by searching for a path in the graph between the vertices corresponding to the first and last base of this read. The entire length of the original reads is thus preserved by the correction process. Moreover, the bases of the original reads that could not be corrected, due to a lack of coverage, are not reported in a different case from the corrected bases. As a result, the corrected long reads cannot further be trimmed or split after correction.

#### PBcR-BLASR (2013)

PBcR-BLASR is a modified version of the PBcR hybrid correction tool, described in Section 2.1.1, that adapts to the problem of self-correction. Thus, the alignment between the reads are first computed with the help of BLASR. The obtained alignments are then used to correct the reads, with the help of the PBcR correction algorithm.

Since it relies on the PBcR algorithm, PBcR-BLASR thus produces split corrected long reads, for the reasons mentioned in the description of the PBcR algorithm, in Section 2.1.1.

#### Sprai (2014)

Sprai starts by computing the alignments between the reads with the help of BLAST+. The set of similar reads that align to a given read can thus be used to define a multiple sequence alignment. This multiple sequence alignment is then polished, with the help of ReAligner [3], in order to maximize its score by modifying its layout, while conserving its global structure. The main aim is to gather similar bases onto identical columns of the matrix representing the multiple sequence alignment. Indeed, local choices are performed when aligning two reads together, in particular in regions denoting insertions and deletions. Such errors being frequent in long reads, the combination of several alignments into a single multiple sequence alignment does not necessarily ensures the coherence of the choices made in each independent alignment. The polishing thus aims to restore this coherence, by locally modifying the multiple sequence alignment, in order to gather enough support at each position.

To this aim, all the sequences included in the multiple sequence alignment are considered independently. Each of these sequences is then realigned to the multiple sequence alignment, from which the sequence being processed has been excluded, via a dynamic programming algorithm. The multiple sequence alignment is thus updated, according to the result of the alignment computation. Other sequences are then considered iteratively, and the process is thus repeated, until the score of the multiple sequence alignment no longer increases. The final multiple sequence alignment obtained is then used to compute the consensus sequence of the original read, with a majority vote at each position.

Sprai thus produces untrimmed corrected long reads. Indeed, the multiple sequence alignments are always built at the scale of the whole reads. Therefore, the consensus computed via majority vote also contain the uncovered bases of the reads. Moreover, the bases coming from the original reads, and that could not be corrected due to a lack of coverage, are not reported in a different case from the corrected bases. As a result, the corrected long reads cannot further be trimmed or split after correction.

#### PBcR-MHAP (2015)

PBcR-MHAP is an upgraded version of PBcR-BLASR. In this version, the computation of the alignments between the reads, previously performed with BLASR, is replaced by a mapping strategy called MHAP (MinHash Alignment Process). In this approach, the *k*-mers of the reads are extracted and converted into integer fingerprints with the help of hash functions. A sketch, whose size is defined by the number of hash functions, is thus built for each read. For each of these hash functions, the *k*-mer of the read that generates the smallest value is chosen as a part of the sketch. Such a *k*-mer is called a *min-mer*. The sketch of a read is thus defined a the ordered set of the integer fingerprints of its min-mers. The fraction of entries shared by two reads sketches can thus be used to estimate their similarity. Thus, if two given sketches share a sufficient similarity, the min-mers they both share are localized on the corresponding reads, and the median difference between their positions is computed, in order to determine the positions of the overlap between the two reads. A second iteration of this process is then applied, only on overlapping regions, in order to obtain a better estimate of the similarity between these sequences.

Once the overlaps between all the reads have been computed, the overlaps can be used to correct the reads. However, the algorithm used both in PBcR and PBcR-BLASR is replaced by two new consensus modules: FalconSense, which is faster but less sensitive, and PBDAGCon, which is slower but more accurate. These two algorithms are not detailed here, but are described in their respective presentations. Moreover, these two algorithms are not applied one after the other, and the choice as to which one to use is left to the user.

PBcR-MHAP thus produces split corrected long reads if FalconSense is used, and untrimmed corrected long reads, for which uncorrected bases are not reported in a different case from the corrected bases, if PBDAGCon is used. We do not further detail the specificities of these corrected reads, since they are explained in the respective description of FalconSense and PBDAGCon.

#### FalconSense (2016)

FalconSense is a correction module, which is included in the assembler FALCON. First, the alignments between the reads are computed with the help of DALIGNER [61]. The set of similar reads which align to a given read is then independently realigned to this read, without allowing mismatches, and thus encoding every edit operation with insertions and deletions. For each alignment position between the original read *A* and one of its similar reads *B*, a tag is generated. Each of these tags is composed of the four following fields:

1. The position of read *A*;
2. The offset from this position on the read *A*. This value differs from 0 when an insertion is present on read *B* at this alignment position, and thus indicates the size of the offset between the position on read *A* and the position on the alignment;
3. The base of read *A* at this position;
4. The base of read *B* at this position.

Once all the tags have been defined, for every position of each alignment between the original read and its similar reads, an alignment graph can be built. This graph is a weighted DAG, whose vertices are defined by the tags. An edge is created between two vertices if their corresponding tags are consecutive in one of the alignments. The weight of such an edge is thus set to 1 if it was not already present, and is incremented otherwise. Once the final graph is built, the tag associated to an edge between two vertices represents the number of reads supporting the connexion between these two vertices.

The consensus sequence of the original read can then be computed, by applying a dynamic programming algorithm to the graph, in order to find the highest weighted path between the vertices corresponding to the first and to the last tags associated to the original read. The bases associated to the fourth field of the tags along that path thus dictate the consensus sequence.

FalconSense thus produces split corrected long reads. Indeed, the consensus sequence of a read is computed by searching for a path in the graph, between the first and the last tag associated to this read. Since these tags are computed according to the alignments associated to this read, its uncovered extremities result in a lack of tags, and thus in a lack of vertices and edges in the graph. As a result, these uncovered extremities cannot be included in the consensus sequence computation, and are thus not reported. Moreover, internal regions of a given read which are not covered by alignments also imply a lack of tags, vertices, and edges in the graph. In this case, several traversals are thus performed in the different subgraphs corresponding to covered regions of this read. Several consensus sequences are thus computed independently for each of these regions.

#### Sparc (2016)

Sparc starts by computing the alignments between the reads with the help of BLASR. For each read, a modified de Bruijn graph is then built, from its set of similar reads, determined during the alignment step. Unlike a classical de Bruijn graph, the vertices of this graph are not only defined from the *k*-mers of the reads, but also from the positions of these *k*-mers in the read. Two independent vertices are thus built for two identical *k*-mers, if they are located at different positions, unlike a classical de Bruijn graph, which only defines a unique vertex in such a case. This necessity to create independent vertices for a same *k*-mer located at different positions can be explained by the small *k* values used by Sparc (*k* ≤ 3 in practice). Moreover, representing identical *k*-mers at different positions with distinct vertices also allows to avoid cycles in the graph, and thus compel it to be a DAG. Although the values of *k* are small, this graph is different from the DAG used in PBDAGCon, as the latter only stores bases of the reads, and not small *k*-mers.

Moreover, Sparc builds a sparse graph, and only stores one *k*-mer in every *n* bases. Edges between the vertices represent consecutive *k*-mers of the reads, and are labeled with the sequences that allow to link the corresponding vertices. The graph is also weighted, each edge being weighted with the number of reads supporting it. The graph is the initialized from the original read being processed, setting the weight of every edge to 1, and is then iteratively updated according to the alignments associated to this read, in the same way as in PBDAG-Con. The weight of the edges are thus incremented if a given alignment corresponds to an existing region of the graph, and new vertices and edges are created otherwise. Once the final graph is built, a consensus sequence can be computed, by searching for the highest weighted path, between the first and the last vertex associated to the original read, with the help of a dynamic programming algorithm, in the fashion as PBDDAGCon.

Sparc thus produces untrimmed corrected long reads. Indeed, the consensus sequence of a read is obtained by searching for a path between the first and the last vertex associated to this read. Since these vertices represent respectively the first and the last *k*-mer of the read, the entire length of the reads is preserved after correction. As for PBDAGCon, bases from the original reads than could not be corrected, due to a lack of coverage, are not reported in a different case from the corrected bases. As a result, the corrected long reads cannot further be trimmed or split after correction.

Moreover, Sparc also proposes a hybrid mode, allowing to use complementary short reads during the alignment step. In this case, the correction process globally remains unchanged. The major difference is that higher weights are associated to edges which are supported by short reads, so they have higher chances of being chosen during the consensus sequence computation.

#### Canu (2017)

The correction module of the assembler Canu is an updated version of PBcR-MHAP. During the mapping step, the creation of the sketches is altered, in order to adjust the probability whether to include or not certain *k*-mers. Indeed, a *k*-mer that appears frequently in the set of reads should have a lower weight, as it is not representative of the true origin of a read within the genome. In contrast, a *k*-mer which is present in few reads but that appears several times in a single read should have a higher weight, as it represents a greater proportion of the length of this read. A combination of these weights, called tf-idf weight (term frequency inverse document frequency) is thus applied to each *k*-mer of the reads.

To this aim, for each entry of a read sketch, instead of applying a single hash function to each *k*-mer, *N* hash functions are applied, where *N* corresponds to the tf-idf weight of the *k*-mer. This way, *k*-mers having a high tf-idf weight are hashed several times, which increases their chances of being part of the sketch. This application of the tf-idf weight to the sketches allows to increase the sensitivity of the overlaps, and to decrease the number of repetitive overlaps. The second iteration of the process, allowing to better estimate the similarity between overlapping regions, is also altered compared to PBcR-MHAP. In particular, a single hash function is used to build the sketches of the overlapping regions.

As in PBcR-MHAP, the overlaps are then used to correct the reads. This step is also improved, and allows to select an optimal subset of overlaps to use for the correction of each read, via a strategy similar to that of the hybrid version of PBcR. However, since the base to base alignments are unavailable, the score associated to each overlap is computed according to its length and to the estimate similarity between the two sequences. Each read is thus only authorized to participate to the correction of its *N* most similar reads, where *N* represents the sequencing depth of the reads. This approach allows to reduce the bias induced by repeated regions, by avoiding to always cover these regions with the same sets of reads.

The retained overlaps of each read are then used for correction. The correction is performed with the help of FalconSense, which has been adopted as the default algorithm, while PBDAGCon support has been discarded. Moreover, the FalconSense algorithm is also slightly altered. In particular, the edges of the graph which have a low weight are removed, to ensure that only well supported edges are traversed during the consensus sequence computation.

The Canu correction module thus produces split corrected long reads, for the reasons mentioned in the description of FalconSense.

#### MECAT (2017)

MECAT also relies on a mapping strategy to compute the overlaps between the reads. This strategy is based on the determination of *k*-mers shared between the reads, in order to define the overlaps. To this aim, the reads are first divided in several blocks, of 1,000 to 2,000 bps. The reads are then indexed in a hash table, using their *k*-mers as keys, and the positions of these *k*-mers in the blocks as the values. To find overlaps between the reads, the *k*-mers from the blocks associated to the reads are processed and queried in the hash table. A block from a given read thus overlaps a block from another read if these two blocks share a sufficient number of *k*-mers. By extension, a given read overlaps another read if at least a pair of blocks from these two reads do overlap.

A filtering step is then applied, via a scoring system, in order to filter out excessive and uninformative overlaps. To this aim, *k*-mers pairs are considered within two overlapping blocks. A distance, called DDF (Distance Difference Factor), is then computed between the pairs of *k*-mers that are shared by the two blocks. This distance is computed according to the occurrence positions of the *k*-mers within their respective block. A distance shorter than a given threshold indicates that the *k*-mers are supporting each other. In such a case, the scores of these *k*-mers are incremented. The *k*-mer having the highest score within a block is then defined as a seed for the following step, if this score is high enough. If several *k*-mers share the same highest score, one of these *k*-mers is randomly chosen.

The scoring system is then extended from the current block to its neighbor blocks, after obtaining the seed. The DDF are computed between this seed and the *k*-mers from the neighbor blocks. As before, if the DDF of a *k*-mers pair is below a given threshold, the score of the seed is incremented. If more than 80% of the DDFs of a neighbor block are below that threshold, the block is marked, and the scores of its *k*-mers are not computed. If some blocks remain unmarked after an iteration, the whole process is repeated on these blocks, and the score of their *k*-mers are computed, as described in the previous paragraph.

After computing all the scores of the *k*-mers that are shared by two overlapping reads, a base to base alignment of these two reads can be computed. To this aim, the *k*-mers of these reads are ordered according to their scores. The *k*-mers displaying the highest scores are then used as seeds, in order to ease the alignment computation. Is the length of the final alignment is at least as large as the size of a block, and if the divergence between the two sequences is low enough, the alignment is considered as valid, and is reported.

To correct a given read, the reads overlapping it are sorted, and considered in the decreasing order of the cumulative scores of their *k*-mers, which are computed during the previous step, via their DDF. Base to base local alignments are thus computed between the read to correct, and the reads overlapping it, in decreasing order of their scores. Moreover, in order to avoid the bias related to chimeric reads and to repeated sequences, an alignment between two reads is filtered out if its length is shorter than 90% of the length of the shortest read. The local alignments computation stops once 100 alignments have been produced, or when all the reads have been processed. Since the DDF score allows to estimate the length of the overlaps, computing local alignments only between the read to correct and its overlapping reads displaying the highest scores allow to gather sufficient information to perform a quick correction, all the while avoiding the computation of repetitive alignments, that would only bring little additional information.

A new correction strategy, combining the principles of both PBDAGCon and FalconSense is then applied. To this aim, the alignments related to the read to correct are summarized in a consensus table that contains the counts of matches, insertions, and deletions, for each position of the read. This table is then browsed, in order to determine three different types of regions, that require different correction approaches.

1. Trivial regions, which are composed of consistent matches and of a small number of insertions (usually *<* 6);
2. Regions which are composed of consistent deletions, and of a small number of insertions (*<* 6);
3. More complex regions, composed of a greater number of insertions (*≥* 6).

For the two first types of regions, correction is performed by determining the consensus base according to the counts stored in the table. For the third type of regions, a DAG is built, and the consensus is computed by searching for the highest weighted path, via a dynamic programming approach. Since such regions are usually short (*<* 10 bps), consensus can be computed quickly from the DAG.

MECAT thus produces split corrected long reads. Indeed, the blocks of the reads for which no overlap could be found cannot be corrected, whether they are located at the extremities or in internal regions of the reads. These blocks are thus removed from the corrected reads sequences, and consecutive blocks for which overlaps could be found are reported independently.

#### FLAS (2019)

FLAS is a wrapper of the MECAT algorithm, which we presented previously. It allows to polish the overlaps before correction, but also to use corrected regions of the reads in order to correct their uncorrected regions. First, the overlaps between the reads are computed with the MECAT mapping strategy. A directed graph is then built from these overlaps. Each vertex of this graph represents a read, and an edge is present between two vertices *u* and *v* if the reads associated to these vertices share at least an overlap, over a sufficient length. Once the graph is built, its maximal cliques are computed, with the help of the Bron-Kerbosch algorithm [10, 20, 21]. Pairs of maximal cliques that share common vertices are then processed, in order to find additional overlaps between the corresponding reads, or to remove erroneous overlaps. Three different cases can indeed be at the origin of common vertices in a given pair of maximal cliques. These common vertices thus require specific handling.

1. The reads from the two cliques originate from the same genome region, and the overlaps between the reads associated to the common vertices are thus correct;
2. The reads from the two cliques originate from different genome regions, but the reads associated to the common vertices span these two regions, and their overlaps are thus correct;
3. The reads from the two cliques originate from different genome regions, and the reads associated to the common vertices originate from a single of these two regions, and share erroneous overlaps with the reads originating from the other region.

These differences can be determined by comparing the number of common vertices, *s*, to the number of vertices of the smallest maximal clique of the pair, *c*. Indeed, among all the overlaps, the number of correct overlaps is usually larger than the number of erroneous overlaps. As a result, if *s* represents less than 50% of the vertices of *c*, the common vertices are considered as belonging to the third case.

Otherwise, common vertices are considered as belonging to the first or second case. To distinguish between these two cases, the number of reads associated to the common vertices that have a region overlapping the reads from one clique, and a different region overlapping the reads from the other clique, is computed. If this number represents at least 50% of the set of the common vertices of the two cliques, the common vertices are considered as belonging to the second case. Otherwise, the common vertices are considered as belonging to the first case.

For the maximal cliques belonging to the first case, the overlaps between the reads associated to the vertices of the cliques are recomputed, in order to produce additional overlaps, and thus be able to correct larger regions of these reads.

For the maximal cliques belonging to the second case, the reads associated to the common vertices span the two genome regions corresponding to the cliques. No supplementary alignment can thus be computed, and no specific action is necessary.

Finally, for the maximal cliques belonging to the third case, the common vertices have to be assigned to a single of the two cliques. To this aim, each common vertex is considered independently. The average similarity of the overlaps between the read associated to the vertex and the other reads of each clique is then computed. The vertex is thus removed from the clique with which it displays the lowest average similarity.

Once the overlaps have been polished, according to these three different cases, the MECAT algorithm is used to perform error correction of the reads.

A second correction step is then applied, where the corrected regions of the reads are used in order to correct the regions that could not be corrected during the first step. To this aim, the overlaps between the corrected regions of the reads are computed with the MECAT mapping strategy. A graph representing the overlaps is then built, in the same fashion as in the previous step. The reads are then aligned to this graph, in order to perform correction. For reads composed of both corrected and uncorrected regions, the corrected regions can be uniquely aligned to the graph, and thus be used as anchor points. The uncorrected regions can then be aligned along paths that allow to link together the corrected regions, and can thus be corrected with the help of the information contained in the followed paths. Moreover, reads solely composed of uncorrected regions can also be aligned to paths of the graph, as it is generated from corrected data, and thus contains a small number of errors. In cases where an uncorrected region of a read aligns with several paths, the path to which the region aligns with the highest identity is chosen as the correction. If no alignment stands out, the path which is supported by the greatest number of reads is chosen.

FLAS thus produces split corrected long reads. Indeed, FLAS relies on MECAT’s mapping and correction strategies. The regions (or blocks, as defined in MECAT) of the reads for which no overlap is found thus cannot be corrected. These regions are thus removed from the corrected reads sequences whether their are located at the reads extremities or not. Consecutive regions for which overlaps could be found are thus reported independently. However, unlike MECAT, FLAS also produces a trimmed version of the corrected reads, which retains the internal regions that could not be corrected due to a lack of overlaps.

#### 2.2.2 De Bruijn graphs

This approach is similar to the hybrid correction approach using de Bruijn graphs, mentioned in Section 2.1.3. In a first step, the graph is built from the long reads *k*-mers, and in a second step, the graph is traversed in order to find paths allowing to correct unanchored regions of the long reads. The main difference with the hybrid approach comes from the fact that the graph is, here, only constructed from the solid *k*-mers from the long reads. The methods adopting this approach mainly differ by the scale at which the graph is build. On the one hand, it can either be built globally, by studying the frequency of all the *k*-mers appearing in the reads. On the other hand, it can be built locally, by first computing overlaps between the long reads, in order to define small similar regions of the long reads, and then building small, local graphs at the scale of these regions.

#### LoRMA (2016)

LoRMA adapts the principle of the hybrid correction tool LoRDEC to the problem of self-correction. The graph is thus built from the solid *k*-mers of the reads. The reads can then be anchored to the graph, with the help of their solid *k*-mers. In the same fashion as LoRDEC, regions composed of weak *k*-mers, and bordered by regions composed of solid *k*-mers, are corrected by searching for paths of the graph that allow to link an anchor from the left flanking region to an anchor of the right flanking region. In a similar way, regions composed of weak *k*-mers, and located at the extremities of the reads, are corrected by using a neighbor solid *k*-mer as anchor point on the graph, and by traversing the graph, using the same stopping conditions as LoRDEC.

The principle of FMLRC, to perform several passes of correction, with increasing values of *k*, is also adopted in LoRMA. Given the initial error rates of the reads, the first correction pass is performed with a small value of *A. k.* Although such small values of *k* do not allow to correct large weak *k*-mers regions, due to the high number of branching paths in the graph, small regions can nevertheless be corrected. Hence, after each correction pass, solid *k*-mers regions grow longer, and a larger value of *k* can thus be used during the following passes. Moreover, the number of branching paths of the graph also decreases, and larger weak *k*-mers can thus be corrected. By iteratively reaching high values of *k*, the whole reads can thus eventually be corrected. Unlike FMLRC, LoRMA however does not use a variable order de Bruijn graph, but instead builds a new graph at each step, according to the chosen value of *k*.

The corrected reads are then split: regions only composed of weak *k*-mers are removed, in order to retain only regions solely composed of solid *k*-mers. The correction of these split reads is then polished via a second correction phase, relying on multiple sequence alignments. To this aim, a de Bruijn graph is built from the corrected reads *k*-mers, without setting any solidity threshold. The simple paths (*i.e.* with no branching paths) of the graph are then enumerated, in order to define the path of the graph which dictates the sequence of each read. Each path is thus composed of a set of simple paths, called segments. For each of these segments, the set of reads that traverse it is saved. Reads that are similar to a given read can thus be retrieved with the help of the graph, via their common *k*-mers, by following the path associated to this read, and selecting the reads that traverse common segments. This subset of similar reads is then filtered, in order to only retain the reads that share a sufficient number of *k*-mers with the initial read.

A consensus sequence can then be computed, from the original read and its set of similar reads. To this aim, the consensus sequence is first initialized as the initial read itself. This consensus is then iteratively updated, by aligning the similar reads. Each of these similar reads is thus considered independently, and aligned to the current consensus. The current multiple sequence alignment is thus updated, and the consensus is then, in turn, updated according to the multiple sequence alignment, via a majority vote at each position. This process is thus repeated until all the similar reads have been considered and aligned. The final multiple sequence alignment is then browsed, and the consensus positions supported by at least two reads are used to polish the correction of the initial read.

Due the the splitting step performed after the correction pass with the increasing-size de Bruijn graphs, LoRMA produces split corrected long reads. Moreover, during the multiple sequence alignment step, bases covered by less than two other reads are reported in a different case from bases covered by a greater number of reads. As a result, the corrected reads can easily further be trimmed or split after correction.

#### Daccord (2017)

Although it also relies on de Bruijn graphs, Daccord differentiates itself from LoRMA, and does not built a unique graph from the solid *k*-mers of the whole set of reads. Indeed, it rather builds a set of local graphs, on small regions of subsets of similar reads, defined after a preliminary alignment step. Daccord thus starts by computing the alignments between the reads, with the help of DALIGNER. An alignment between two reads *A* and *B* is thus represented by an alignment tuple (*A_b_, A_e_, B_b_, B_e_, S, E*), where:

- *A_b_* and *A_e_* represent, respectively, the beginning and ending positions of the alignment, on *A*;
- *B_b_* and *B_e_* represent, respectively, the beginning and ending positions of the alignment, on *B*;
- *S* represents the orientation of *B* relatively to *A* (forward or reverse-complement);
- *E* represent the edit script (*i.e.* the sequence of edit operations allowing to transform *A*[*A_b_*..*A_e_*] into *B*[*B_b_*..*B_e_*], if *S* = 0, or into *B*[*B_b_*..*B_e_*] if *S* = 1 (where *B* represents the reverse-complement of *B*).

A set of alignment tuples between a given read *A* and other reads thus define an alignment pile for the read *A.* The alignment pile of a given read *A* thus allows to represent the set of reads that align to *A*, and that are required for the construction of the local de Bruijn graph allowing to correct this read. These graphs are defined on small windows of less than 100 bps of the alignment pile. With the help of the information contained in this alignment pile, the sequences of the read that need to be included in the different windows can easily be extracted. Moreover, only the sequences from the reads that fully span a given window are used to build its associated de Bruijn graph. In particular, reads whose alignments only begin or end inside this window are thus not considered. Furthermore, the windows can be overlapping, and do not necessarily partition the alignment pile into a set of non-overlapping intervals.

Each window of the alignment pile is then processed independently. First, the de Bruijn graph of the window is built from the *k*-mers of factors of the reads that are included in this window, and that respect the aforementioned conditions. As for the previously described methods that rely on de Bruijn graphs, a source vertex and a target vertex are defined in order to guide the traversal of the graph, which yields a consensus sequence for the window. The source vertex is defined, in the left part of the window, as the most frequent first *k*-mer found in the factors of the reads included in the window. Symmetrically, the target vertex is defined, in the right part of the window, as the most frequent last *k*-mer found in the factors of the reads included in the window. Several pairs of such *k*-mers are however considered, in order to reduce the number of cases in which no path could be be found.

Unlike other methods based on the traversal of de Bruijn graphs that were previously presented, the graph is not traversed from the source towards the target nor from the target towards the source. Indeed, Daccord rather traverses the graph in both directions simultaneously, and searches for common vertices that allow to join the two paths. The traversals can thus yield a set of different paths. In this case, the consensus of the window is chosen by aligning the sequences of the obtained paths to the factor of the read *A* which is included in the window, and by choosing the sequence having the smallest edit distance with this factor. The procedure is thus repeated with all the other windows of the alignment pile of the read to correct.

Once the consensus sequences of all the windows have been computed, the read can eventually be corrected. To this aim, the consensus associated to each window of the alignment pile is aligned to the corresponding factor of the read. The edit scripts that allow to transform the factors of the read *A* associated to each window into their respective consensus sequences are thus computed. The windows are then once again considered independently, in order to define position pairs, from these edit scripts. Let a window beginning at position *b* on the read *A*, with an associated edit script *S*. Position pairs are defined as follows, considering operations from left to right:

- position pair (*b* + *i,* 0) is assigned to the *i*-th non-insertion operation in *S*;
- position pair (*b* + *i, −d*) is associated to the *d*-th insertion, before the *i*-th non-insertion operation in *S*.

Each position pair is annotated with the corresponding base of the window consensus. Once all the windows have been processed, position pairs are sorted, and the consensus sequence of the read is obtained with a majority vote on each position pair, by observing their associated bases.

Daccord thus produces split corrected long reads. Indeed, it is likely that the path searching in the graph does not yield a consensus for a given window. In this case, this windows cannot be considered during the realignment step of the consensus sequences to the initial read. As a result, its edit script, as well as the position pairs associated to it, cannot be determined. These missing consensus sequences thus result in missing position pairs in the sequence of ordered position pairs. The consensus sequence of the read thus cannot be computed on these missing position pairs, and multiple consensus sequences, corresponding to sequences of consecutive position pairs, are produced independently.

#### 2.2.3 Combination of strategies

As for hybrid correction, some methods also rely on combinations of the two previously described strategies. For instance, CONSENT relies on both multiple sequence alignment and de Bruijn graphs, which it combines into a two-step error correction process.

#### CONSENT (2019)

CONSENT first computes overlaps between the reads, using a mapping approach, with the help of Minimap2 [48]. From these alignments, as in Daccord, alignment piles are then defined for each read. Small windows, of a few hundreds of bps, are then defined on these alignment piles. Each window is processed independently, in two different steps. First, a multiple sequence alignment strategy is used, in order to compute a consensus sequence. Then, the consensus further goes through a second correction step, in which a local de Bruijn graph is built and traversed, in order to further polish remaining errors.

For the multiple alignment step, unlike other methods that would compute independent, pairwise alignments between the window to correct and other windows of the pile, and then summarize them into a DAG, CONSENT computes an actual MSA between all the sequences. To this aim, it makes use of poaV2 [44, 43], an algorithm that computes MSA with the help of partial order graphs. These graphs are DAGs, and are used as data structures containing all the MSA information. For a given MSA, the graph is thus enriched at each step, by aligning the new sequences to it, and updating it with additional vertices and edges, if necessary. Unlike other methods, this strategy thus allows CONSENT to both compute MSA and built the DAG used for consensus computation at the same time.

However, poaV2 is resource consuming, even on windows of a few hundreds bps. To address this issue, CONSENT relies on an efficient segmentation strategy, that allows to compute MSA and consensus on much shorter sequences, thus greatly reducing computational costs. First, common *k*-mers between the sequences included in the window being processed are computed. Using a dynamic programming algorithm, the longest chain of consecutive *k*-mers, common to the sequences, is then computed. This way, MSA and consensus sequences only need to be computed on subsequences bordered by the *k*-mers of the chain. CONSENT then reconstructs the global consensus, by concatenating the local consensus and the *k*-mers that were used for segmentation.

After consensus computation, a few erroneous bases might remain. To further enhance the quality of the corrected window, CONSENT thus performs a second correction phase, where the consensus sequence is polished with the help of a de Bruijn graph. First, the graph is thus built from the solid *k*-mers of the sequences included in the window. Due to the small size of the windows, a small *k*-mer size (usually, *k* = 9) can also be used. As in other de Bruijn graph based methods presented previously, solid *k*-mers of the consensus are used as anchor points on the graph, and weak *k*-mer regions are corrected, by searching for paths of the graph that allow to link together two anchors. In the same fashion, extremities of the consensus are polished by following the highest weighted edges of the graph, until the length of the path reaches the length of the extremity to correct, or until a branching path is reached.

Finally, after computing and polishing the consensus of a given window, the obtained sequence is locally aligned to the original read, around the positions the window originates from, to actually perform correction. This process is thus applied to each window of the alignment pile, until the read is completely corrected.

CONSENT thus produces trimmed corrected long reads. Indeed, since windows cannot be defined on the uncovered extremities of the long reads, CONSENT cannot process and correct these extremities. They are thus removed from the sequences of the corrected long reads. Moreover, during the consensus polishing step, CONSENT reports bases corresponding to solid *k*-mers and bases corresponding to weak *k*-mers in a different case. The corrected long reads can thus easily further be split after correction.

### 2.3 Summary

In the section, we described the state-of-the-art of available methods for the error correction of long reads, whether they adopt a hybrid or a self-correction approach. Four main approaches were described for self-correction: the alignment of short reads to the long reads, the alignment of contigs obtained from short reads assembly and long reads, the use of de Bruijn graphs, and the use of Hidden Markov Models. Other methods also combine different strategies to benefit from their different advantages. Two approaches were described for self-correction: multiple sequence alignment, and use of de Bruijn graphs. As for hybrid correction, other methods also combine these two strategies to counterbalance their advantages and drawbacks. The algorithmic principles of each error correction method have also been described. Available hybrid correction tools are summarized in Table 1. Available self-correction tools are summarized in Table 2. These tables also recall the approaches these different methods rely on, the way they output the reads (*i.e.* trimmed / split or not), the year they were released, as well as the technology (PacBio or ONT) they were validated on, in their respective publications.

## 3 Qualitative comparison

In this section, we present an in-depth benchmark of available hybrid and self-correction tools described in Section 2, and summarized in Table 1 and in Table 2. The following methods are however excluded from the benchmark, for the reasons mentioned below:

- FalconSense: this tool could not be installed on any of the computers used in our experiments;
- HECIL: this tool did not manage to correct any read, and produced a set of corrected read identical to the set of uncorrected reads, for all the datasets;
- LSCplus: this tool is not available for download anymore;
- MiRCA: this tool stopped and reported an error during the correction of all the datasets;
- ParLECH: this tool could not be installed on any of the computers used in our experiments;
- PBcR: this tool stopped and reported an error during the correction of all the datasets;
- PBcR-BLASR and PBcR-MHAP: this two tools are preliminary versions of the correction module from the assembler Canu. Only the latter, which is the most recent, is thus evaluated in the benchmark;
- PBDAGCon: this tool did not manage to produce any corrected read for all the datasets;
- Sparc: this tool stopped and reported an error during the correction of all the datasets;
- Sprai: this tool stopped and reported an error during the correction of all the datasets.

### 3.1 Datasets

We evaluated these correction methods on a wide variety of datasets, composed of both simulated and real data, displaying various error rates, read length and sequencing depths, and ranging from small bacterial to large mammal genomes. These datasets are summarized in Table 3.

**Table 3:** Characteristics of the datasets used during the experiments. ^1^http://www.genoscope.cns.fr/externe/nas/datasets/MinION/acineto/ ^2^http://www.genoscope.cns.fr/externe/nas/datasets/MinION/ecoli/ ^3^http://www.genoscope.cns.fr/externe/nas/datasets/MinION/yeast/ ^4^ Only reads from chromosome 1 were used. This dataset contains ONT ultra-long reads, reaching lengths up to 340 kbp. ^5^http://www.genoscope.cns.fr/externe/nas/datasets/Illumina/acineto/ ^6^http://www.genoscope.cns.fr/externe/nas/datasets/Illumina/ecoli/ ^7^http://www.genoscope.cns.fr/externe/nas/datasets/Illumina/yeast/

| Dataset | Number of reads | Number of bases (Mbp) | Average length (bp) | Coverage | Error rate (%) | Accession |
| --- | --- | --- | --- | --- | --- | --- |
| **Simulated PacBio data** |  |  |  |  |  |  |
| *A. baylyi* 20x | 8,765 | 72 | 8,202 | 20x | 18.57 | - |
| *E. coli* 20x | 11,306 | 93 | 8,226 | 20x | 18.65 | - |
| *S. cerevisiae* 20x | 30,132 | 247 | 8,204 | 20x | 18.61 | - |
| *C. elegans* 20x | 244,277 | 2,004 | 8,204 | 20x | 18.62 | - |
| *E. coli* 30x | 16,959 | 140 | 8,235 | 30x | 12.29 | - |
| *S. cerevisiae* 30x | 45,198 | 371 | 8,216 | 30x | 12.28 | - |
| *C. elegans* 30x | 366,416 | 3,006 | 8,204 | 30x | 12.28 | - |
| *E. coli* 60x (low) | 33,918 | 279 | 8,211 | 60x | 12.28 | - |
| *S. cerevisiae* 60x (low) | 90,397 | 742 | 8,204 | 60x | 12.29 | - |
| *C. elegans* 60x (low) | 732,832 | 6,011 | 8,220 | 60x | 12.28 | - |
| *A. baylyi* 60x (medium) | 26,296 | 216 | 8,213 | 60x | 18.59 | - |
| *E. coli* 60x (medium) | 33,918 | 279 | 8,217 | 60x | 18.62 | - |
| *S. cerevisiae* 60x (medium) | 90,397 | 743 | 8,221 | 60x | 18.61 | - |
| *C. elegans* 60x (medium) | 732,832 | 6,022 | 8,218 | 60x | 18.60 | - |
| **Real ONT data** |  |  |  |  |  |  |
| *A. baylyi* real | 89,011 | 381 | 4,284 | 106x | 29.91 | Genoscope<sup>1</sup> |
| *E. coli* real | 22,270 | 134 | 5,999 | 29x | 20.54 | Genoscope<sup>2</sup> |
| *S. cerevisiae* real | 205,923 | 1,173 | 5,698 | 95x | 44.51 | Genoscope<sup>3</sup> |
| *C. elegans* real | 363,500 | 2,008 | 5,524 | 20x | 28.93 | ERR1802061 |
| *H. sapiens*<sup>4</sup> | 1,075,867 | 7,256 | 6,744 | 29x | 17.60 | PRJEB23027 |
| **Illumina data** |  |  |  |  |  |  |
| *A. baylyi* | 900,000 | 224 | 250 | 50x | 1 | ERR788913<sup>5</sup> |
| *E. coli* | 775,500 | 232 | 300 | 50x | 1 | Genoscope<sup>6</sup> |
| *S. cerevisiae* | 2,500,000 | 625 | 250 | 50x | 1 | Genoscope<sup>7</sup> |
| *C. elegans* | 20,057,100 | 5,000 | 250 | 50x | 1 | ART [30] |

### 3.2 Benchmark summary

To evaluate the quality of the correction provided by each tool, we used ELECTOR [54], a software specially developed for large scale error correction tools benchmark, that allows to assess both the correction of simulated and real data. In particular, it reports the error rates of the reads before and after correction, as well as the recall and the precision of the assessed tools, among other metrics, when ran on simulated data. With real data, it is able to perform remapping of the reads to the reference genome with the help of Minimap2 [48], and reports the average identity of the alignments as well as the genome coverage, among other metrics. It is also able to perform assembly of the corrected reads, with the help of Miniasm [47], and reports the number of aligned contigs, the NGA50, the NGA75, and the genome coverage, among other metrics. All tools were ran with default or recommended parameters. The command lines that were used for each correction tool are detailed in Supplementary Materials.

We present results in various subsections, according to the characteristics of the assessed datasets. In particular, we distinguish seven different categories of datasets.

1. Datasets with high error rate and high coverage. This corresponds to the *A. baylyi* real and *S. cerevisiae* real datasets of Table 3. We present these results in Section 3.3, Table 4.
2. Datasets with high error rate and low coverage. This corresponds to the *C. elegans* real dataset of Table
3. We present these results in Section 3.4, Table 5.
4. Datasets with medium error rate and low coverage. This corresponds to the *A. baylyi* 20x, *E. coli* 20x, *S. cerevisiae* 20x, *C. elegans* 20x and *E. coli* real datasets of Table 3. We present these results in Section 3.5, Tables 6 and 7.
5. Datasets with medium error rate and medium coverage. This corresponds to the *A. baylyi* 60x (medium), *E. coli* 60x (medium), *S. cerevisiae* 60x (medium) and *C. elegans* 60x (medium) datasets of Table 3. We present these results in Section 3.6, Table 8.
6. Datasets with low error rate and low coverage. This corresponds to the *E. coli* 30x, *S. cerevisiae* 30x and *C. elegans* 30x datasets of Table 3. We present these results in Section 3.7, Table 9.
7. Datasets with low error rate and medium coverage. This corresponds to the *E. coli* 60x (low), *S. cerevisiae* 60x (low) and *C. elegans* 60x (low) datasets of Table 3. We present these results in Section 3.8, Table 10.
8. Datasets containing ONT ultra-long reads. This corresponds to the *H. sapiens* dataset of Table 3. We present these results in Section 3.9, Table 11.

**Table 4:**
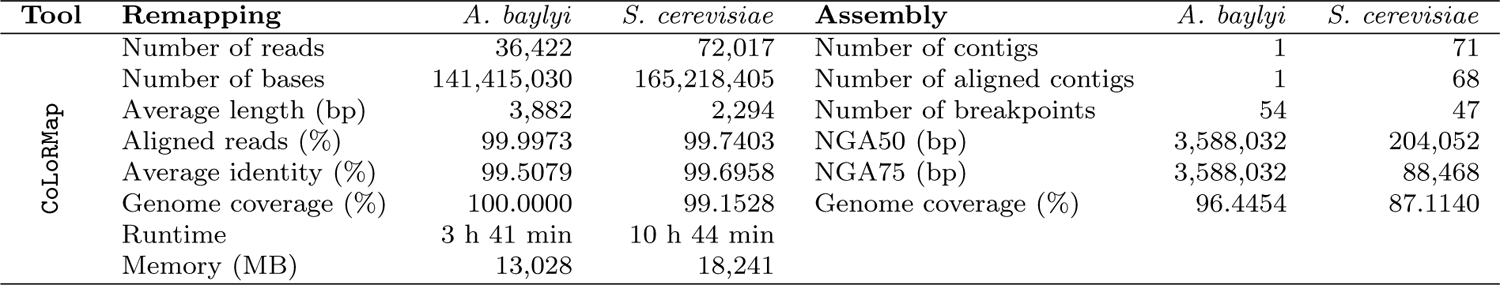
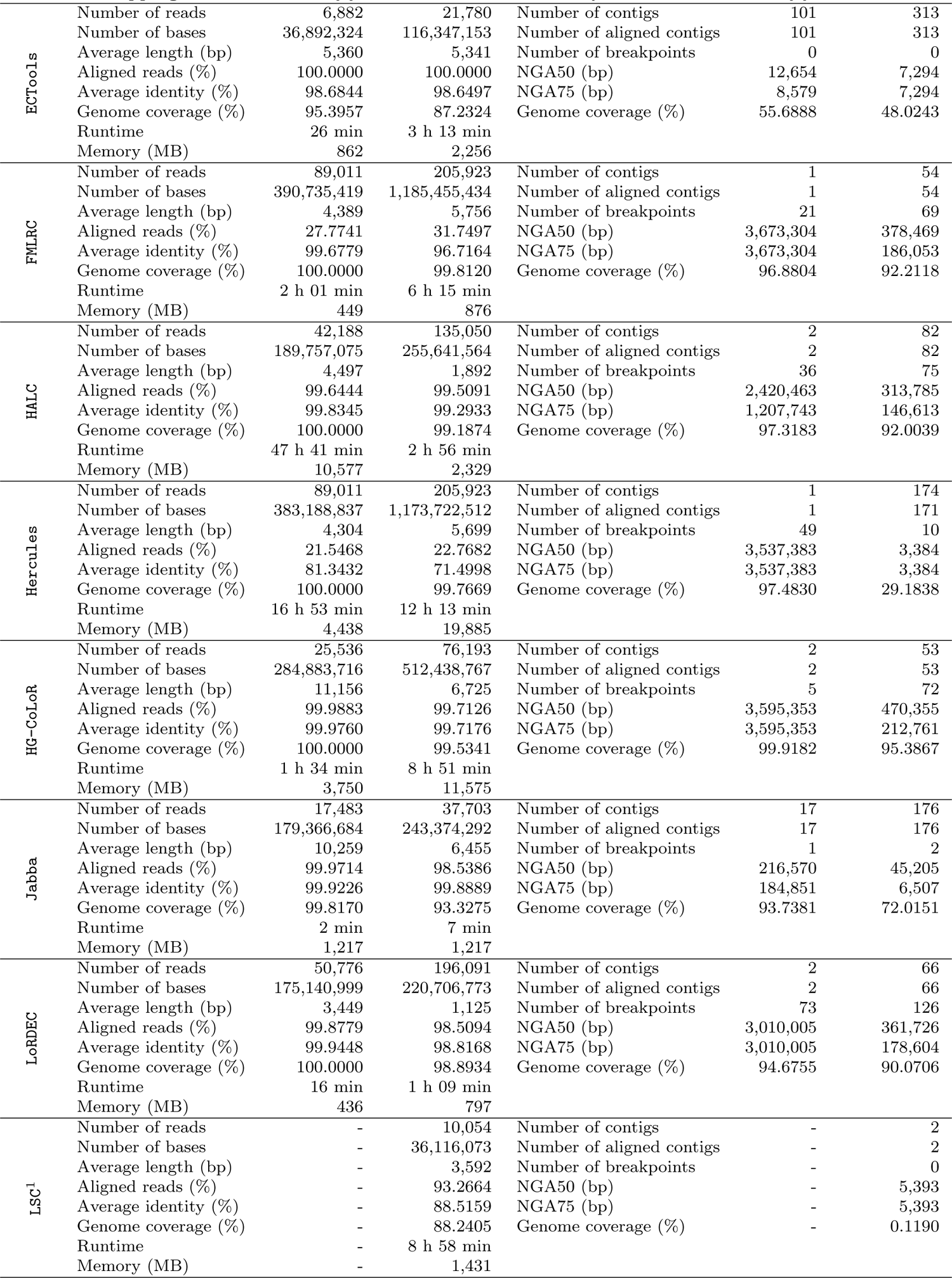
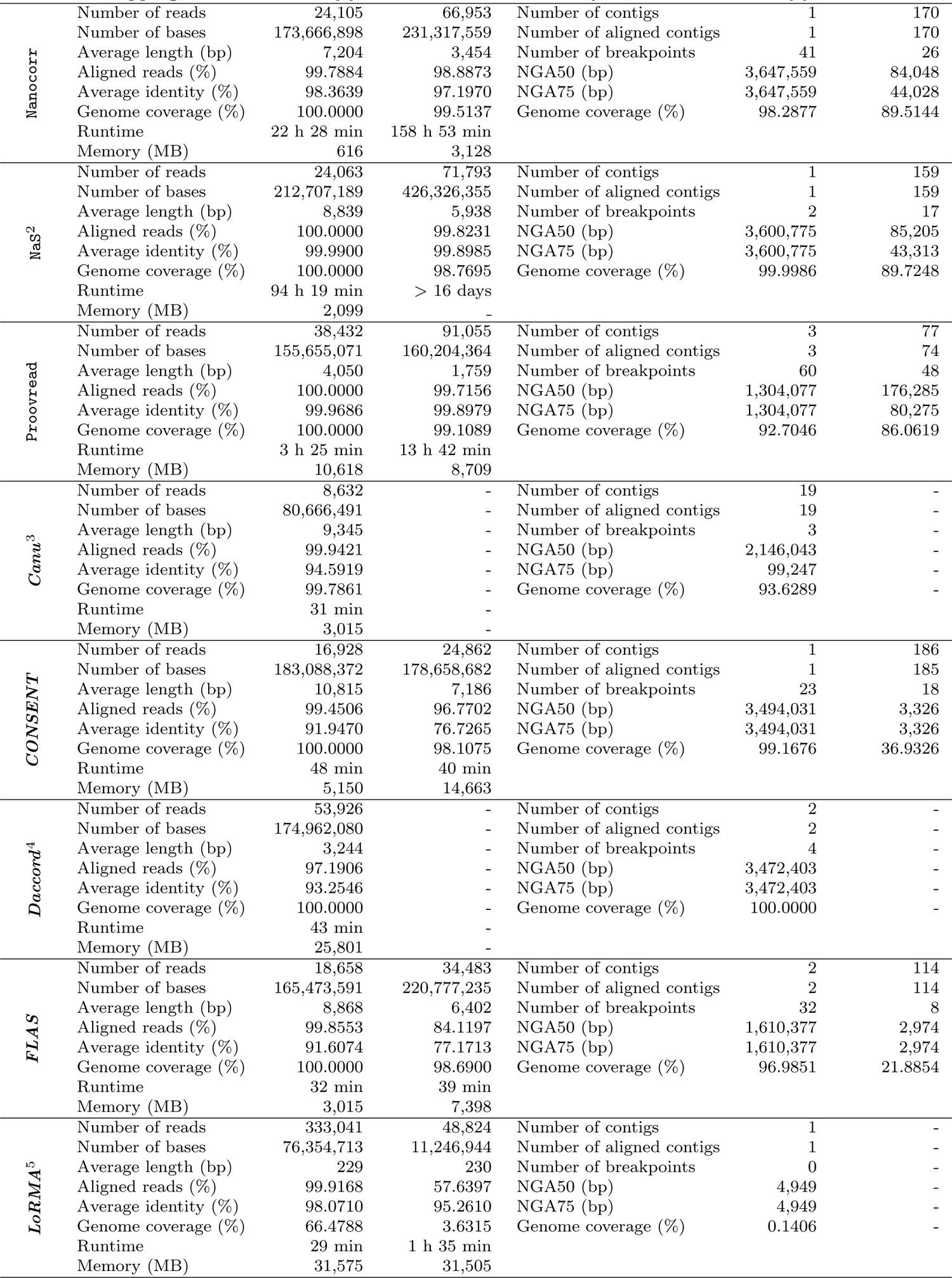
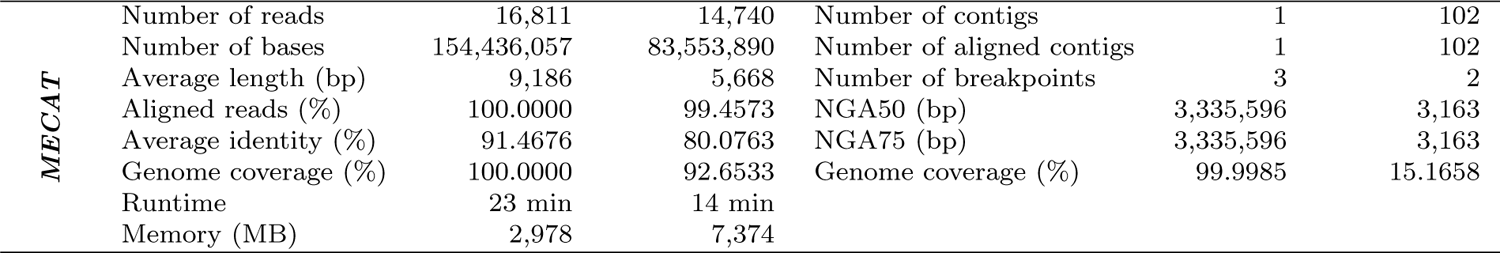
Comparison of the different error correction tools, on high error rate and high coverage datasets. This corresponds to the *A. baylyi* real and *S. cerevisiae* real datasets of Table 3. ^1^ LSC stopped and reported and error during the correction of the *A. baylyi* dataset. ^2^ NaS was stopped after 16 days on the *S. cerevisiae* datasets. Corrected long reads were obtained from the Genoscope website. ^3^ ***Canu*** stopped and reported an error during the correction of the *S. cerevisiae* dataset. ^4^ ***Daccord*** could not perform correction on the *S. cerevisiae* dataset, since the alignment step with DALIGNER required more than 128 GB of memory. ^5^ ***LoRMA*** corrected reads could not be assembled for the *S. cerevisiae* dataset.

**Table 5:** Comparison of the different error correction tools, on high error rate and low coverage datasets. This corresponds to the *C. elegans* real dataset of Table 3. ^1^ ECTools corrected long reads could not be assembled. ^2^ ***LoRMA*** corrected long reads could not be assembled.

| Tool | Remapping |  | Assembly |  |
| --- | --- | --- | --- | --- |
| CoLoRMap | Number of reads | 184,092 | Number of contigs | 1,246 |
|  | Number of bases | 418,509,369 | Number of aligned contigs | 1,237 |
|  | Average length (bp) | 2,273 | Number of breakpoints | 617 |
|  | Aligned reads (%) | 99.9473 | NGA50 (bp) | 58,411 |
|  | Average identity (%) | 98.6614 | NGA75 (bp) | 11,122 |
|  | Genome coverage (%) | 96.4079 | Genome coverage (%) | 76.6546 |
|  | Runtime | 91 h 18 min |  |  |
|  | Memory (MB) | 31,349 |  |  |
| ECTools<sup>1</sup> | Number of reads | 18 | Number of contigs | - |
|  | Number of bases | 54,323 | Number of aligned contigs | - |
|  | Average length (bp) | 3,017 | Number of breakpoints | - |
|  | Aligned reads (%) | 100.0000 | NGA50 (bp) | - |
|  | Average identity (%) | 98.4596 | NGA75 (bp) | - |
|  | Genome coverage (%) | 0.016 | Genome coverage (%) | - |
|  | Runtime | 81 h 25 min |  |  |
|  | Memory (MB) | 5,023 |  |  |
| FMLRC | Number of reads | 363,500 | Number of contigs | 1,047 |
|  | Number of bases | 2,063,059,647 | Number of aligned contigs | 1,042 |
|  | Average length (bp) | 5,675 | Number of breakpoints | 713 |
|  | Aligned reads (%) | 78.1986 | NGA50 (bp) | 91,545 |
|  | Average identity (%) | 96.5611 | NGA75 (bp) | 40,293 |
|  | Genome coverage (%) | 99.9932 | Genome coverage (%) | 82.9120 |
|  | Runtime | 7 h 54 min |  |  |
|  | Memory (MB) | 7,141 |  |  |
| Hercules | Number of reads | 363,500 | Number of contigs | 1,356 |
|  | Number of bases | 2,012,600,582 | Number of aligned contigs | 1,348 |
|  | Average length (bp) | 5,536 | Number of breakpoints | 425 |
|  | Aligned reads (%) | 72.8468 | NGA50 (bp) | 36,090 |
|  | Average identity (%) | 82.3142 | NGA75 (bp) | 690 |
|  | Genome coverage (%) | 99.9776 | Genome coverage (%) | 66.7194 |
|  | Runtime | 20 h 36 min |  |  |
|  | Memory (MB) | 32,000 |  |  |
| HG-CoLoR | Number of reads | 305,777 | Number of contigs | 989 |
|  | Number of bases | 1,567,516,029 | Number of aligned contigs | 988 |
|  | Average length (bp) | 5,126 | Number of breakpoints | 461 |
|  | Aligned reads (%) | 99.8653 | NGA50 (bp) | 96,058 |
|  | Average identity (%) | 99.5430 | NGA75 (bp) | 41,829 |
|  | Genome coverage (%) | 99.9655 | Genome coverage (%) | 82.6239 |
|  | Runtime | 83 h 10 min |  |  |
|  | Memory (MB) | 19,836 |  |  |
| Jabba | Number of reads | 270,929 | Number of contigs | 880 |
|  | Number of bases | 433,372,687 | Number of aligned contigs | 880 |
|  | Average length (bp) | 1,599 | Number of breakpoints | 15 |
|  | Aligned reads (%) | 97.2901 | NGA50 (bp) | 2,046 |
|  | Average identity (%) | 99.9430 | NGA75 (bp) | 2,046 |
|  | Genome coverage (%) | 95.2852 | Genome coverage (%) | 30.0434 |
|  | Runtime | 59 min |  |  |
|  | Memory (MB) | 13,362 |  |  |
| LoRDEC | Number of reads | 167,699 | Number of contigs | 21 |
|  | Number of bases | 221,676,779 | Number of aligned contigs | 16 |
|  | Average length (bp) | 1,321 | Number of breakpoints | 9 |
|  | Aligned reads (%) | 96.0137 | NGA50 (bp) | 589 |
|  | Average identity (%) | 98.2718 | NGA75 (bp) | 589 |
|  | Genome coverage (%) | 85.0782 | Genome coverage (%) | 0.0854 |
|  | Runtime | 1 h 15 min |  |  |
|  | Memory (MB) | 2,373 |  |  |
| Proovread | Number of reads | 364,244 | Number of contigs | 1,095 |
|  | Number of bases | 1,329,812,416 | Number of aligned contigs | 1,086 |
|  | Average length (bp) | 3,650 | Number of breakpoints | 509 |
|  | Aligned reads (%) | 99.7194 | NGA50 (bp) | 80,171 |
|  | Average identity (%) | 99.6065 | NGA75 (bp) | 31,114 |
|  | Genome coverage (%) | 99.9358 | Genome coverage (%) | 79.6994 |
|  | Runtime | 119 h 48 min |  |  |
|  | Memory (MB) | 20,254 |  |  |
| Canu | Number of reads | 340,826 | Number of contigs | 1,272 |
|  | Number of bases | 1,843,278,326 | Number of aligned contigs | 1,259 |
|  | Average length (bp) | 5,408 | Number of breakpoints | 512 |
|  | Aligned reads (%) | 76.7204 | NGA50 (bp) | 54,194 |
|  | Average identity (%) | 90.6784 | NGA75 (bp) | 416 |
|  | Genome coverage (%) | 99.9783 | Genome coverage (%) | 76.9836 |
|  | Runtime | 14 h 19 min |  |  |
|  | Memory (MB) | 9,178 |  |  |
| CONSENT | Number of reads | 256,568 | Number of contigs | 1,279 |
|  | Number of bases | 1,475,025,831 | Number of aligned contigs | 1,268 |
|  | Average length (bp) | 5,749 | Number of breakpoints | 624 |
|  | Aligned reads (%) | 96.7615 | NGA50 (bp) | 49,620 |
|  | Average identity (%) | 91.6888 | NGA75 (bp) | 1,606 |
|  | Genome coverage (%) | 99.7682 | Genome coverage (%) | 73.1300 |
|  | Runtime | 9 h 50 min |  |  |
|  | Memory (MB) | 20,382 |  |  |
| LoRMA<sup>2</sup> | Number of reads | 38,121 | Number of contigs | - |
|  | Number of bases | 10,277,321 | Number of aligned contigs | - |
|  | Average length (bp) | 269 | Number of breakpoints | - |
|  | Aligned reads (%) | 91.5506 | NGA50 (bp) | - |
|  | Average identity (%) | 97.6599 | NGA75 (bp) | - |
|  | Genome coverage (%) | 1.6269 | Genome coverage (%) | - |
|  | Runtime | 1 h 13 min |  |  |
|  | Memory (MB) | 32,275 |  |  |

**Table 6:** Comparison of the different error correction tools, on medium error rate and low coverage simulated datasets. This corresponds to the *A. baylyi* 20x, *E. coli* 20x, *S. cerevisiae* 20x and *C. elegans* 20x datasets of Table 3. ^1^ Nanocorr was not launched on the *C. elegans* dataset due to its large runtimes. ^2^ NaS was not launched on the *C. elegans* dataset due to its large runtimes. ^3^ ***Daccord*** could not perform correction on the *C. elegans* dataset, since the alignment step with DALIGNER required more than 128 GB of memory.

| Tool | Metric | Dataset |  |  |  |
| --- | --- | --- | --- | --- | --- |
|  |  | *A. baylyi* | *E. coli* | *S. cerevisiae* | *C. elegans* |
| CoLoRMap | Number of bases | 64,353,820 | 81,060,058 | 210,966,804 | 571,054,288 |
|  | Error rate (%) | 0.1336 | 0.1946 | 0.2655 | 2.6255 |
|  | Deletions | 71,639 | 165,412 | 434,983 | 14,745,290 |
|  | Insertions | 7,062 | 9,193 | 51,627 | 374,284 |
|  | Substitutions | 51,954 | 62,188 | 279,378 | 3,722,575 |
|  | Recall (%) | 99.9923 | 99.9890 | 99.9805 | 99.8445 |
|  | Precision (%) | 99.8707 | 99.8118 | 99.7413 | 97.4526 |
|  | Trimmed / split reads | 1,666 | 2,067 | 6,970 | 64,051 |
|  | Mean missing size (bp) | 939.2 | 1,497.5 | 1,396.2 | 3,234.8 |
|  | Extended reads | 8,463 | 10,730 | 28,117 | 96,953 |
|  | Mean extension size (bp) | 260.3 | 308.8 | 251.6 | 233 |
|  | Low quality reads | 12 | 20 | 121 | 332 |
|  | Short reads | 689 | 1,036 | 2,166 | 181,741 |
|  | Homopolymer ratio | 1.0000 | 1.0000 | 0.9826 | 0.9962 |
|  | Runtime | 57 min | 1 h 25 min | 4 h 42 min | 125 h 44 min |
|  | Memory (MB) | 6,198 | 9,659 | 13,544 | 32,188 |
| ECTools | Number of bases | 36,892,324 | 6,261,214 | 116,347,153 | 54,323 |
|  | Error rate (%) | 1.2086 | 1.3991 | 1.2428 | 1.4652 |
|  | Deletions | 77,464 | 13,883 | 246,662 | 123 |
|  | Insertions | 395,759 | 76,702 | 1,275,965 | 702 |
|  | Substitutions | 40,670 | 7,026 | 140,703 | 45 |
|  | Recall (%) | 98.8241 | 98.6228 | 98.8048 | 98.5410 |
|  | Precision (%) | 98.7923 | 98.6012 | 98.7581 | 98.5348 |
|  | Trimmed / split reads | 5,628 | 1,595 | 18,270 | 18 |
|  | Mean missing size (bp) | 2,730.3 | 4,103.1 | 2,747.6 | 4,770.3 |
|  | Extended reads | 0 | 0 | 0 | 0 |
|  | Mean extension size (bp) | 0 | 0 | 0 | 0 |
|  | Low quality reads | 0 | 0 | 0 | 0 |
|  | Short reads | 0 | 0 | 0 | 0 |
|  | Homopolymer ratio | 1.0064 | 1.0000 | 0.9991 | 1.0045 |
|  | Runtime | 17 min | 8 min | 57 h 12 min | 77 h 59 min |
|  | Memory (MB) | 862 | 917 | 2,256 | 5,028 |
| FMLRC | Number of bases | 64,715,552 | 83,732,334 | 223,529,739 | 1,832,773,801 |
|  | Error rate (%) | 0.1194 | 0.1930 | 1.0016 | 3.3582 |
|  | Deletions | 26,253 | 54,707 | 643,464 | 20,717,465 |
|  | Insertions | 89,640 | 185,196 | 2,289,467 | 52,872,229 |
|  | Substitutions | 22,637 | 45,795 | 546,972 | 16,628,231 |
|  | Recall (%) | 99.9142 | 99.8498 | 99.1797 | 97.8709 |
|  | Precision (%) | 99.8815 | 99.8081 | 99.0020 | 96.6643 |
|  | Trimmed / split reads | 20 | 28 | 114 | 3,090 |
|  | Mean missing size (bp) | 26.2 | 29.4 | 17.2 | 32.8 |
|  | Extended reads | 0 | 0 | 9 | 295 |
|  | Mean extension size (bp) | 0 | 0 | 290.8 | 31 |
|  | Low quality reads | 13 | 29 | 120 | 744 |
|  | Short reads | 0 | 0 | 0 | 0 |
|  | Homopolymer ratio | 1.0000 | 1.0000 | 0.9741 | 1.0000 |
|  | Runtime | 23 min | 29 min | 1 h 19 min | 8 h 02 min |
|  | Memory (MB) | 387 | 408 | 906 | 7,939 |
| HALC | Number of bases | 63,708,698 | 81,199,351 | 212,266,193 | 1,588,220,052 |
|  | Error rate (%) | 0.1113 | 0.1537 | 0.4212 | 1.5288 |
|  | Deletions | 59,157 | 115,973 | 1,035,978 | 35,970,722 |
|  | Insertions | 8,273 | 15,034 | 100,874 | 2,520,363 |
|  | Substitutions | 5,206 | 13,764 | 198,853 | 2,919,936 |
|  | Recall (%) | 99.9937 | 99.9918 | 99.9734 | 99.8945 |
|  | Precision (%) | 99.8909 | 99.8497 | 99.5872 | 98.4961 |
|  | Trimmed / split reads | 2,358 | 3,816 | 12,043 | 153,855 |
|  | Mean missing size (bp) | 341.7 | 547.3 | 577.5 | 777.2 |
|  | Extended reads | 24 | 24 | 71 | 772 |
|  | Mean extension size (bp) | 25.2 | 31.1 | 53.2 | 40.9 |
|  | Low quality reads | 2 | 2 | 160 | 2,582 |
|  | Short reads | 101 | 254 | 3,436 | 170,934 |
|  | Homopolymer ratio | 1.0000 | 1.0000 | 1.0580 | 0.9978 |
|  | Runtime | 22 min | 24 min | 1 h 19 min | 5 h 59 min |
|  | Memory (MB) | 597 | 965 | 2,115 | 2,323 |
|  |  | *A. baylyi* | *E. coli* | *S. cerevisiae* | *C. elegans* |
| Hercules | Number of bases | 70,375,149 | 91,064,066 | 242,189,693 | 1,976,944,911 |
|  | Error rate (%) | 10.5079 | 10.6411 | 10.9420 | 11.9249 |
|  | Deletions | 1,319,811 | 1,753,865 | 4,655,235 | 40,229,132 |
|  | Insertions | 7,040,655 | 9,211,444 | 25,099,150 | 216,151,231 |
|  | Substitutions | 1,113,866 | 1,447,112 | 4,003,822 | 35,147,508 |
|  | Recall (%) | 89.6376 | 89.4974 | 89.3519 | 88.3131 |
|  | Precision (%) | 89.4946 | 89.3613 | 89.0615 | 88.0792 |
|  | Trimmed / split reads | 8 | 14 | 42 | 706 |
|  | Mean missing size (bp) | 22.2 | 20.5 | 20.1 | 30.4 |
|  | Extended reads | 5 | 14 | 65 | 133 |
|  | Mean extension size (bp) | 22.6 | 24.6 | 194.1 | 23 |
|  | Low quality reads | 13 | 29 | 143 | 678 |
|  | Short reads | 0 | 0 | 0 | 0 |
|  | Homopolymer ratio | 0.9982 | 1.0000 | 1.0000 | 1.0014 |
|  | Runtime | 6 h 23 min | 7 h 47 min | 32 h 38 min | 199 h 42 min |
|  | Memory (MB) | 1,724 | 1,633 | 18,771 | 31,039 |
| HG-CoLoR | Number of bases | 65,065,102 | 84,089,814 | 219,744,436 | 1,726,223,265 |
|  | Error rate (%) | 0.0430 | 0.0691 | 0.2959 | 0.6524 |
|  | Deletions | 11,561 | 28,955 | 339,174 | 9,986,160 |
|  | Insertions | 15,803 | 33,409 | 548,419 | 5,156,404 |
|  | Substitutions | 1,869 | 3,147 | 79,127 | 775,253 |
|  | Recall (%) | 99.9991 | 99.9982 | 99.9900 | 99.9682 |
|  | Precision (%) | 99.9574 | 99.9315 | 99.7071 | 99.3554 |
|  | Trimmed / split reads | 133 | 498 | 4,562 | 71,079 |
|  | Mean missing size (bp) | 264.7 | 274.5 | 677.4 | 866.3 |
|  | Extended reads | 5,969 | 8,180 | 19,237 | 140,791 |
|  | Mean extension size (bp) | 67 | 71.1 | 72.5 | 72.2 |
|  | Low quality reads | 2 | 0 | 99 | 621 |
|  | Short reads | 0 | 4 | 496 | 8,849 |
|  | Homopolymer ratio | 1.0000 | 1.0000 | 0.9741 | 1.0000 |
|  | Runtime | 51 min | 51 min | 4 h 55 min | 88 h 10 min |
|  | Memory (MB) | 1,432 | 1,517 | 3,237 | 19,730 |
| Jabba | Number of bases | 68,653,634 | 87,851,697 | 216,794,630 | 1,480,673,578 |
|  | Error rate (%) | 0.0862 | 0.0636 | 0.1058 | 0.1909 |
|  | Deletions | 53,665 | 42,166 | 112,063 | 1,865,824 |
|  | Insertions | 9,913 | 7,041 | 95,757 | 649,951 |
|  | Substitutions | 1,919 | 466 | 21,954 | 102,639 |
|  | Recall (%) | 99.9977 | 99.9975 | 99.9973 | 99.9929 |
|  | Precision (%) | 99.9150 | 99.9370 | 99.8950 | 99.8106 |
|  | Trimmed / split reads | 639 | 1,204 | 6,387 | 87,670 |
|  | Mean missing size (bp) | 3,323.5 | 2,735.3 | 2,166.6 | 2,853.8 |
|  | Extended reads | 8,267 | 10,402 | 25,214 | 168,261 |
|  | Mean extension size (bp) | 850.4 | 922.7 | 1,080.5 | 1,043.1 |
|  | Low quality reads | 10 | 18 | 115 | 661 |
|  | Short reads | 179 | 386 | 2,954 | 34,239 |
|  | Homopolymer ratio | 1.0000 | 1.0000 | 0.9563 | 1.0000 |
|  | Runtime | 2 min | 2 min | 5 min | 45 min |
|  | Memory (MB) | 1,217 | 1,218 | 1,217 | 13,360 |
| LoRDEC | Number of bases | 60,892,408 | 77,969,503 | 188,228,237 | 1,154,508,245 |
|  | Error rate (%) | 0.0712 | 0.1474 | 0.5400 | 1.2643 |
|  | Deletions | 48,836 | 128,344 | 1,385,989 | 23,642,857 |
|  | Insertions | 8,473 | 21,205 | 122,252 | 1,584,355 |
|  | Substitutions | 6,144 | 17,896 | 229,925 | 1,845,645 |
|  | Recall (%) | 99.9921 | 99.9890 | 99.9483 | 99.8871 |
|  | Precision (%) | 99.9313 | 99.8570 | 99.4730 | 98.7542 |
|  | Trimmed / split reads | 2,951 | 4,786 | 22,470 | 210,051 |
|  | Mean missing size (bp) | 1,155.2 | 1,033.9 | 857.6 | 1,350.9 |
|  | Extended reads | 0 | 0 | 3 | 30 |
|  | Mean extension size (bp) | 0 | 0 | 1,285.3 | 156.9 |
|  | Low quality reads | 2 | 0 | 252 | 1,966 |
|  | Short reads | 260 | 737 | 22,811 | 474,725 |
|  | Homopolymer ratio | 1.0000 | 1.0000 | 0.9741 | 0.9995 |
|  | Runtime | 6 min | 8 min | 28 min | 6 h 01 min |
|  | Memory (MB) | 438 | 455 | 799 | 2,238 |
|  |  | *A. baylyi* | *E. coli* | *S. cerevisiae* | *C. elegans* |
| LSC | Number of bases | 37,111,831 | 44,310,080 | 122,224,269 | 771,778,062 |
|  | Error rate (%) | 5.8260 | 5.2598 | 6.1549 | 6.7716 |
|  | Deletions | 372,086 | 438,428 | 1,630,573 | 14,922,136 |
|  | Insertions | 1,835,695 | 1,991,268 | 6,295,946 | 40,404,903 |
|  | Substitutions | 253,915 | 286,969 | 895,587 | 5,874,295 |
|  | Recall (%) | 94.6923 | 95.2419 | 94.6340 | 94.7734 |
|  | Precision (%) | 94.2025 | 94.7706 | 93.8772 | 93.2623 |
|  | Trimmed / split reads | 5,990 | 7,307 | 20,257 | 147,403 |
|  | Mean missing size (bp) | 3,008.4 | 3,031.8 | 2,962.6 | 2,934.9 |
|  | Extended reads | 24 | 85 | 145 | 289 |
|  | Mean extension size (bp) | 22.7 | 24 | 24.3 | 26.1 |
|  | Low quality reads | 0 | 0 | 47 | 271 |
|  | Short reads | 599 | 944 | 2,453 | 29,243 |
|  | Homopolymer ratio | 1.0026 | 1.0000 | 0.9833 | 0.9938 |
|  | Runtime | 47 min | 47 min | 9 h 42 min | 117 h 34 min |
|  | Memory (MB) | 337 | 328 | 1,756 | 1,852 |
| Nanocorr<sup>1</sup> | Number of bases | 64,372,749 | 83,338,435 | 220,133,609 | - |
|  | Error rate (%) | 0.4099 | 0.3043 | 0.4723 | - |
|  | Deletions | 72,260 | 77,573 | 276,444 | - |
|  | Insertions | 301,605 | 261,194 | 1,023,357 | - |
|  | Substitutions | 54,832 | 57,152 | 258,241 | - |
|  | Recall (%) | 99.6491 | 99.7715 | 99.6679 | - |
|  | Precision (%) | 99.5928 | 99.6989 | 99.5322 | - |
|  | Trimmed / split reads | 1,574 | 1,602 | 5,076 | - |
|  | Mean missing size (bp) | 322.8 | 279.4 | 349.6 | - |
|  | Extended reads | 0 | 0 | 1 | - |
|  | Mean extension size (bp) | 0 | 0 | 20 | - |
|  | Low quality reads | 1 | 0 | 46 | - |
|  | Short reads | 0 | 0 | 0 | - |
|  | Homopolymer ratio | 1.0000 | 1.0000 | 1.0158 | - |
|  | Runtime | 2 h 52 min | 3 h 17 min | 39 h 52 min | - |
|  | Memory (MB) | 173 | 166 | 2,345 | - |
| NaS<sup>2</sup> | Number of bases | 58,963,977 | 78,034,042 | 207,085,378 | - |
|  | Error rate (%) | 2.1095 | 1.3035 | 2.7483 | - |
|  | Deletions | 714,139 | 530,425 | 2,102,677 | - |
|  | Insertions | 1,100,737 | 948,398 | 5,112,655 | - |
|  | Substitutions | 94,241 | 86,401 | 504,562 | - |
|  | Recall (%) | 99.9158 | 99.9468 | 99.9077 | - |
|  | Precision (%) | 97.8969 | 98.7009 | 97.2615 | - |
|  | Trimmed / split reads | 339 | 274 | 2,187 | - |
|  | Mean missing size (bp) | 2,361.4 | 2,370.8 | 2,437.7 | - |
|  | Extended reads | 6,688 | 8,070 | 22,658 | - |
|  | Mean extension size (bp) | 1,371.8 | 1,781.2 | 1,488.4 | - |
|  | Low quality reads | 1,651 | 1,751 | 5,040 | - |
|  | Short reads | 0 | 0 | 0 | - |
|  | Homopolymer ratio | 1.0000 | 1.0000 | 1.0000 | - |
|  | Runtime | 24 h 24 min | 28 h 48 min | 217 h 20 min | - |
|  | Memory (MB) | 2,099 | 2,099 | 2,099 | - |
| Proovread | Number of bases | 62,131,071 | 80,950,481 | 210,578,217 | 1,651,572,671 |
|  | Error rate (%) | 0.1122 | 0.1220 | 0.2075 | 0.5320 |
|  | Deletions | 73,995 | 127,118 | 670,850 | 21,888,508 |
|  | Insertions | 1,364 | 2,255 | 26,710 | 239,748 |
|  | Substitutions | 1,066 | 3,344 | 30,808 | 464,923 |
|  | Recall (%) | 99.9944 | 99.9938 | 99.9880 | 99.9599 |
|  | Precision (%) | 99.8897 | 99.8808 | 99.7964 | 99.4820 |
|  | Trimmed / split reads | 3,221 | 4,485 | 14,211 | 98,127 |
|  | Mean missing size (bp) | 299.5 | 268.1 | 332.5 | 599.5 |
|  | Extended reads | 11 | 18 | 45 | 242 |
|  | Mean extension size (bp) | 27.3 | 23.8 | 257 | 79.3 |
|  | Low quality reads | 1 | 0 | 124 | 1,357 |
|  | Short reads | 861 | 752 | 2,834 | 24,984 |
|  | Homopolymer ratio | 1.0000 | 1.0000 | 0.9669 | 0.9935 |
|  | Runtime | 52 min | 1 h 18 min | 4 h 24 min | 79 h 33 min |
|  | Memory (MB) | 5,784 | 6,029 | 10,435 | 20,213 |
|  |  | *A. baylyi* | *E. coli* | *S. cerevisiae* | *C. elegans* |
| *Canu* | Number of bases | 66,632,351 | 86,443,218 | 229,555,492 | 1,873,188,109 |
|  | Error rate (%) | 4.9496 | 5.2422 | 5.0619 | 4.9561 |
|  | Deletions | 738,704 | 1,011,985 | 2,574,320 | 20,233,048 |
|  | Insertions | 3,517,266 | 4,804,729 | 12,252,413 | 97,517,920 |
|  | Substitutions | 622,602 | 841,679 | 2,197,172 | 17,594,481 |
|  | Recall (%) | 95.2430 | 94.9490 | 95.1477 | 95.2663 |
|  | Precision (%) | 95.0573 | 94.7647 | 94.9459 | 95.0524 |
|  | Trimmed / split reads | 547 | 764 | 2,216 | 14,407 |
|  | Mean missing size (bp) | 27.1 | 27.2 | 35.1 | 29.5 |
|  | Extended reads | 53 | 66 | 178 | 1,462 |
|  | Mean extension size (bp) | 32.3 | 27.4 | 30.7 | 32.5 |
|  | Low quality reads | 0 | 0 | 43 | 305 |
|  | Short reads | 0 | 0 | 0 | 0 |
|  | Homopolymer ratio | 1.0055 | 1.0000 | 0.9937 | 1.0120 |
|  | Runtime | 12 min | 14 min | 32 min | 4 h 37 min |
|  | Memory (MB) | 2,974 | 2,821 | 3,274 | 6,950 |
| *CONSENT* | Number of bases | 48,390,086 | 61,487,428 | 166,101,116 | 1,359,131,353 |
|  | Error rate (%) | 8.2534 | 8.5423 | 8.2652 | 9.5548 |
|  | Deletions | 4,778,369 | 6,371,753 | 16,549,340 | 155,680,250 |
|  | Insertions | 1,613,389 | 2,076,177 | 5,404,368 | 50,056,903 |
|  | Substitutions | 114,628 | 142,283 | 455,785 | 6,084,369 |
|  | Recall (%) | 97.9538 | 97.9155 | 98.0349 | 97.9553 |
|  | Precision (%) | 91.8579 | 91.5687 | 91.8483 | 90.5794 |
|  | Trimmed / split reads | 8,332 | 10,835 | 28,621 | 230,249 |
|  | Mean missing size (bp) | 1,507.1 | 1,581.7 | 1,498 | 1,401 |
|  | Extended reads | 100 | 112 | 267 | 2,373 |
|  | Mean extension size (bp) | 133.8 | 110.3 | 106.9 | 92.2 |
|  | Low quality reads | 0 | 0 | 57 | 471 |
|  | Short reads | 0 | 0 | 0 | 0 |
|  | Homopolymer ratio | 1.0070 | 1.0000 | 0.9788 | 0.9765 |
|  | Runtime | 6 min | 8 min | 22 min | 3 h 49 min |
|  | Memory (MB) | 1,020 | 1,552 | 4,514 | 14,522 |
| *Daccord*<sup>3</sup> | Number of bases | 64,669,977 | 83,773,362 | 222,050,951 | - |
|  | Error rate (%) | 0.4694 | 0.3965 | 0.5447 | - |
|  | Deletions | 121,756 | 75,627 | 470,101 | - |
|  | Insertions | 275,390 | 321,532 | 977,499 | - |
|  | Substitutions | 25,146 | 33,860 | 200,541 | - |
|  | Recall (%) | 99.8647 | 99.8817 | 99.8591 | - |
|  | Precision (%) | 99.5371 | 99.6077 | 99.4630 | - |
|  | Trimmed / split reads | 328 | 119 | 991 | - |
|  | Mean missing size (bp) | 445.5 | 369.9 | 410.4 | - |
|  | Extended reads | 0 | 0 | 0 | - |
|  | Mean extension size (bp) | 0 | 0 | 0 | - |
|  | Low quality reads | 0 | 0 | 49 | - |
|  | Short reads | 17 | 4 | 54 | - |
|  | Homopolymer ratio | 1.0061 | 1.0000 | 0.9810 | - |
|  | Runtime | 16 min | 24 min | 1 h 10 min | - |
|  | Memory (MB) | 3,509 | 4,538 | 14,111 | - |
| *FLAS* | Number of bases | 35,567,295 | 44,048,436 | 125,103,108 | 662,199,783 |
|  | Error rate (%) | 2.1958 | 2.2848 | 2.4742 | 9.1804 |
|  | Deletions | 777,449 | 879,707 | 2,963,186 | 71,270,551 |
|  | Insertions | 590,885 | 801,154 | 2,362,512 | 40,983,626 |
|  | Substitutions | 75,331 | 89,736 | 322,562 | 9,060,762 |
|  | Recall (%) | 99.6847 | 99.6914 | 99.6456 | 99.2995 |
|  | Precision (%) | 97.8309 | 97.7424 | 97.5575 | 90.8881 |
|  | Trimmed / split reads | 4,992 | 6,372 | 17,066 | 110,818 |
|  | Mean missing size (bp) | 2,040.3 | 2,097.1 | 2,051.1 | 1,806.9 |
|  | Extended reads | 44 | 78 | 215 | 5,956 |
|  | Mean extension size (bp) | 196.4 | 343.2 | 280 | 198.2 |
|  | Low quality reads | 1,868 | 2,591 | 6,081 | 85,161 |
|  | Short reads | 97 | 149 | 339 | 2,725 |
|  | Homopolymer ratio | 0.9806 | 1.0000 | 0.9783 | 0.9964 |
|  | Runtime | 2 min | 3 min | 9 min | 1 h 13 min |
|  | Memory (MB) | 1,231 | 1,354 | 2,259 | 10,371 |
|  |  | *A. baylyi* | *E. coli* | *S. cerevisiae* | *C. elegans* |
| **LoRMA** | Number of bases | 676,120 | 884,592 | 6,279,242 | 11,394,637 |
|  | Error rate (%) | 0.6305 | 0.9261 | 1.6624 | 5.0838 |
|  | Deletions | 3,032 | 5,139 | 195,481 | 996,348 |
|  | Insertions | 1,112 | 744 | 21,776 | 83,683 |
|  | Substitutions | 1,156 | 2,136 | 14,201 | 110,710 |
|  | Recall (%) | 99.9766 | 99.9733 | 99.8771 | 99.7216 |
|  | Precision (%) | 99.3790 | 99.0871 | 98.3563 | 94.9446 |
|  | Trimmed / split reads | 340 | 667 | 2,990 | 36,351 |
|  | Mean missing size (bp) | 3,183.5 | 3,115.5 | 1,927.7 | 1,165.9 |
|  | Extended reads | 0 | 0 | 6 | 24 |
|  | Mean extension size (bp) | 0 | 0 | 1,383.2 | 195.4 |
|  | Low quality reads | 14 | 0 | 201 | 2,476 |
|  | Short reads | 446 | 1,248 | 8,683 | 125,774 |
|  | Homopolymer ratio | 1.0000 | 1.0000 | 1.0000 | 1.0000 |
|  | Runtime | 5 min | 5 min | 20 min | 3 h 42 min |
|  | Memory (MB) | 32,118 | 32,183 | 32,183 | 32,041 |
| **MECAT** | Number of bases | 46,854,371 | 58,979,203 | 162,057,920 | 870,965,775 |
|  | Error rate (%) | 0.5340 | 0.5243 | 0.6555 | 0.6540 |
|  | Deletions | 223,296 | 267,547 | 975,507 | 5,379,870 |
|  | Insertions | 66,725 | 82,121 | 263,724 | 1,566,201 |
|  | Substitutions | 5,883 | 7,575 | 52,975 | 139,668 |
|  | Recall (%) | 99.8289 | 99.8317 | 99.8015 | 99.8196 |
|  | Precision (%) | 99.4817 | 99.4915 | 99.3636 | 99.3597 |
|  | Trimmed / split reads | 4,802 | 6,304 | 16,034 | 120,325 |
|  | Mean missing size (bp) | 2,036.7 | 2,114.5 | 2,069.5 | 2,320.8 |
|  | Extended reads | 0 | 0 | 1 | 41 |
|  | Mean extension size (bp) | 0 | 0 | 23 | 354.7 |
|  | Low quality reads | 0 | 0 | 50 | 583 |
|  | Short reads | 0 | 0 | 0 | 1 |
|  | Homopolymer ratio | 1.0008 | 1.0057 | 0.9810 | 0.9938 |
|  | Runtime | 22 sec | 26 sec | 1 min 20 sec | 18 min 20 sec |
|  | Memory (MB) | 1,202 | 1,322 | 2,207 | 10,340 |

Additionally, for the sake of clarity, the names of hybrid correction tools are reported in normal font, while the names of self-correction tools are reported in italic. Moreover, in tables, we always report the results of hybrid correction tools first, and then the results of self-correction tools.

### 3.3 High error rate, high coverage

Here, we present results on the *A. baylyi* real and *S. cerevisiae* real datasets of Table 3. For these datasets, when possible, we evaluated the split corrected long reads. When split reads were not available, we evaluated the trimmed corrected long reads. Finally, when both split and trimmed reads were not available, we evaluated the complete corrected long reads. These results are summarized in Table 4 and commented below.

Most hybrid correction tools performed well and managed to reduce the error rates of the reads to around 1% or less, even on the highly noisy *S. cerevisiae* dataset, displaying an initial error rate of 44.51%. Actually, only LSC, and Hercules displayed poor performances, reaching error rates of 6.7% for the former, and up to 28.5% for the latter, on the *S. cerevisiae* dataset. Moreover, even on the *A. baylyi* dataset, containing a smaller proportion of errors, Hercules still displayed an error rate of 18.6%. In terms of assembly, all hybrid tools displayed a fairly low number of contigs, and covered the reference genomes quite well, except for ECTools, which only reached around 50% of genome coverage on the two datasets. A few additional tools yielded low quality assemblies on the *S. cerevisiae* dataset, namely Hercules, which only reached 29% of genome coverage, Jabba, which reached 72%, and especially LSC, which could not yield a proper assembly.

Conversely, self-correction tools displayed poor performances, and did not manage to greatly reduce the error rates of the reads for any of the datasets. Indeed, the lowest error rate is reached by ***LoRMA***, and is mostly caused by the fact it aggressively splits the reads, as suggested by their average length, reaching around 230 bp on the two datasets. Although accurate, these extremely short corrected reads could thus not yield a proper assembly on the *A. baylyi* dataset, and could not be assembled at all on the *S. cerevisiae*. Apart from ***LoRMA***, the lowest error rate was reached by ***Canu*** on the *A. baylyi* dataset, with 5.4%, and by ***MECAT*** on the *S. cerevisiae* dataset, with 19.9%. In terms of assembly, and apart from ***LoRMA***, all tools performed well on the *A. baylyi* dataset, except for ***Canu***, which only reached 93.6% of genome coverage. However, on the *S. cerevisiae* dataset, all tools displayed unsatisfying results, reaching greater number of contigs and smaller genome coverage than most hybrid correction tools. More precisely, the highest genome coverage was reached by ***CONSENT***, with only 36.9%.

In terms of runtime, all hybrid correction tools but LoRDEC and Jabba performed slower than self-correction methods. It is however worth noting that Jabba was the fastest tool, requiring at most 7 minutes to perform correction on the *S. cerevisiae* dataset. The largest runtime was reached by NaS, requiring more than 94 hours on the *A. baylyi* dataset, and more than 16 days on the *S. cerevisiae*, leading to the experiment being stopped before it finished. In comparison, the slowest self-correction tool only displayed a runtime of 1.5 hour on the *S. cerevisiae* dataset. Conversely, in terms of memory consumption, self-correction methods were more demanding than most hybrid correction methods. This can especially be seen with ***LoRMA*** requiring almost 32 GB on both datasets. Furthermore, on the *S. cerevisiae* dataset, only CoLoRMap and Hercules displayed a significantly higher memory consumption than other self-correction tools, ***LoRMA*** excluded.

These results thus suggest that, even with sequencing depth as high as 100x, self-correction methods cannot perform well on extremely noisy datasets, reaching error rates of 30% and more, which was common in first long-read sequencing experiments. Although more time consuming, hybrid correction methods should thus be preferred on such data, since they do manage to properly perform correction, efficiently lower the error rates of the reads, and even lead to satisfying assembly results.

### 3.4 High error rate, low coverage

Here, we present results on the *C. elegans* real dataset of Table 3. For this dataset, when possible, we evaluated the split corrected long reads. When split reads were not available, we evaluated the trimmed corrected long reads. Finally, when both split and trimmed reads were not available, we evaluated the complete corrected long reads. These results are summarized in Table 5 and commented below. Moreover, for these experiments, we excluded the following tools from the comparison, for the ensuing reasons:

- HALC, because it stopped and reported an error during correction;
- LSC, because of its large runtimes on previous experiments;
- Nanocorr, because of its large runtimes on previous experiments;
- NaS, because of its large runtimes on previous experiments;
- ***Daccord***, because it could not perform correction, since the alignment step with DALIGNER required more than 128 GB of memory;
- ***FLAS***, because it stopped and reported an error during correction;
- ***MECAT***, because it stopped and reported an error during correction.

On this dataset, conclusions that were drawn in Section 3.3 are further confirmed. Indeed, since the error rate remained high, but the sequencing depth only reached 20x, self-correction methods performed worse than previously. In particular, both ***FLAS*** and ***MECAT*** could not process this dataset, and reported an error during correction. Moreover, ***CONSENT***, the best-performing self-correction method on this dataset, only managed to reduce the error rate to 8.3%. These results are in accordance with the fact self-correction tools only rely on long reads, and thus necessitate larger sequencing depths to perform efficient correction.

On the other hand, hybrid correction tools, for which the sequencing depth of the long reads has no impact on correction, still performed well. However, since the studied genome was more complex than those of Section 3.4, corrected reads reached a slightly higher error rate, of a little more than 1% on average. As in the previous experiments, Hercules still displayed the highest error rate, with 17.6%. Conversely, HG-CoLoR and Jabba provided the best results, reducing the error rates of the reads to less than 0.5% and less than 0.1% respectively. Jabba however produced a higher number of split reads than HG-CoLoR, as can be seen from the average length of the reads. The assembly yielded from Jabba corrected reads thus only reached a genome coverage of 30%, while that yielded from HG-CoLoR corrected reads reached more than 82%. ECTools corrected reads could not be assembled, and LoRDEC corrected reads yielded an unsatisfying assembly, also due to the large proportion of split reads. Other methods yielded comparable assemblies, which were of higher quality than those obtained from reads corrected with self-correction methods. However, all assemblies were of lower quality than the ones presented in Section 3.3, which can be explained by the lower sequencing depth of the long reads.

In terms of runtime, hybrid correction tools were still much slower than self-correction tools. Indeed, ***Canu***, which was the slowest self-correction tool on this dataset, only required 14 hours to run, while hybrid correction tools such as CoLoRMap, ECTools, and HG-CoLoR required more than 80 hours. However, in the same fashion as in Section 3.3, hybrid correction tools LoRDEC and Jabba were the fastest, both only requiring around 1 hour to run. In terms of memory consumption, all hybrid and self-correction tools were comparable. However, the least demanding tools were the hybrid correctors ECTools and LoRDEC, which required, respectively, only 5 GB and 2 GB, while ***Canu***, the least demanding self-correction tool, required 9 GB. On the other hand, most demanding tools were CoLoRMap, Hercules and ***LoRMA***, which all required around 32 GB.

These results suggest that, when the error rate of the reads is high and the sequencing depth is low, self-correction methods cannot perform well. These observations are in accordance with those of Section 3.3, indicating that self-correction tools cannot perform well on high error-rates datasets, even with high coverage depth. Logically, lowering the coverage of the reads, thus only worsens the results provided by self-correction tools. As a result, although they are more time consuming, hybrid correction methods should be preferred on such data, since they do manage to properly perform correction, and since the sequencing depth of the long reads has no impact on their performances.

### 3.5 Medium error rate, low coverage

Here, we present results on the *A. baylyi* 20x, *E. coli* 20x, *S. cerevisiae* 20x, *C. elegans* 20x and *E. coli* real datasets of Table 3. For these datasets, when possible, we evaluated the split corrected long reads. When split reads were not available, we evaluated the trimmed corrected long reads. Finally, when both split and trimmed reads were not available, we evaluated the complete corrected long reads. These results are summarized in Table 6 for simuated data, and in Table 7 for real data. Results on simulated data are commented in Section 3.5.1, and results on real data are commented in Section 3.5.2.

#### 3.5.1 Simulated data

On these datasets, most hybrid correction tools managed to reduce the error rates to around 1% or less, on the *A. baylyi*, *E. coli* and *S. cerevisiae* datasets. In the same fashion as in Section 3.3, only LSC and Hercules displayed poor performances on these three datasets, displaying error rates up to 6.1% for the former, and up to 10.9% for the latter, on the *S. cerevisiae* dataset. Moreover, it is worth noting that NaS, which performed very well during the experiments of Section 3.3, did not manage to reduce the error rates of the reads below 1% on any of these datasets, and even reached error rates up to 2.7% on the *S. cerevisiae* datasets. Another interesting point is that all hybrid methods displayed performance drops on the *C. elegans* dataset, even though it displayed the same error rate as all the other datasets. On this dataset, only HG-CoLoR, Jabba and Proovread managed to reduce the error rate below 1%. As explained in Section 3.4 this can be explained by the fact that the genome of *C. elegans* is more complex and contains more repetitions than the genomes of the other species studied here.

As for self-correction, although the error rates of the original datasets have been divided by 1.5-2.4 compared to the experiments of Section 3.3 and Section 3.4, most methods still displayed poor performances. Indeed, of all the self-correction methods, only ***Daccord*** and ***MECAT*** managed to reduce the error rates of the reads below 1%. However, due to its high memory requirements, ***Daccord*** could not scale to the *C. elegans* dataset. Moreover, compared to hybrid correction methods, ***MECAT*** reported a significantly smaller number of bases, and thus corrected less reads. ***LoRMA*** performed well in terms of reduction of the error rates as well, but as previously mentioned, aggressively split the reads, as underlined by the trimmed / split reads and mean missing size metrics. Moreover, on the *C. elegans* dataset, most self-correction methods did not display the same performance drop as hybrid correction methods, apart from ***FLAS*** and ***LoRMA***. This underlines the fact that self-correction seems to be more suited for more complex, repeat-rich, genomes.

Moreover, it is also interesting to note that metrics reported by ELECTOR provide additional insight as to how the different tools behave. For instance, as previously mentioned, they show that ***LoRMA*** tends to split a vast majority of the reads, and that large proportions of the original reads are missing from these split corrected reads. More generally, it also shows that graph-based methods, through their traversal procedures, can also extend the reads beyond their initial extremities. This can especially be seen on the metrics reported for HG-CoLoR and Jabba. Graph-based methods can however also split the reads, as previously mentioned with ***LoRMA***, but as can also be seen with LoRDEC. On the other hand, alignment-based methods, whether hybrid or not, tend to split the reads rather than to extend them. This can be seen on metrics reported for HALC, LSC, ***FLAS*** and ***MECAT***. Finally, although most methods display comparable recall and precision, it is worth noting that some tools, such as ***CONSENT***, ***FLAS*** and NaS, display higher disparities. More precisely, these tools display a higher recall and a lower precision, meaning that they do manage to identify the errors correctly, but tend to fail to replace these erroneous bases with the correct ones.

In terms of runtime, all hybrid correction tools but LoRDEC and Jabba once again performed slower than self-correction methods. On all these datasets, the fastest tool was ***MECAT***, requiring as little as 22 seconds to perform on the *A. baylyi* dataset, and at most 18 min to perform on the *C. elegans* dataset. The slowest correction tool was once again NaS, requiring up to 9 days to perform on the *S. cerevisiae* dataset. Comparatively, the slowest self-correction tool, ***Canu***, only required 4 h 37 min to correct the larger *C. elegans* dataset. In terms of memory consumption, self-correction tools were once again generally more demanding than hybrid correction methods, CoLoRMap and Hercules excluded, on the *A. baylyi*, *E. coli* and *S. cerevisiae* datasets. However, on the *C. elegans* dataset, a greater number of hybrid correction tools displayed higher memory requirements than self-correction tools. ***CONSENT***, the most memory-consuming self-correction tool after ***LoRMA***, required 14.5 GB, and was indeed comparable to Jabba, whereas other hybrid tools such as Proovread and HG-CoLoR displayed a memory peak of 20 GB.

These results suggest that, even when error rates reach less than 20%, self-correction methods cannot perform well on low coverage datasets. In fact, only ***MECAT*** and ***Daccord*** could provide satisfying correction on these datasets, but the latter was limited by its high-memory requirements, and could not scale to the larger *C. elegans* dataset, limiting its application. Hybrid correction thus remains more suited for such data, although it can be slower and more memory consuming. However, it is also worth noting that these hybrid correction methods tend to see their performances drop as the complexity and the number of repeats of the studied organisms grow. This is not the case for self-correction methods, and a tool such as ***MECAT*** could thus be preferred when studying high-complexity organisms. However, it outputs less corrected reads than most hybrid tools, and might thus not be suited for all types of downstream analyses, including assembly.

#### 3.5.2 Real data

As on the previous datasets of Section 3.5.1, almost all hybrid correction tools managed to reduce the error rate below 1% on this dataset. However, once again, LSC and Hercules displayed poor performances, reaching error rates of 8.8% for the former, and up to 14.2% for the latter. On this dataset however, NaS once again performed well. This can be explained by the fact that it was designed for and tested on ONT data, and is thus probably biased towards better ONT data correction. In terms of assembly, all tools but LSC, Jabba and ECTools, the latter not being able to yield any assembly, resulted in a single-contig assembly, covering a large proportion of the reference genome, the smallest being of 97.78% for LoRDEC.

Same conclusion as Section 3.5.1 also apply to self-correction tools on this dataset. However, here, even ***Daccord*** and ***MECAT*** did not manage to reduce the error rate of the reads below 1%. Excluding ***LoRMA***, which once again split the majority of the reads, the two best performing tools were ***Daccord***, reaching an error rate of 3.2%, and ***CONSENT***, reaching an error rate of 5.7%. Interestingly, in terms of assembly, the vast majority of self-correction tools did not manage to yield a single-contig assembly, and ***FLAS*** assembly even reached a total of 19 contigs. Only ***Canu*** and ***CONSENT*** managed to yield satisfying, single-contig assemblies, covering the whole reference genome. Despite being composed of a greater number of contigs, it is however worth noting that other assemblies still covered the reference genome well.

In terms of runtime, all hybrid correction tools but LoRDEC, Jabba and ECTools once again performed slower than self-correction methods, although, as previously stated, ECTools provided unsatisfying results, especially in terms of assembly. On this dataset, the fastest tool was Jabba, requiring 2 min, followed by ***MECAT***, which required 4 min. This underlines the fact that unlike on PacBio data, ***MECAT*** tends to be slightly slower when correcting ONT reads. Conversely, the slowest correction tool was NaS, requiring 3 days to perform correction, while ***Canu***, the slowest self-correction tool only required 35 min. In terms of memory consumption, both hybrid and self-correction tools were comparable, although hybrid tools such as CoLoRMap, and Proovread, and self-correction tools such as ***Daccord*** and ***LoRMA*** displayed higher memory requirements, the latter once again being the most memory-consuming tool, requiring a whole 32 GB.

These results further confirm conclusions drawn in Section 3.5.1, and once again suggest that self-correction methods cannot perform well on low coverage datasets, when error rates reach around 20%. Moreover, on this real ONT dataset, ***MECAT*** did not perform as good as on the simulated PacBio datasets of Section 3.5.1, both in terms of reduction of the error rate and in terms of quality of the assembly. These results thus reinforce the previous conclusions, confirming ***MECAT*** is not best suited for downstream analyses such as assembly, but also showing ***MECAT*** seems to be less efficient when dealing with ONT data. As a result, hybrid correction once again proves to be more efficient for handling such data, although it can be slower and more memory consuming.

### 3.6 Medium error rate, medium coverage

Here, we present results on the *A. baylyi* 60x (medium), *E. coli* 60x (medium), *S. cerevisiae* 60x (medium) and *C. elegans* 60x (medium) datasets of Table 3. For these datasets, when possible, we evaluated the split corrected long reads. When split reads were not available, we evaluated the trimmed corrected long reads. Finally, when both split and trimmed reads were not available, we evaluated the complete corrected long reads. These results are summarized in Table 8 and commented below. Moreover, for these experiments, we excluded the following tools from the comparison, for the ensuing reasons:

- ECTools, because of its unsatisfying results on previous experiments;
- Hercules, because of its unsatisfying results and large runtimes on previous experiments;
- LSC, because of its unsatisfying results on previous experiments;
- Nanocorr, because of its large runtimes on previous experiments;
- NaS, because of its large runtimes on previous experiments;

On these datasets, as in Section 3.5.1, all assessed hybrid correction tools managed to reduce the error rates to around 1% or less, on the *A. baylyi*, *E. coli* and *S. cerevisiae* datasets, and hybrid methods that were ran of the *C. elegans* dataset displayed performance drops, especially FMLRC, which reached an error rate of 3.42%.

However, for self-correction, unlike in Section 3.5.1, most tools did manage to reduce the error rates to around 1% on all four datasets. Actually, only ***CONSENT*** reached an error rate of more than 1% on all datasets, and only ***CONSENT*** and ***FLAS*** reached an error rate of more than 2% on the *C. elegans* dataset. Once again, ***Daccord*** performed best on the first three datasets, but could not be run on *C. elegans* due to its memory requirements, and ***LoRMA*** reported a large proportion of split reads. Despite this fact, most self-correction methods nonetheless reported less corrected bases than hybrid methods, and ***LoRMA*** and ***MECAT*** especially performed poorly regarding this metric. Moreover, it is worth noting that, compared to the results of Section 3.5.1, for which the error rates were the same, self-correction methods performed better on these datasets, for which coverage was higher. This underlines the fact that, with sufficient coverage, self-correction methods are well suited for the processing of moderately high error rates datasets, especially given the fact that, apart from ***FLAS*** and ***LoRMA***, they do not suffer from performance drops when applied to complex and repeat-rich genomes.

In terms of runtime, even though the slowest methods have been excluded, all hybrid correction tools except Jabba once again performed slower than self-correction methods. On all datasets, the fastest tool was once again ***MECAT***, requiring 2 min to perform on the *A. baylyi* dataset, and at most 1 h 37 min to perform on the *C. elegans* dataset. Conversely, the slowest tool was Proovread, requiring almost 18 hours to perform on the *S. cerevisiae* dataset. In comparison, ***LoRMA***, which was the slowest self-correction tool on this dataset, only required a little less than 2 hours to perform correction. In terms of memory consumption, self-correction tools were once again more demanding than most hybrid correction tools on all datasets, CoLoRMap and Proovread excluded. However, it is worth noting that hybrid correction tool Jabba displayed an important peak in memory consumption on the *C. elegans* dataset, requiring more than 13 GB, while it only required around 1 GB on the three other datasets. On this dataset, Jabba was thus comparable to or more demanding than most self-correction tools.

These results suggest that, when error rates reach 20% or less, self-correction methods can perform well as long as the coverage is sequencing depth is sufficiently high, and that a sequencing depth of around 60x is sufficient. In terms of quality of the results, self-correction methods are comparable to hybrid correction methods on such datasets. Moreover, these results also show that self-correction methods usually perform faster than hybrid correction methods, although they require larger amounts of memory. These observations are particularly interesting, given the additional fact that, unlike hybrid correction, self-correction methods do not see their performances drop as the complexity of the studied organisms grows. As a result, although hybrid correction remains efficient, self-correction is also well suited for such data, and can even be preferred when studying complex organisms, or when fast processing is needed.

### 3.7 Low error rate, low coverage

Here, we present results on the *E. coli* 30x, *S. cerevisiae* 30x and *C. elegans* 30x datasets of Table 3. For these datasets, when possible, we evaluated the trimmed corrected long reads. When trimmed reads were not available, we evaluated the complete corrected long reads. These results are summarized in Table 9 and commented below. Moreover, for these experiments, we excluded the following tools from the comparison, for the ensuing reasons:

- ECTools, because of its unsatisfying results on previous experiments;
- Hercules, because of its unsatisfying results and large runtimes on previous experiments;
- LSC, because of its unsatisfying results on previous experiments;
- Nanocorr, because of its large runtimes on previous experiments;
- NaS, because of its large runtimes on previous experiments;

On these datasets, both hybrid and self-correction methods, except ***LoRMA***, did manage to reduce the error rate to around 1% or less, on all datasets. More precisely, only the hybrid methods FMLRC, HG-CoLoR and LoRDEC reached an error rate of more than 1% on the *C. elegans* dataset. In comparison, on this dataset, all self-correction tools but ***LoRMA*** managed to reduce the error rate below 1%. This observation is in accordance with that previously made in Section 3.5.1 and Section 3.6, indicating that self-correction methods usually perform better on complex, repeat-rich organisms, compared to hybrid correction methods, which usually suffer from performance drops. Moreover, it is interesting to note that, on these datasets, self-correction tools such as ***FLAS***, ***CONSENT*** and ***Daccord*** reported as many corrected bases as hybrid methods, although ***LoRMA*** and ***MECAT*** still performed poorly regarding this metric. This is especially interesting, since it shows self-correction methods can not only provide high-quality correction on such datasets, but also manage to correct a large number of reads. Overall, hybrid correction and self-correction methods were thus highly comparable on these three datasets, in terms of quality of the results. This underlines the fact that, even with low coverage, self-correction methods do manage to perform well on datasets for which the error rate reaches around 12%.

**Table 7:**
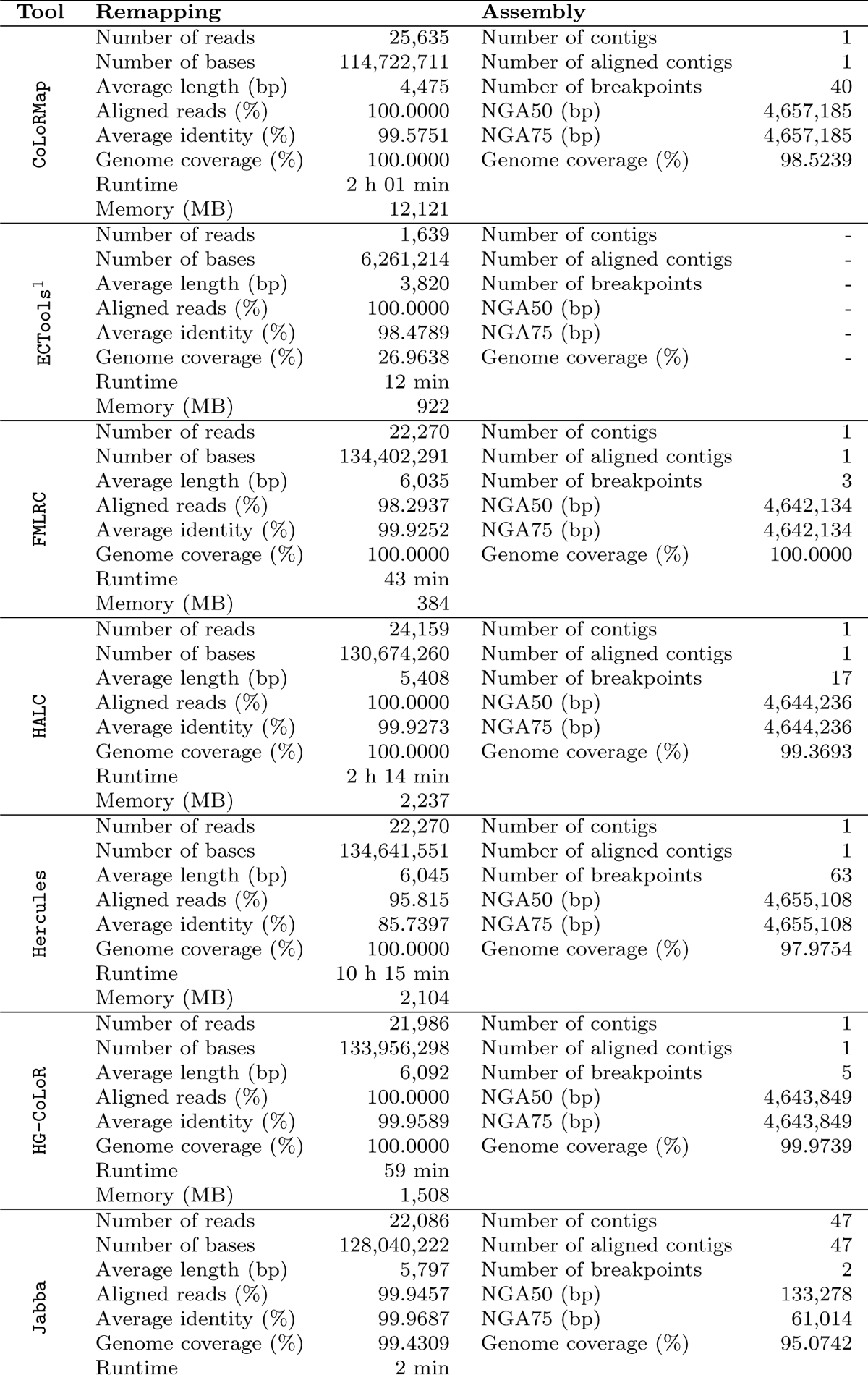
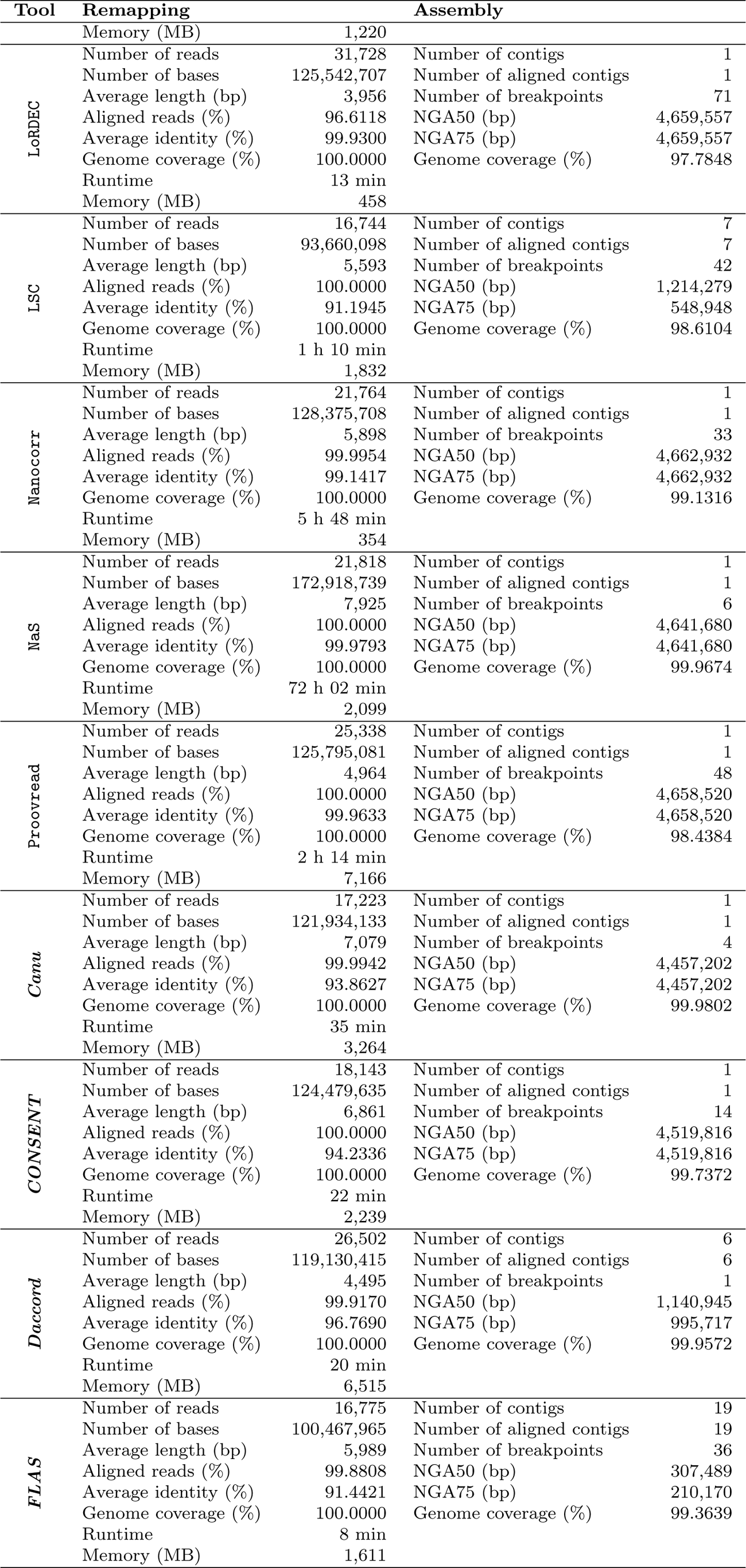
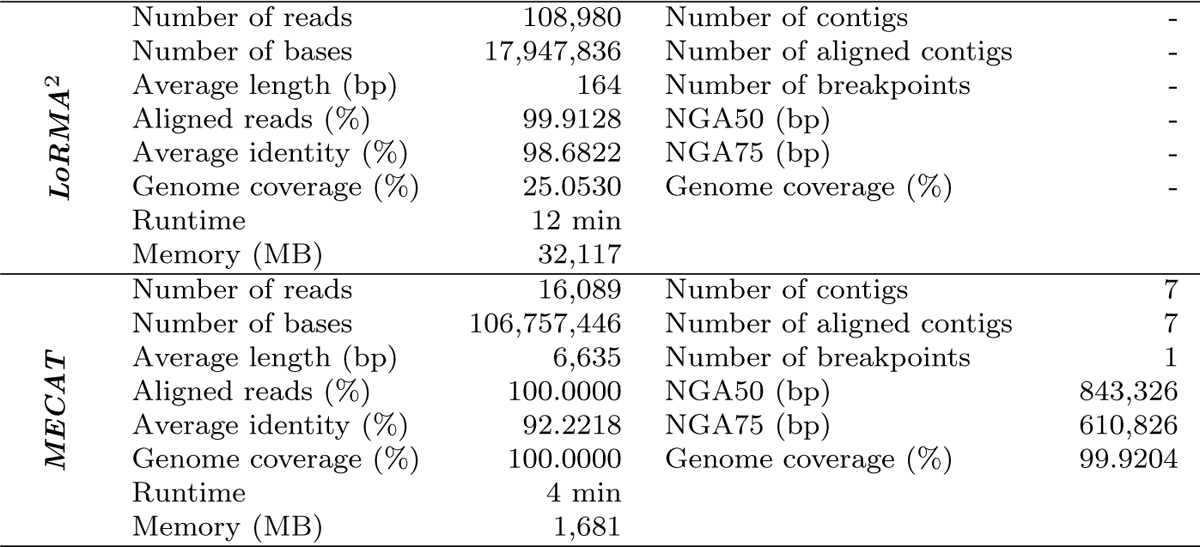
Comparison of the different error correction tools, on a medium error rate and low coverage real dataset. This corresponds to the *E. coli* real dataset of Table 3. ^1^ ECTools corrected reads could not be assembled. ^2^ ***LoRMA*** corrected reads could not be assembled.

**Table 8:**
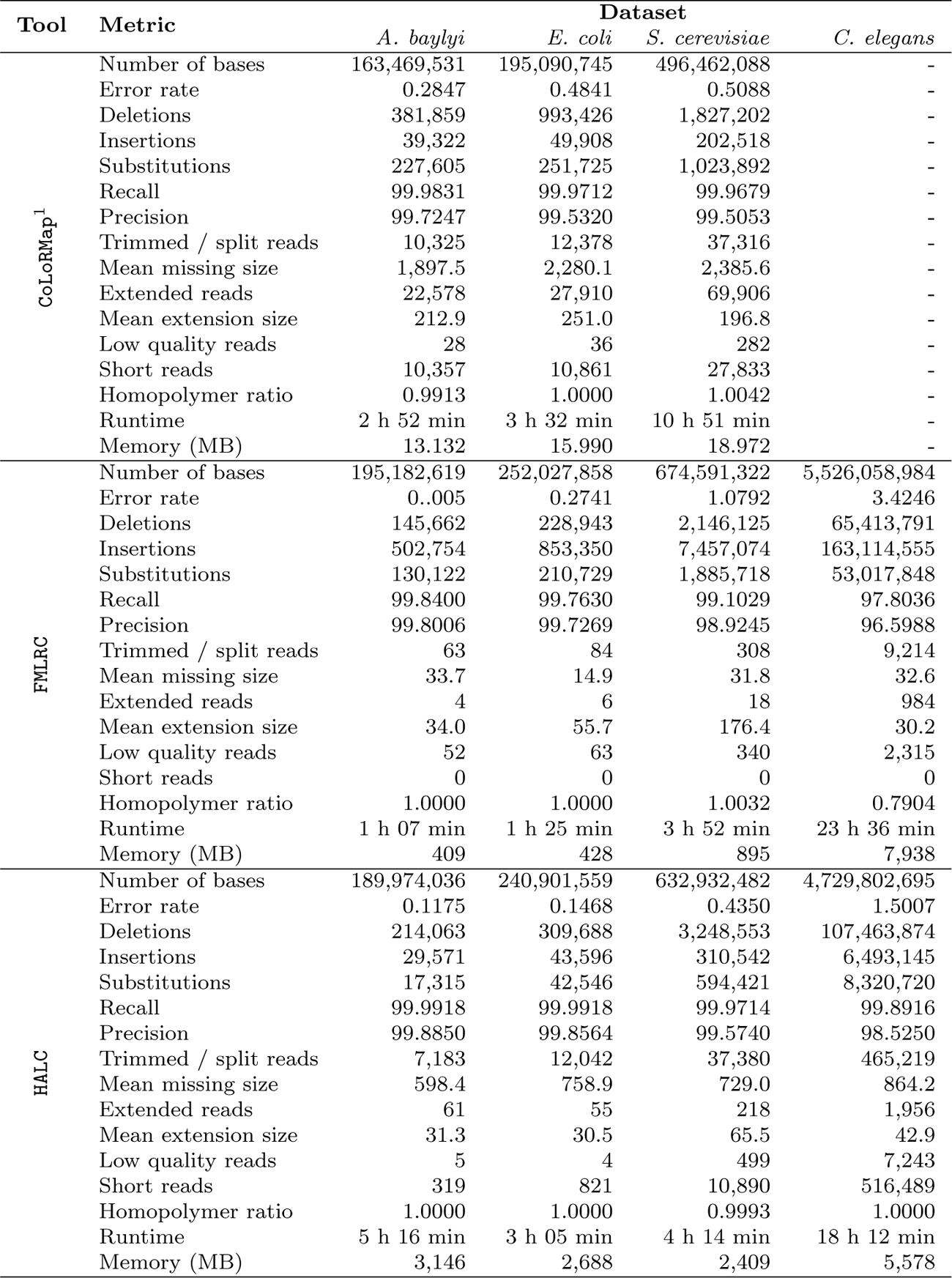
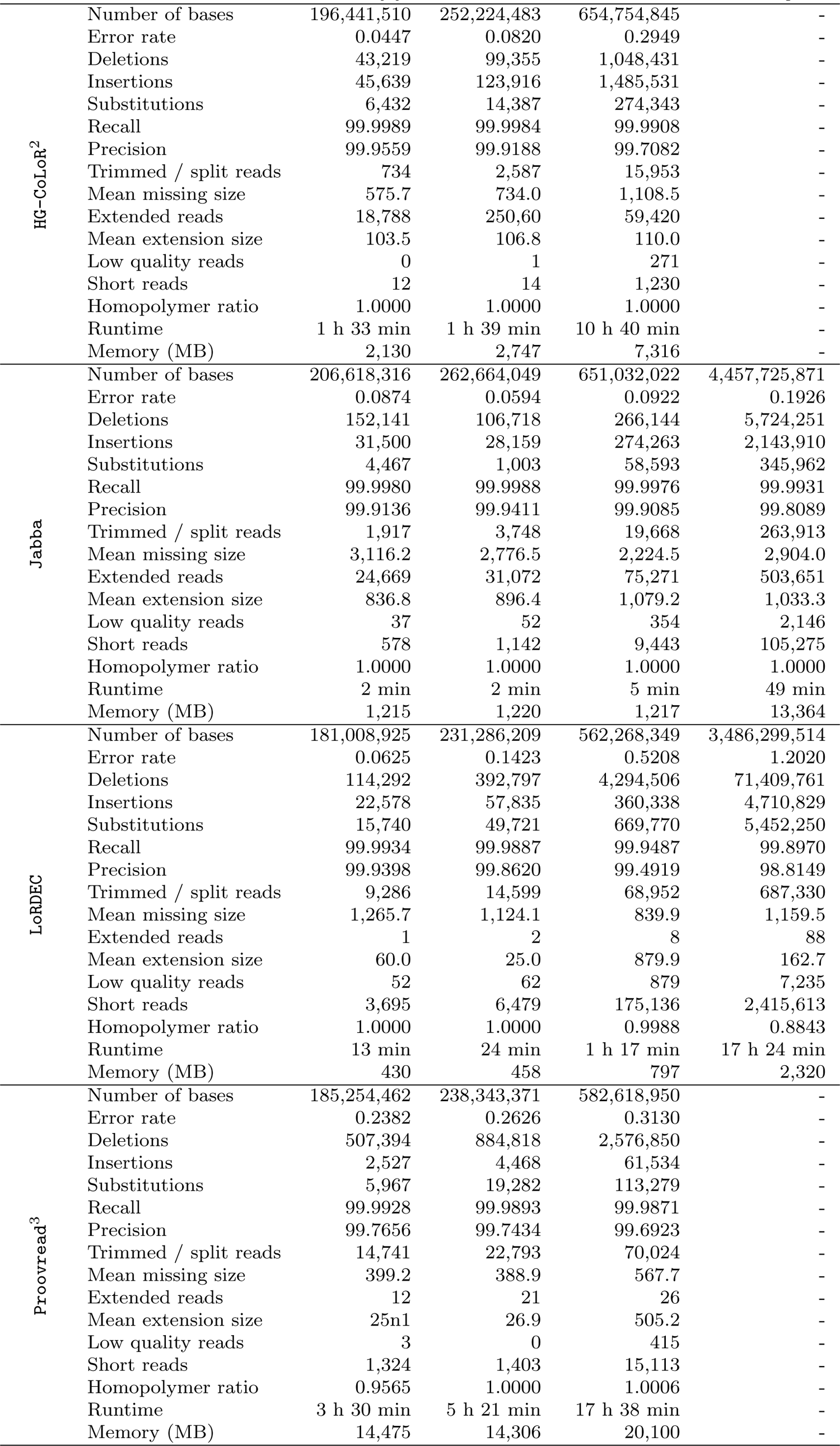
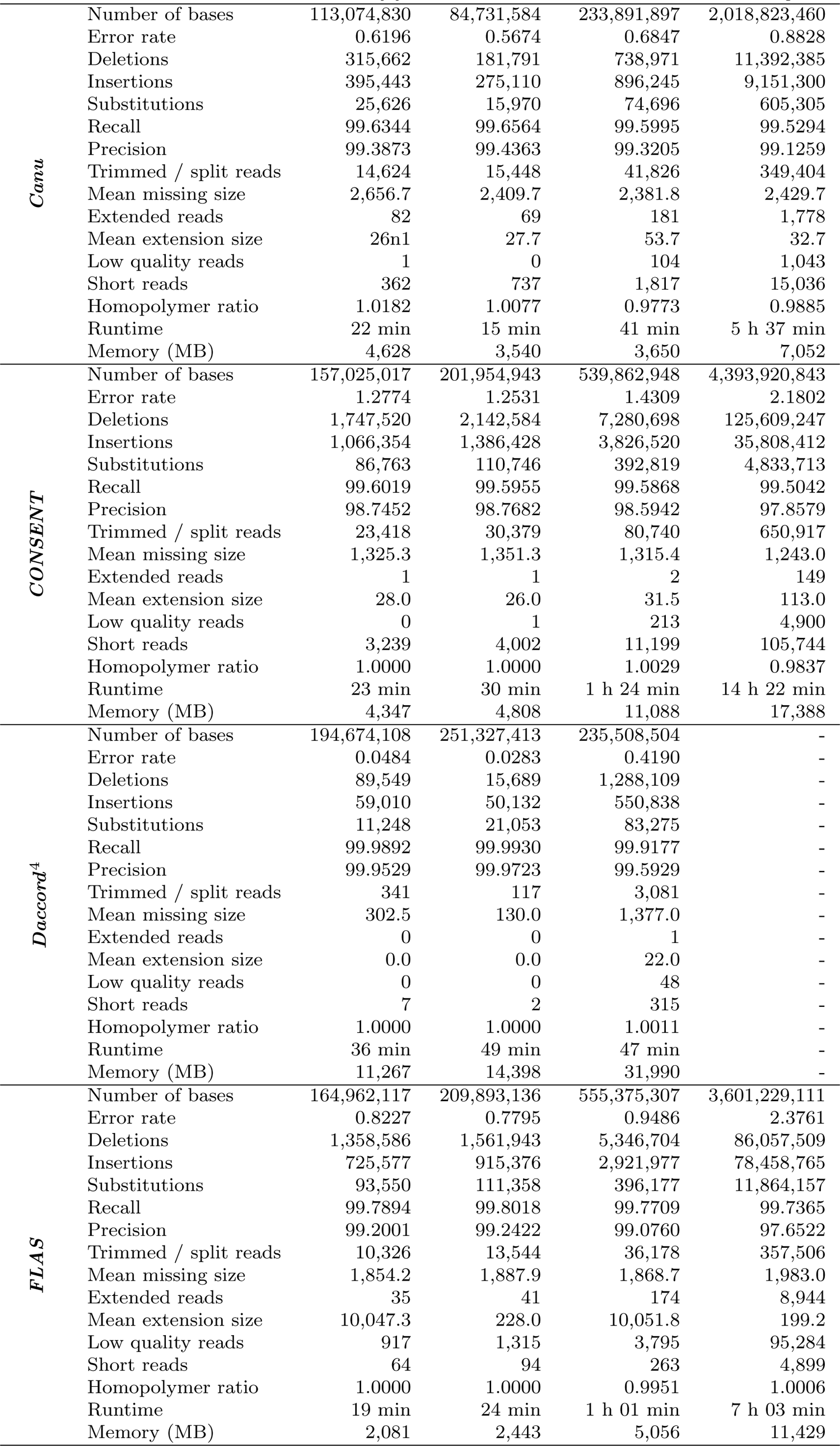
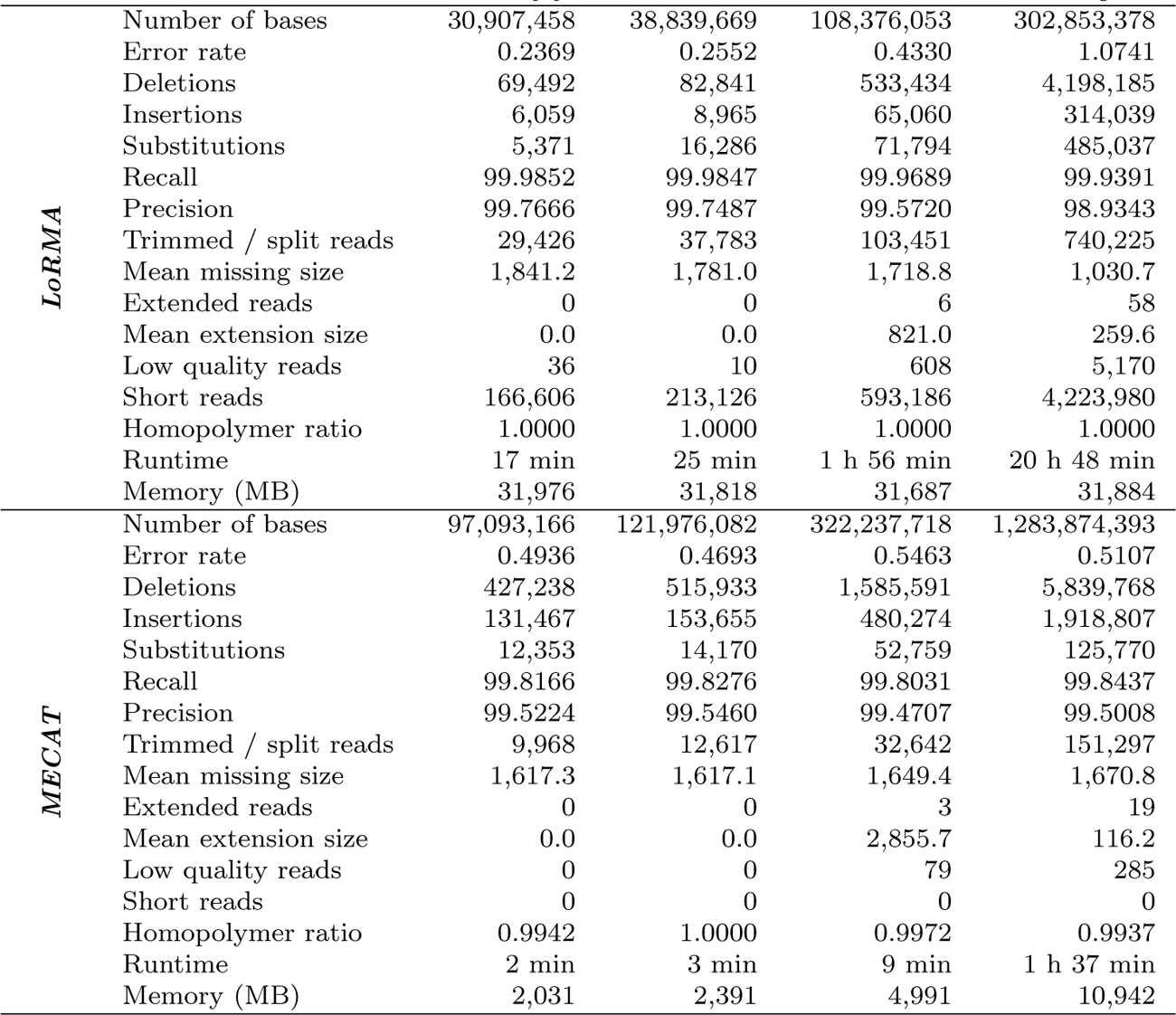
Comparison of the different error correction tools, on medium error rate and medium coverage datasets. This corresponds to the *A. baylyi* 60x (medium), *E. coli* 60x (medium), *S. cerevisiae* 60x (medium) and *C. elegans* 60x (medium) datasets of Table 3. ^1^ CoLoRMap was not launched on the *C. elegans* dataset due to its large runtimes. ^2^ HG-CoLoR was not launched on the *C. elegans* dataset due to its large runtimes. ^3^ Proovread was not launched on the *C. elegans* dataset due to its large runtimes. ^4^ ***Daccord*** could not perform correction on the *C. elegans* dataset, since the alignment step with DALIGNER required more than 128 GB of memory.

| Tool | Metric | Dataset |  |  |  |
| --- | --- | --- | --- | --- | --- |
|  |  | *A. baylyi* | *E. coli* | *S. cerevisiae* | *C. elegans* |
| CoLoRMap<sup>1</sup> | Number of bases | 163,469,531 | 195,090,745 | 496,462,088 | - |
|  | Error rate | 0.2847 | 0.4841 | 0.5088 | - |
|  | Deletions | 381,859 | 993,426 | 1,827,202 | - |
|  | Insertions | 39,322 | 49,908 | 202,518 | - |
|  | Substitutions | 227,605 | 251,725 | 1,023,892 | - |
|  | Recall | 99.9831 | 99.9712 | 99.9679 | - |
|  | Precision | 99.7247 | 99.5320 | 99.5053 | - |
|  | Trimmed / split reads | 10,325 | 12,378 | 37,316 | - |
|  | Mean missing size | 1,897.5 | 2,280.1 | 2,385.6 | - |
|  | Extended reads | 22,578 | 27,910 | 69,906 | - |
|  | Mean extension size | 212.9 | 251.0 | 196.8 | - |
|  | Low quality reads | 28 | 36 | 282 | - |
|  | Short reads | 10,357 | 10,861 | 27,833 | - |
|  | Homopolymer ratio | 0.9913 | 1.0000 | 1.0042 | - |
|  | Runtime | 2 h 52 min | 3 h 32 min | 10 h 51 min | - |
|  | Memory (MB) | 13.132 | 15.990 | 18.972 | - |
| FMLRC | Number of bases | 195,182,619 | 252,027,858 | 674,591,322 | 5,526,058,984 |
|  | Error rate | 0.005 | 0.2741 | 1.0792 | 3.4246 |
|  | Deletions | 145,662 | 228,943 | 2,146,125 | 65,413,791 |
|  | Insertions | 502,754 | 853,350 | 7,457,074 | 163,114,555 |
|  | Substitutions | 130,122 | 210,729 | 1,885,718 | 53,017,848 |
|  | Recall | 99.8400 | 99.7630 | 99.1029 | 97.8036 |
|  | Precision | 99.8006 | 99.7269 | 98.9245 | 96.5988 |
|  | Trimmed / split reads | 63 | 84 | 308 | 9,214 |
|  | Mean missing size | 33.7 | 14.9 | 31.8 | 32.6 |
|  | Extended reads | 4 | 6 | 18 | 984 |
|  | Mean extension size | 34.0 | 55.7 | 176.4 | 30.2 |
|  | Low quality reads | 52 | 63 | 340 | 2,315 |
|  | Short reads | 0 | 0 | 0 | 0 |
|  | Homopolymer ratio | 1.0000 | 1.0000 | 1.0032 | 0.7904 |
|  | Runtime | 1 h 07 min | 1 h 25 min | 3 h 52 min | 23 h 36 min |
|  | Memory (MB) | 409 | 428 | 895 | 7,938 |
| HALC | Number of bases | 189,974,036 | 240,901,559 | 632,932,482 | 4,729,802,695 |
|  | Error rate | 0.1175 | 0.1468 | 0.4350 | 1.5007 |
|  | Deletions | 214,063 | 309,688 | 3,248,553 | 107,463,874 |
|  | Insertions | 29,571 | 43,596 | 310,542 | 6,493,145 |
|  | Substitutions | 17,315 | 42,546 | 594,421 | 8,320,720 |
|  | Recall | 99.9918 | 99.9918 | 99.9714 | 99.8916 |
|  | Precision | 99.8850 | 99.8564 | 99.5740 | 98.5250 |
|  | Trimmed / split reads | 7,183 | 12,042 | 37,380 | 465,219 |
|  | Mean missing size | 598.4 | 758.9 | 729.0 | 864.2 |
|  | Extended reads | 61 | 55 | 218 | 1,956 |
|  | Mean extension size | 31.3 | 30.5 | 65.5 | 42.9 |
|  | Low quality reads | 5 | 4 | 499 | 7,243 |
|  | Short reads | 319 | 821 | 10,890 | 516,489 |
|  | Homopolymer ratio | 1.0000 | 1.0000 | 0.9993 | 1.0000 |
|  | Runtime | 5 h 16 min | 3 h 05 min | 4 h 14 min | 18 h 12 min |
|  | Memory (MB) | 3,146 | 2,688 | 2,409 | 5,578 |
|  |  | *A. baylyi* | *E. coli* | *S. cerevisiae* | *C. elegans* |
| HG-CoLoR<sup>2</sup> | Number of bases | 196,441,510 | 252,224,483 | 654,754,845 | - |
|  | Error rate | 0.0447 | 0.0820 | 0.2949 | - |
|  | Deletions | 43,219 | 99,355 | 1,048,431 | - |
|  | Insertions | 45,639 | 123,916 | 1,485,531 | - |
|  | Substitutions | 6,432 | 14,387 | 274,343 | - |
|  | Recall | 99.9989 | 99.9984 | 99.9908 | - |
|  | Precision | 99.9559 | 99.9188 | 99.7082 | - |
|  | Trimmed / split reads | 734 | 2,587 | 15,953 | - |
|  | Mean missing size | 575.7 | 734.0 | 1,108.5 | - |
|  | Extended reads | 18,788 | 250,60 | 59,420 | - |
|  | Mean extension size | 103.5 | 106.8 | 110.0 | - |
|  | Low quality reads | 0 | 1 | 271 | - |
|  | Short reads | 12 | 14 | 1,230 | - |
|  | Homopolymer ratio | 1.0000 | 1.0000 | 1.0000 | - |
|  | Runtime | 1 h 33 min | 1 h 39 min | 10 h 40 min | - |
|  | Memory (MB) | 2,130 | 2,747 | 7,316 | - |
| Jabba | Number of bases | 206,618,316 | 262,664,049 | 651,032,022 | 4,457,725,871 |
|  | Error rate | 0.0874 | 0.0594 | 0.0922 | 0.1926 |
|  | Deletions | 152,141 | 106,718 | 266,144 | 5,724,251 |
|  | Insertions | 31,500 | 28,159 | 274,263 | 2,143,910 |
|  | Substitutions | 4,467 | 1,003 | 58,593 | 345,962 |
|  | Recall | 99.9980 | 99.9988 | 99.9976 | 99.9931 |
|  | Precision | 99.9136 | 99.9411 | 99.9085 | 99.8089 |
|  | Trimmed / split reads | 1,917 | 3,748 | 19,668 | 263,913 |
|  | Mean missing size | 3,116.2 | 2,776.5 | 2,224.5 | 2,904.0 |
|  | Extended reads | 24,669 | 31,072 | 75,271 | 503,651 |
|  | Mean extension size | 836.8 | 896.4 | 1,079.2 | 1,033.3 |
|  | Low quality reads | 37 | 52 | 354 | 2,146 |
|  | Short reads | 578 | 1,142 | 9,443 | 105,275 |
|  | Homopolymer ratio | 1.0000 | 1.0000 | 1.0000 | 1.0000 |
|  | Runtime | 2 min | 2 min | 5 min | 49 min |
|  | Memory (MB) | 1,215 | 1,220 | 1,217 | 13,364 |
| LoRDEC | Number of bases | 181,008,925 | 231,286,209 | 562,268,349 | 3,486,299,514 |
|  | Error rate | 0.0625 | 0.1423 | 0.5208 | 1.2020 |
|  | Deletions | 114,292 | 392,797 | 4,294,506 | 71,409,761 |
|  | Insertions | 22,578 | 57,835 | 360,338 | 4,710,829 |
|  | Substitutions | 15,740 | 49,721 | 669,770 | 5,452,250 |
|  | Recall | 99.9934 | 99.9887 | 99.9487 | 99.8970 |
|  | Precision | 99.9398 | 99.8620 | 99.4919 | 98.8149 |
|  | Trimmed / split reads | 9,286 | 14,599 | 68,952 | 687,330 |
|  | Mean missing size | 1,265.7 | 1,124.1 | 839.9 | 1,159.5 |
|  | Extended reads | 1 | 2 | 8 | 88 |
|  | Mean extension size | 60.0 | 25.0 | 879.9 | 162.7 |
|  | Low quality reads | 52 | 62 | 879 | 7,235 |
|  | Short reads | 3,695 | 6,479 | 175,136 | 2,415,613 |
|  | Homopolymer ratio | 1.0000 | 1.0000 | 0.9988 | 0.8843 |
|  | Runtime | 13 min | 24 min | 1 h 17 min | 17 h 24 min |
|  | Memory (MB) | 430 | 458 | 797 | 2,320 |
| Proovread<sup>3</sup> | Number of bases | 185,254,462 | 238,343,371 | 582,618,950 | - |
|  | Error rate | 0.2382 | 0.2626 | 0.3130 | - |
|  | Deletions | 507,394 | 884,818 | 2,576,850 | - |
|  | Insertions | 2,527 | 4,468 | 61,534 | - |
|  | Substitutions | 5,967 | 19,282 | 113,279 | - |
|  | Recall | 99.9928 | 99.9893 | 99.9871 | - |
|  | Precision | 99.7656 | 99.7434 | 99.6923 | - |
|  | Trimmed / split reads | 14,741 | 22,793 | 70,024 | - |
|  | Mean missing size | 399.2 | 388.9 | 567.7 | - |
|  | Extended reads | 12 | 21 | 26 | - |
|  | Mean extension size | 25n1 | 26.9 | 505.2 | - |
|  | Low quality reads | 3 | 0 | 415 | - |
|  | Short reads | 1,324 | 1,403 | 15,113 | - |
|  | Homopolymer ratio | 0.9565 | 1.0000 | 1.0006 | - |
|  | Runtime | 3 h 30 min | 5 h 21 min | 17 h 38 min | - |
|  | Memory (MB) | 14,475 | 14,306 | 20,100 | - |
|  |  | *A. baylyi* | *E. coli* | *S. cerevisiae* | *C. elegans* |
| *Canu* | Number of bases | 113,074,830 | 84,731,584 | 233,891,897 | 2,018,823,460 |
|  | Error rate | 0.6196 | 0.5674 | 0.6847 | 0.8828 |
|  | Deletions | 315,662 | 181,791 | 738,971 | 11,392,385 |
|  | Insertions | 395,443 | 275,110 | 896,245 | 9,151,300 |
|  | Substitutions | 25,626 | 15,970 | 74,696 | 605,305 |
|  | Recall | 99.6344 | 99.6564 | 99.5995 | 99.5294 |
|  | Precision | 99.3873 | 99.4363 | 99.3205 | 99.1259 |
|  | Trimmed / split reads | 14,624 | 15,448 | 41,826 | 349,404 |
|  | Mean missing size | 2,656.7 | 2,409.7 | 2,381.8 | 2,429.7 |
|  | Extended reads | 82 | 69 | 181 | 1,778 |
|  | Mean extension size | 26n1 | 27.7 | 53.7 | 32.7 |
|  | Low quality reads | 1 | 0 | 104 | 1,043 |
|  | Short reads | 362 | 737 | 1,817 | 15,036 |
|  | Homopolymer ratio | 1.0182 | 1.0077 | 0.9773 | 0.9885 |
|  | Runtime | 22 min | 15 min | 41 min | 5 h 37 min |
|  | Memory (MB) | 4,628 | 3,540 | 3,650 | 7,052 |
| *CONSENT* | Number of bases | 157,025,017 | 201,954,943 | 539,862,948 | 4,393,920,843 |
|  | Error rate | 1.2774 | 1.2531 | 1.4309 | 2.1802 |
|  | Deletions | 1,747,520 | 2,142,584 | 7,280,698 | 125,609,247 |
|  | Insertions | 1,066,354 | 1,386,428 | 3,826,520 | 35,808,412 |
|  | Substitutions | 86,763 | 110,746 | 392,819 | 4,833,713 |
|  | Recall | 99.6019 | 99.5955 | 99.5868 | 99.5042 |
|  | Precision | 98.7452 | 98.7682 | 98.5942 | 97.8579 |
|  | Trimmed / split reads | 23,418 | 30,379 | 80,740 | 650,917 |
|  | Mean missing size | 1,325.3 | 1,351.3 | 1,315.4 | 1,243.0 |
|  | Extended reads | 1 | 1 | 2 | 149 |
|  | Mean extension size | 28.0 | 26.0 | 31.5 | 113.0 |
|  | Low quality reads | 0 | 1 | 213 | 4,900 |
|  | Short reads | 3,239 | 4,002 | 11,199 | 105,744 |
|  | Homopolymer ratio | 1.0000 | 1.0000 | 1.0029 | 0.9837 |
|  | Runtime | 23 min | 30 min | 1 h 24 min | 14 h 22 min |
|  | Memory (MB) | 4,347 | 4,808 | 11,088 | 17,388 |
| *Daccord*<sup>4</sup> | Number of bases | 194,674,108 | 251,327,413 | 235,508,504 | - |
|  | Error rate | 0.0484 | 0.0283 | 0.4190 | - |
|  | Deletions | 89,549 | 15,689 | 1,288,109 | - |
|  | Insertions | 59,010 | 50,132 | 550,838 | - |
|  | Substitutions | 11,248 | 21,053 | 83,275 | - |
|  | Recall | 99.9892 | 99.9930 | 99.9177 | - |
|  | Precision | 99.9529 | 99.9723 | 99.5929 | - |
|  | Trimmed / split reads | 341 | 117 | 3,081 | - |
|  | Mean missing size | 302.5 | 130.0 | 1,377.0 | - |
|  | Extended reads | 0 | 0 | 1 | - |
|  | Mean extension size | 0.0 | 0.0 | 22.0 | - |
|  | Low quality reads | 0 | 0 | 48 | - |
|  | Short reads | 7 | 2 | 315 | - |
|  | Homopolymer ratio | 1.0000 | 1.0000 | 1.0011 | - |
|  | Runtime | 36 min | 49 min | 47 min | - |
|  | Memory (MB) | 11,267 | 14,398 | 31,990 | - |
| *FLAS* | Number of bases | 164,962,117 | 209,893,136 | 555,375,307 | 3,601,229,111 |
|  | Error rate | 0.8227 | 0.7795 | 0.9486 | 2.3761 |
|  | Deletions | 1,358,586 | 1,561,943 | 5,346,704 | 86,057,509 |
|  | Insertions | 725,577 | 915,376 | 2,921,977 | 78,458,765 |
|  | Substitutions | 93,550 | 111,358 | 396,177 | 11,864,157 |
|  | Recall | 99.7894 | 99.8018 | 99.7709 | 99.7365 |
|  | Precision | 99.2001 | 99.2422 | 99.0760 | 97.6522 |
|  | Trimmed / split reads | 10,326 | 13,544 | 36,178 | 357,506 |
|  | Mean missing size | 1,854.2 | 1,887.9 | 1,868.7 | 1,983.0 |
|  | Extended reads | 35 | 41 | 174 | 8,944 |
|  | Mean extension size | 10,047.3 | 228.0 | 10,051.8 | 199.2 |
|  | Low quality reads | 917 | 1,315 | 3,795 | 95,284 |
|  | Short reads | 64 | 94 | 263 | 4,899 |
|  | Homopolymer ratio | 1.0000 | 1.0000 | 0.9951 | 1.0006 |
|  | Runtime | 19 min | 24 min | 1 h 01 min | 7 h 03 min |
|  | Memory (MB) | 2,081 | 2,443 | 5,056 | 11,429 |
|  |  | *A. baylyi* | *E. coli* | *S. cerevisiae* | *C. elegans* |
| **LoRMA** | Number of bases | 30,907,458 | 38,839,669 | 108,376,053 | 302,853,378 |
|  | Error rate | 0.2369 | 0.2552 | 0.4330 | 1.0741 |
|  | Deletions | 69,492 | 82,841 | 533,434 | 4,198,185 |
|  | Insertions | 6,059 | 8,965 | 65,060 | 314,039 |
|  | Substitutions | 5,371 | 16,286 | 71,794 | 485,037 |
|  | Recall | 99.9852 | 99.9847 | 99.9689 | 99.9391 |
|  | Precision | 99.7666 | 99.7487 | 99.5720 | 98.9343 |
|  | Trimmed / split reads | 29,426 | 37,783 | 103,451 | 740,225 |
|  | Mean missing size | 1,841.2 | 1,781.0 | 1,718.8 | 1,030.7 |
|  | Extended reads | 0 | 0 | 6 | 58 |
|  | Mean extension size | 0.0 | 0.0 | 821.0 | 259.6 |
|  | Low quality reads | 36 | 10 | 608 | 5,170 |
|  | Short reads | 166,606 | 213,126 | 593,186 | 4,223,980 |
|  | Homopolymer ratio | 1.0000 | 1.0000 | 1.0000 | 1.0000 |
|  | Runtime | 17 min | 25 min | 1 h 56 min | 20 h 48 min |
|  | Memory (MB) | 31,976 | 31,818 | 31,687 | 31,884 |
| **MECAT** | Number of bases | 97,093,166 | 121,976,082 | 322,237,718 | 1,283,874,393 |
|  | Error rate | 0.4936 | 0.4693 | 0.5463 | 0.5107 |
|  | Deletions | 427,238 | 515,933 | 1,585,591 | 5,839,768 |
|  | Insertions | 131,467 | 153,655 | 480,274 | 1,918,807 |
|  | Substitutions | 12,353 | 14,170 | 52,759 | 125,770 |
|  | Recall | 99.8166 | 99.8276 | 99.8031 | 99.8437 |
|  | Precision | 99.5224 | 99.5460 | 99.4707 | 99.5008 |
|  | Trimmed / split reads | 9,968 | 12,617 | 32,642 | 151,297 |
|  | Mean missing size | 1,617.3 | 1,617.1 | 1,649.4 | 1,670.8 |
|  | Extended reads | 0 | 0 | 3 | 19 |
|  | Mean extension size | 0.0 | 0.0 | 2,855.7 | 116.2 |
|  | Low quality reads | 0 | 0 | 79 | 285 |
|  | Short reads | 0 | 0 | 0 | 0 |
|  | Homopolymer ratio | 0.9942 | 1.0000 | 0.9972 | 0.9937 |
|  | Runtime | 2 min | 3 min | 9 min | 1 h 37 min |
|  | Memory (MB) | 2,031 | 2,391 | 4,991 | 10,942 |

**Table 9:**
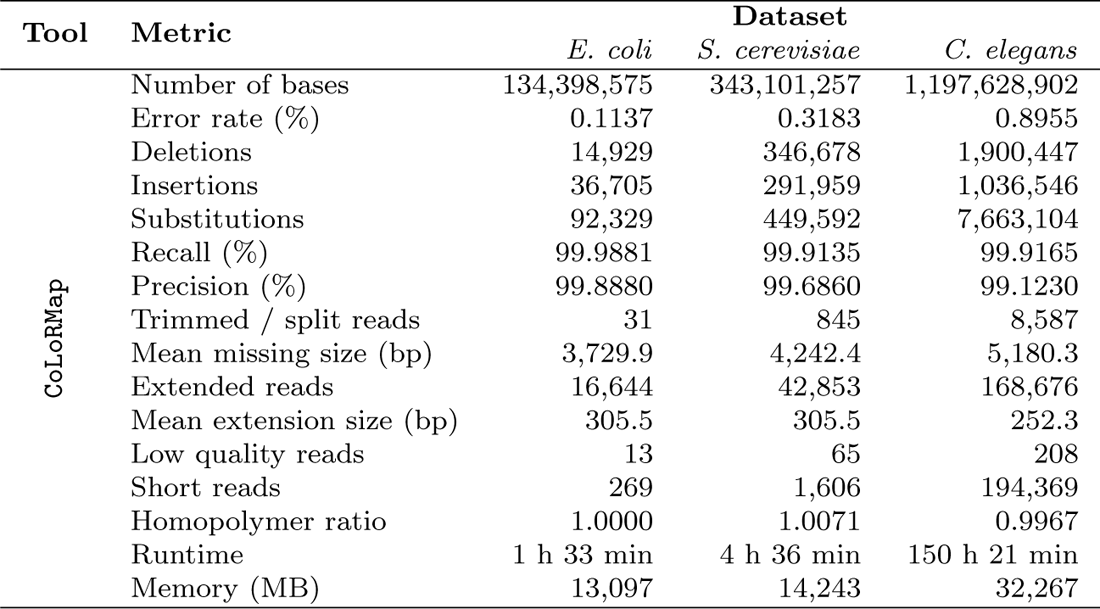
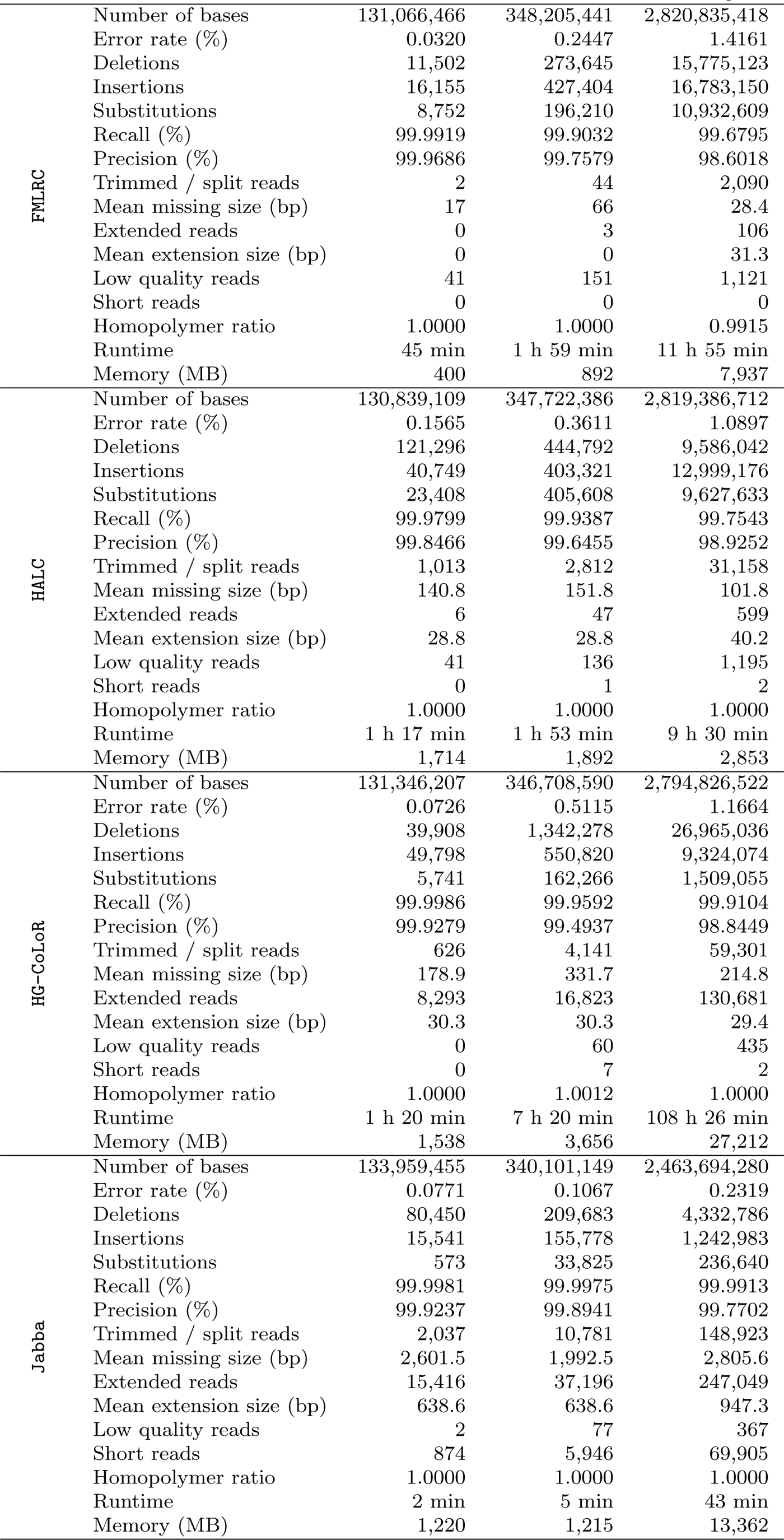
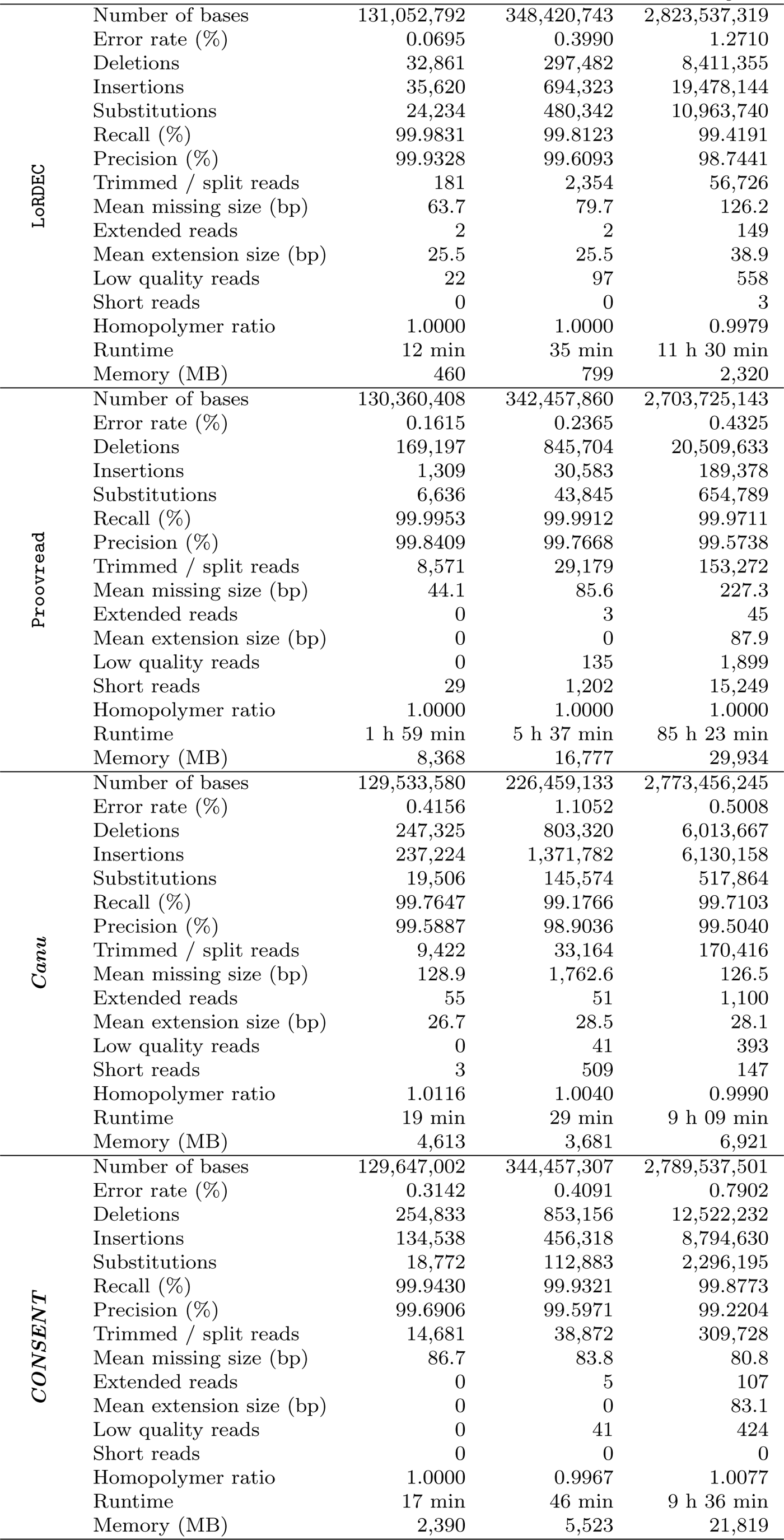
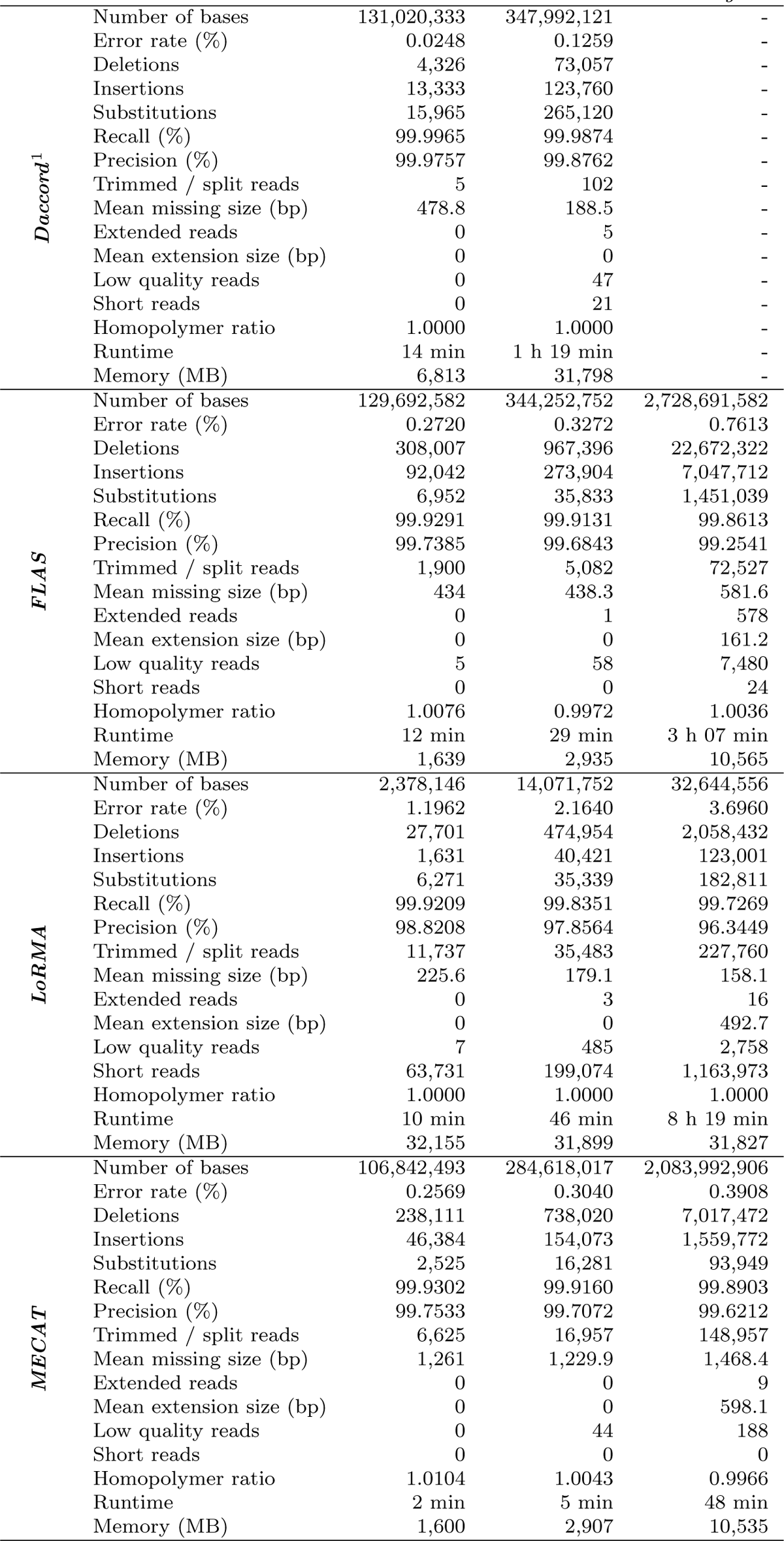
Comparison of the different error correction tools, on low error rate and low coverage datasets. This corresponds to the *E. coli* 30x, *S. cerevisiae* 30x and *C. elegans* 30x datasets of Table 3. ^1^ ***Daccord*** could not perform correction, since the alignment step with DALIGNER required more than 128 GB of memory.

**Table 10:** Comparison of the different error correction tools, on low error rate and medium coverage datasets. This corresponds to the *E. coli* 60x (low), *S. cerevisiae* 60x (low) and *C. elegans* 60x (low) datasets of Table 3. ^1^ CoLoRMap was not launched on the *C. elegans* dataset due to its large runtimes. ^2^ HG-CoLoR was not launched on the *C. elegans* dataset due to its large runtimes. ^3^ Proovread was not launched on the *C. elegans* dataset due to its large runtimes. ^4^ ***Daccord*** could not perform correction on the *C. elegans* dataset, since the alignment step with DALIGNER required more than 128 GB of memory.

| Tool | Metric | Dataset |  |  |
| --- | --- | --- | --- | --- |
|  |  | *E. coli* | *S. cerevisiae* | *C. elegans* |
| CoLoRMap<sup>1</sup> | Number of bases | 266,210,819 | 663,617,071 | - |
|  | Error rate (%) | 0.1621 | 0.6143 | - |
|  | Deletions | 98,293 | 1,930,344 | - |
|  | Insertions | 112,511 | 1,216,405 | - |
|  | Substitutions | 199,934 | 1,036,618 | - |
|  | Recall (%) | 99.9631 | 99.7755 | - |
|  | Precision (%) | 99.8400 | 99.3917 | - |
|  | Trimmed / split reads | 185 | 3,913 | - |
|  | Mean missing size (bp) | 3,787.2 | 3,746.2 | - |
|  | Extended reads | 33,047 | 82,438 | - |
|  | Mean extension size (bp) | 297.7 | 297.7 | - |
|  | Low quality reads | 33 | 145 | - |
|  | Short reads | 653 | 4,679 | - |
|  | Homopolymer ratio | 1.0089 | 0.9946 | - |
|  | Runtime | 3 h 01 min | 8 h 08 min | - |
|  | Memory (MB) | 19,898 | 24,375 | - |
| FMLRC | Number of bases | 261,387,632 | 695,239,831 | 5,652,356,620 |
|  | Error rate (%) | 0.0292 | 0.2469 | 1.4213 |
|  | Deletions | 24,753 | 551,321 | 31,671,968 |
|  | Insertions | 31,069 | 872,824 | 33,701,759 |
|  | Substitutions | 17,368 | 391,349 | 21,909,721 |
|  | Recall (%) | 99.9944 | 99.9006 | 99.6764 |
|  | Precision (%) | 99.9714 | 99.7557 | 98.5966 |
|  | Trimmed / split reads | 14 | 106 | 4,147 |
|  | Mean missing size (bp) | 17.7 | 48.5 | 28.9 |
|  | Extended reads | 0 | 8 | 242 |
|  | Mean extension size (bp) | 0 | 0 | 28.1 |
|  | Low quality reads | 85 | 360 | 2,315 |
|  | Short reads | 0 | 0 | 0 |
|  | Homopolymer ratio | 1.0000 | 1.0000 | 0.9856 |
|  | Runtime | 1 h 28 min | 3 h 57 min | 23 h 25 min |
|  | Memory (MB) | 403 | 893 | 7,937 |
| HALC | Number of bases | 260,933,848 | 694,333,491 | 5,649,091,260 |
|  | Error rate (%) | 0.1522 | 0.3648 | 1.0880 |
|  | Deletions | 233,633 | 883,877 | 19,179,388 |
|  | Insertions | 84,745 | 822,236 | 25,917,654 |
|  | Substitutions | 48,696 | 827,403 | 19,226,301 |
|  | Recall (%) | 99.9785 | 99.9354 | 99.7550 |
|  | Precision (%) | 99.8509 | 99.6420 | 98.9269 |
|  | Trimmed / split reads | 2,033 | 5,707 | 61,894 |
|  | Mean missing size (bp) | 147.3 | 131.6 | 102 |
|  | Extended reads | 11 | 60 | 1,284 |
|  | Mean extension size (bp) | 22.4 | 22.4 | 42.6 |
|  | Low quality reads | 79 | 329 | 2,512 |
|  | Short reads | 0 | 1 | 0 |
|  | Homopolymer ratio | 1.0000 | 1.0000 | 0.9852 |
|  | Runtime | 4 h 57 min | 4 h 25 min | 19 h 10 min |
|  | Memory (MB) | 2,747 | 2,487 | 5,716 |
| HG-CoLoR<sup>2</sup> | Number of bases | 261,879,085 | 690,178,663 | - |
|  | Error rate (%) | 0.0771 | 0.5995 | - |
|  | Deletions | 87,607 | 2,824,556 | - |
|  | Insertions | 113,291 | 1,924,555 | - |
|  | Substitutions | 14,055 | 374,714 | - |
|  | Recall (%) | 99.9987 | 99.9433 | - |
|  | Precision (%) | 99.9234 | 99.4059 | - |
|  | Trimmed / split reads | 1,569 | 9,700 | - |
|  | Mean missing size (bp) | 264.1 | 496.2 | - |
|  | Extended reads | 19,498 | 40,638 | - |
|  | Mean extension size (bp) | 33.2 | 33.2 | - |
|  | Low quality reads | 1 | 235 | - |
|  | Short reads | 0 | 45 | - |
|  | Homopolymer ratio | 1.0000 | 1.0000 | - |
|  | Runtime | 2 h 03 min | 12 h 23 min | - |
|  | Memory (MB) | 2,744 | 7,297 | - |
|  |  | *E. coli* | *S. cerevisiae* | *C. elegans* |
| Jabba | Number of bases | 267,665,882 | 678,936,284 | 4,934,623,442 |
|  | Error rate (%) | 0.0716 | 0.1040 | 0.2312 |
|  | Deletions | 146,631 | 387,538 | 8,496,067 |
|  | Insertions | 30,775 | 340,301 | 2,534,990 |
|  | Substitutions | 1,171 | 63,194 | 419,150 |
|  | Recall (%) | 99.9984 | 99.9975 | 99.9916 |
|  | Precision (%) | 99.9292 | 99.8967 | 99.7710 |
|  | Trimmed / split reads | 4,095 | 21,726 | 298,954 |
|  | Mean missing size (bp) | 2,507.2 | 1,996 | 2,804.2 |
|  | Extended reads | 30,786 | 74,343 | 493,302 |
|  | Mean extension size (bp) | 633.2 | 633.2 | 949.5 |
|  | Low quality reads | 1 | 183 | 731 |
|  | Short reads | 1,649 | 11,922 | 140,889 |
|  | Homopolymer ratio | 1.0000 | 1.0000 | 1.0000 |
|  | Runtime | 2 min | 5 min | 49 min |
|  | Memory (MB) | 1,220 | 1,215 | 13,360 |
| LoRDEC | Number of bases | 261,360,235 | 695,578,111 | 5,657,161,182 |
|  | Error rate (%) | 0.0684 | 0.3984 | 1.2731 |
|  | Deletions | 65,549 | 613,150 | 16,882,129 |
|  | Insertions | 70,097 | 1,389,483 | 38,979,226 |
|  | Substitutions | 47,336 | 974,285 | 21,920,160 |
|  | Recall (%) | 99.9832 | 99.8136 | 99.4201 |
|  | Precision (%) | 99.9339 | 99.6100 | 98.7420 |
|  | Trimmed / split reads | 393 | 4,657 | 113,843 |
|  | Mean missing size (bp) | 65.3 | 78.9 | 126.3 |
|  | Extended reads | 0 | 2 | 273 |
|  | Mean extension size (bp) | 0 | 0 | 48 |
|  | Low quality reads | 35 | 214 | 1,263 |
|  | Short reads | 0 | 1 | 11 |
|  | Homopolymer ratio | 1.0000 | 1.0000 | 1.0160 |
|  | Runtime | 20 min | 1 h 09 min | 23 h 30 min |
|  | Memory (MB) | 457 | 794 | 2,332 |
| Proovread<sup>3</sup> | Number of bases | 259,647,167 | 671,307,477 | - |
|  | Error rate (%) | 0.1689 | 0.2568 | - |
|  | Deletions | 358,693 | 2,080,733 | - |
|  | Insertions | 2,837 | 56,965 | - |
|  | Substitutions | 15,499 | 102,130 | - |
|  | Recall (%) | 99.9948 | 99.9902 | - |
|  | Precision (%) | 99.8335 | 99.7467 | - |
|  | Trimmed / split reads | 18,693 | 68,598 | - |
|  | Mean missing size (bp) | 54.3 | 148.2 | - |
|  | Extended reads | 4 | 10 | - |
|  | Mean extension size (bp) | 20.2 | 20.2 | - |
|  | Low quality reads | 0 | 501 | - |
|  | Short reads | 172 | 7,658 | - |
|  | Homopolymer ratio | 1.0000 | 1.0000 | - |
|  | Runtime | 4 h 07 min | 11 h 51 min | - |
|  | Memory (MB) | 15,245 | 23,591 | - |
| Canu | Number of bases | 218,652,792 | 599,325,043 | 5,112,430,638 |
|  | Error rate (%) | 0.7404 | 0.7919 | 0.7934 |
|  | Deletions | 563,365 | 1,678,564 | 14,729,573 |
|  | Insertions | 827,596 | 2,412,967 | 20,996,517 |
|  | Substitutions | 72,521 | 240,442 | 1,897,014 |
|  | Recall (%) | 99.4781 | 99.4488 | 99.4573 |
|  | Precision (%) | 99.2658 | 99.2148 | 99.2131 |
|  | Trimmed / split reads | 27,690 | 73,001 | 569,082 |
|  | Mean missing size (bp) | 985.8 | 858.1 | 651.3 |
|  | Extended reads | 47 | 176 | 1,658 |
|  | Mean extension size (bp) | 29.0 | 53.7 | 37.2 |
|  | Low quality reads | 0 | 104 | 941 |
|  | Short reads | 187 | 354 | 1,845 |
|  | Homopolymer ratio | 1.0158 | 1.0108 | 1.0065 |
|  | Runtime | 24 min | 1 h 11 min | 9 h 30 min |
|  | Memory (MB) | 3,674 | 3,710 | 7,050 |
|  |  | *E. coli* | *S. cerevisiae* | *C. elegans* |
| *CONSENT* | Number of bases | 259,078,481 | 689,502,352 | 5,603,760,432 |
|  | Error rate (%) | 0.1722 | 0.2918 | 0.5730 |
|  | Deletions | 341,549 | 1,280,545 | 18,391,216 |
|  | Insertions | 55,787 | 582,869 | 12,737,445 |
|  | Substitutions | 8,354 | 115,614 | 2,826,988 |
|  | Recall (%) | 99.9843 | 99.9481 | 99.9119 |
|  | Precision (%) | 99.8306 | 99.7125 | 99.4345 |
|  | Trimmed / split reads | 27,380 | 71,027 | 571,107 |
|  | Mean missing size (bp) | 71.8 | 68.1 | 67.1 |
|  | Extended reads | 0 | 2 | 187 |
|  | Mean extension size (bp) | 0 | 0 | 122.2 |
|  | Low quality reads | 0 | 107 | 849 |
|  | Short reads | 0 | 0 | 0 |
|  | Homopolymer ratio | 1.0000 | 1.0000 | 0.9653 |
|  | Runtime | 36 min | 1 h 46 min | 27 h 04 min |
|  | Memory (MB) | 4,849 | 11,325 | 32,284 |
| *Daccord<sup>4</sup>* | Number of bases | 261,274,775 | 694,845,174 | - |
|  | Error rate (%) | 0.0214 | 0.0400 | - |
|  | Deletions | 6,353 | 51,217 | - |
|  | Insertions | 19,335 | 132,765 | - |
|  | Substitutions | 28,842 | 75,540 | - |
|  | Recall (%) | 99.9971 | 99.9928 | - |
|  | Precision (%) | 99.9790 | 99.9606 | - |
|  | Trimmed / split reads | 13 | 73 | - |
|  | Mean missing size (bp) | 58.8 | 276.8 | - |
|  | Extended reads | 0 | 4 | - |
|  | Mean extension size (bp) | 0 | 0 | - |
|  | Low quality reads | 0 | 113 | - |
|  | Short reads | 0 | 0 | - |
|  | Homopolymer ratio | 1.0000 | 1.0000 | - |
|  | Runtime | 54 min | 2 h 26 min | - |
|  | Memory (MB) | 18,450 | 32,190 | - |
| *FLAS* | Number of bases | 260,324,873 | 689,491,882 | 5,583,925,852 |
|  | Error rate (%) | 0.1547 | 0.2034 | 0.3997 |
|  | Deletions | 378,728 | 1,273,157 | 25,708,666 |
|  | Insertions | 53,059 | 220,507 | 5,628,454 |
|  | Substitutions | 3,141 | 30,665 | 1,249,658 |
|  | Recall (%) | 99.9546 | 99.9418 | 99.9175 |
|  | Precision (%) | 99.8526 | 99.8049 | 99.6104 |
|  | Trimmed / split reads | 403 | 1,274 | 30,950 |
|  | Mean missing size (bp) | 182.4 | 223.4 | 305.4 |
|  | Extended reads | 1 | 5 | 588 |
|  | Mean extension size (bp) | 102 | 102 | 156.7 |
|  | Low quality reads | 4 | 241 | 7,506 |
|  | Short reads | 0 | 0 | 14 |
|  | Homopolymer ratio | 0.9967 | 1.0000 | 0.9890 |
|  | Runtime | 38 min | 1 h 30 min | 10 h 45 min |
|  | Memory (MB) | 2,428 | 4,984 | 13,682 |
| *LoRMA* | Number of bases | 170,581,660 | 442,516,834 | 780,732,399 |
|  | Error rate (%) | 0.1285 | 0.2225 | 0.6446 |
|  | Deletions | 163,750 | 1,197,979 | 5,515,626 |
|  | Insertions | 25,117 | 129,027 | 369,539 |
|  | Substitutions | 21,604 | 106,535 | 581,292 |
|  | Recall (%) | 99.9865 | 99.9785 | 99.9547 |
|  | Precision (%) | 99.8743 | 99.7812 | 99.3649 |
|  | Trimmed / split reads | 40,347 | 108,516 | 1,154,643 |
|  | Mean missing size (bp) | 1,007 | 970.3 | 412.3 |
|  | Extended reads | 0 | 4 | 13 |
|  | Mean extension size (bp) | 0 | 0 | 685.7 |
|  | Low quality reads | 4 | 797 | 3,575 |
|  | Short reads | 187,376 | 522,390 | 9,256,368 |
|  | Homopolymer ratio | 1.0000 | 1.0000 | 1.0000 |
|  | Runtime | 1 h 39 min | 5 h 25 min | 31 h 04 min |
|  | Memory (MB) | 31,682 | 31,828 | 32,104 |
|  |  | *E. coli* | *S. cerevisiae* | *C. elegans* |
| *MECAT* | Number of bases | 232,519,767 | 616,247,287 | 4,937,676,257 |
|  | Error rate (%) | 0.1714 | 0.2088 | 0.2675 |
|  | Deletions | 392,085 | 1,228,519 | 12,514,560 |
|  | Insertions | 43,487 | 168,401 | 1,985,795 |
|  | Substitutions | 2,006 | 14,471 | 107,294 |
|  | Recall (%) | 99.9547 | 99.9428 | 99.9258 |
|  | Precision (%) | 99.8362 | 99.7996 | 99.7415 |
|  | Trimmed / split reads | 3,622 | 9,606 | 112,978 |
|  | Mean missing size (bp) | 365 | 380.7 | 566.8 |
|  | Extended reads | 0 | 0 | 30 |
|  | Mean extension size (bp) | 0 | 0 | 165 |
|  | Low quality reads | 0 | 92 | 443 |
|  | Short reads | 0 | 0 | 0 |
|  | Homopolymer ratio | 0.9967 | 1.0000 | 1.0000 |
|  | Runtime | 5 min | 16 min | 2 h 43 min |
|  | Memory (MB) | 2,387 | 4,954 | 10,563 |

**Table 11:**
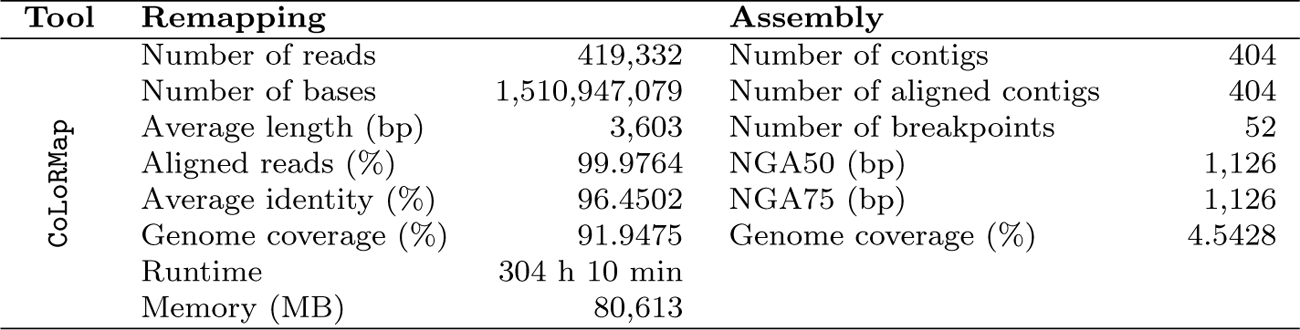
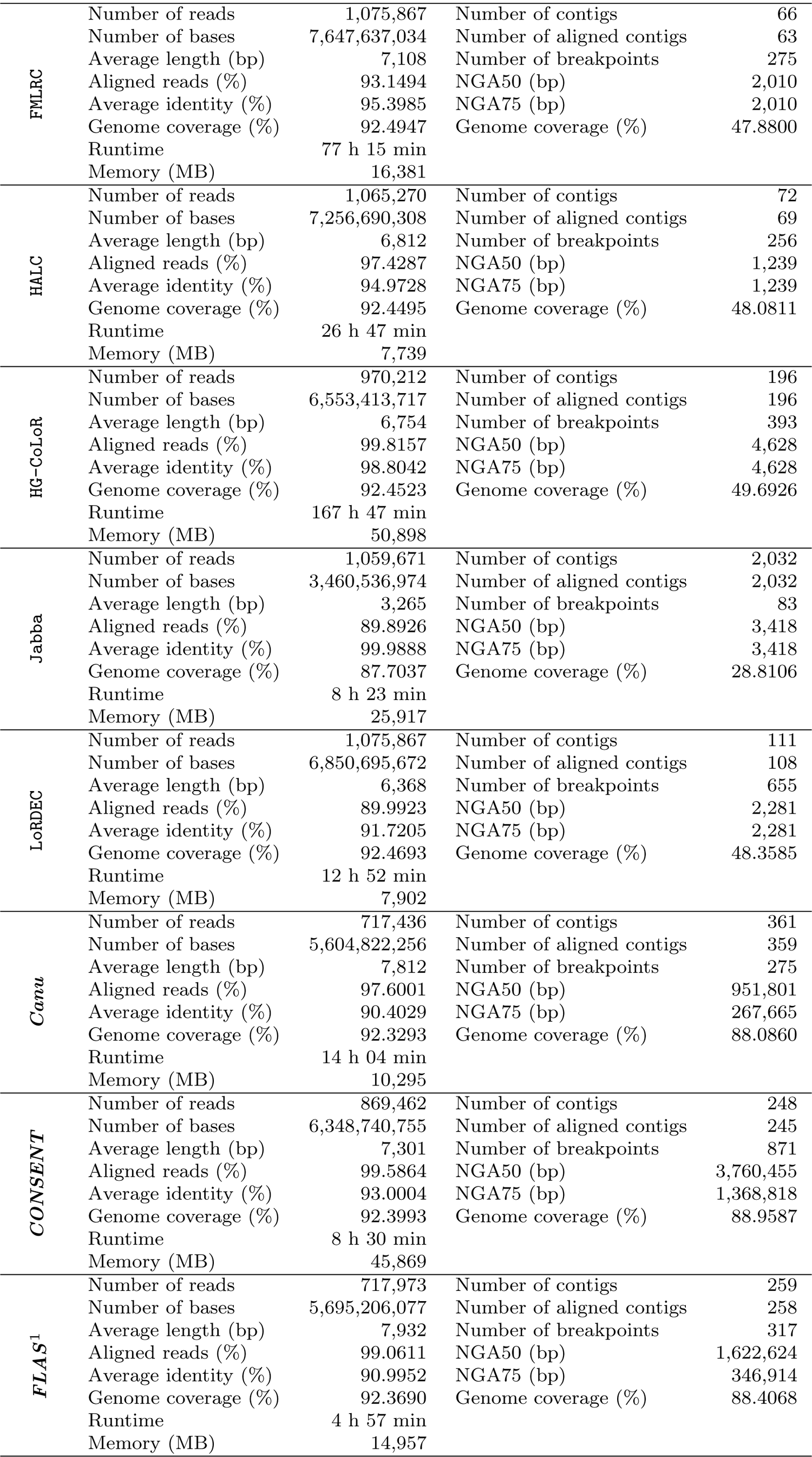
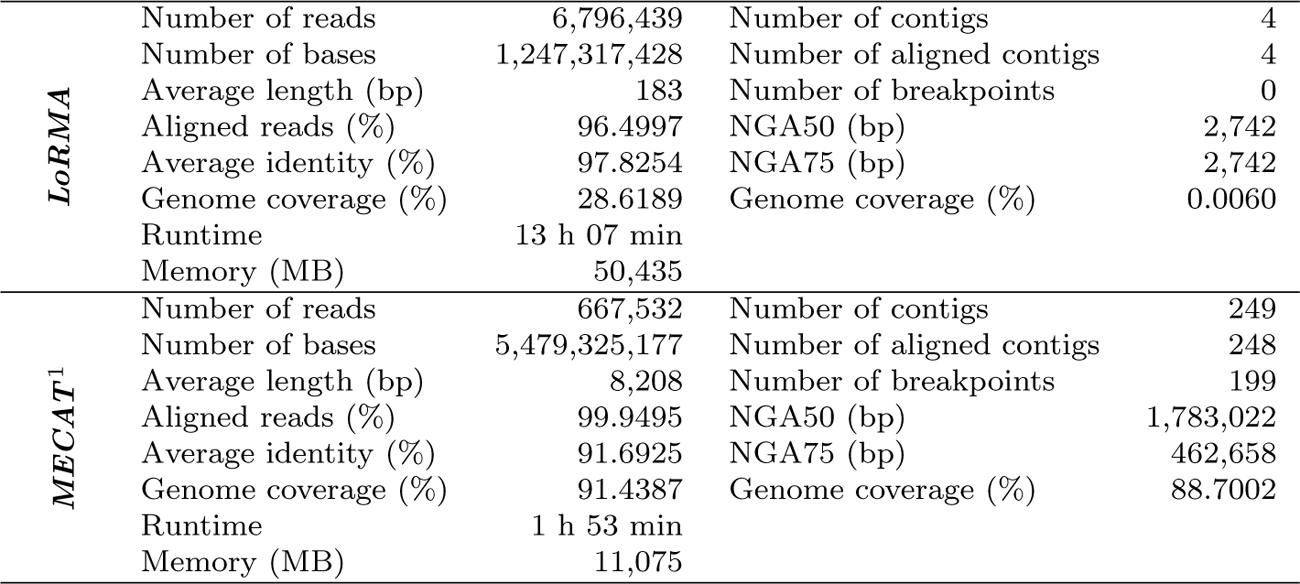
Comparison of the different error correction tools, on a dasaset containing ONT ultra-long reads. This corresponds to the *H. sapiens* dataset of Table 3. ^1^ Reads longer than 50 kbp were filtered out, since they caused the programs to stop with an error.

In terms of runtime, all hybrid correction tools except Jabba once again performed slower than self-correction methods. More precisely, Jabba was even the fastest tool on all these datasets, requiring at most 43 min to perform on the *C. elegans* dataset. ***MECAT***, the fastest self-correction tool nonetheless also performed correction fast, since it only required 48 min on that same dataset. Jabba excluded, however, self-correction methods performed faster than most hybrid correction methods. This can especially be seen on the *C. elegans* dataset, where ***CONSENT*** and ***Canu***, the two slowest self-correction tools, both required a little more than 9 hours to perform, while hybrid correction tools such as CoLoRMap and HG-CoLoR required respectively 4.5 and 6 days to perform. In terms memory consumption, self-correction tools were more demanding than most hybrid correction tools on all datasets. However, hybrid correction tools such as CoLoRMap, HG-CoLoR and Proovread required more memory than most self-correction tools, ***LoRMA*** excluded, on the *C. elegans* dataset.

These results thus suggest than, when error rates reach around 12% or less, self-correction methods can be satisfyingly applied, even when the sequencing depth only reaches 30x. Indeed, both self-correction and hybrid correction methods were comparable in terms of quality of the results, on all of these datasets. Moreover, these results also show that self-correction methods usually perform faster than hybrid correction methods, and that their memory consumption can also be lower on large datasets. These observations are especially interesting, since they suggest that, given the error rate of the reads is low enough, self-correction methods can provide a high-quality correction, even on low coverage datasets, and can thus allow to avoid the cost of having to sequence both long and short reads to perform efficient analysis. As a result, this suggests that self-correction is best suited, and should thus be preferred when analyzing such low error rates and low coverage datasets.

### 3.8 Low error rate, medium coverage

Here, we present results on the *E. coli* 60x (low), *S. cerevisiae* 60x (low) and *C. elegans* 60x (low) datasets of Table 3. For these datasets, when possible, we evaluated the trimmed corrected long reads. When trimmed reads were not available, we evaluated the complete corrected long reads. These results are summarized in Table 10 and commented below. Moreover, for these experiments, we excluded the following tools from the comparison, for the ensuing reasons:

- ECTools, because of its unsatisfying results on previous experiments;
- Hercules, because of its unsatisfying results and large runtimes on previous experiments;
- LSC, because of its unsatisfying results on previous experiments;
- Nanocorr, because of its large runtimes on previous experiments;
- NaS, because of its large runtimes on previous experiments;

On these datasets, both hybrid and self-correction methods did manage to reduce the error rate to around 1% or less, on all datasets. More precisely, only the hybrid methods FMLRC and LoRDEC reached an error rate of more than 1% on the *C. elegans* dataset. In comparison, on this dataset, all self-correction tools managed to reduce the error rate well below 1%, the highest error rate being that of ***Canu*** on the *C. elegans* dataset, and only reaching 0.79%. This observation is in accordance with that previously made in Section 3.7, but also further shows that self-correction methods provide better correction as the sequencing depth grows. Moreover, it is also interesting to note that, like in Section 3.7, most self-correction tools reported as many corrected bases as hybrid methods, although ***LoRMA*** still performed poorly regarding this metric, and ***MECAT*** still reported slightly less corrected bases than other methods. Overall, hybrid correction and self-correction methods were thus highly comparable in terms of quality of the results on these three datasets, and self-correction methods even performed better than in Section 3.7. This underlines the fact that, given the error rate of the reads reaches around 12%, self-correction methods perform well on low coverage dataset, but also benefit from larger sequencing depths, providing higher quality correction as coverage grows higher.

In terms of runtime, all hybrid correction tools except Jabba once again performed slower than self-correction methods. More precisely, Jabba even was the fastest tool on all these datasets, requiring at most 49 min to perform on the *C. elegans* dataset, while ***MECAT***, the fastest self-correction tool, required 2 h 43 min. Jabba excluded, however, self-correction methods performed faster than most hybrid correction methods. This can especially be seen on the *C. elegans* dataset, where hybrid correction tools such as CoLoRMap, HG-CoLoR and Proovred were not run due to their unreasonable runtimes. In comparison, ***LoRMA*** and ***CONSENT***, the slowest self-correction tools on this dataset, required respectively 31 hours and 27 hours to perform, while other self-correction tools required 10 hours or less. In terms of memory consumption, self-correction tools were once again more demanding than most hybrid correction tools on all datasets, although hybrid tools such as CoLoRMap and Proovread required more resources than all other tools except ***LoRMA*** and ***Daccord***.

These observations are in accordance with conclusions drawn in Section 3.7, indicating that, when error rates reach around 12% or less, self-correction methods perform as well as hybrid correction methods, or even outperform them on complex datasets. However, this results also show that self-correction methods can also benefit from higher sequencing depth, and manage to perform better as it grows larger. Moreover, these results further show that self-correction methods perform faster than hybrid correction methods, even more so when the sequencing depth is large. As a result, this suggests that self-correction is best suited, both in terms on quality of the results and in terms of resource consumption, and should thus be preferred when analyzing low error rates datasets, especially when the sequencing depth grows large.

### 3.9 Ultra-long reads

Here, we present results on the *H. sapiens* dataset of Table 3. For this dataset, when possible, we evaluated the trimmed corrected long reads. When trimmed reads were not available, we evaluated the complete corrected long reads. These results are summarized in Table 11 and commented below. Moreover, for these experiments, we excluded the following tools from the comparison, for the ensuing reasons:

- ECTools, because of its unsatisfying results on previous experiments;
- Hercules, because of its unsatisfying results and large runtimes on previous experiments;
- LSC, because of its unsatisfying results on previous experiments;
- Nanocorr, because of its large runtimes on previous experiments;
- NaS, because of its large runtimes on previous experiments;
- Proovread, because it could not be installed on the computer we performed these experiments on;
- ***Daccord***, because it could not perform correction, since the alignment step with DALIGNER required more than 128 GB of memory.

On this dataset, most tools, both hybrid and non-hybrid, failed to efficiently reduce the error rate of the reads to 1% or less. More precisely, the two best performing tools were Jabba and HG-CoLoR, which respectively reached an error rate of 0.1% and 1.2%. However, it is worth noting that Jabba output two times less bases than HG-CoLoR, and that the average length of its corrected reads only reached 3,265, indicating that it split a large proportion of reads, and thus generated high quality, but short fragments. In the same fashion, CoLoRMap also produced corrected reads reaching a short average length, despite the fact it did not trim or split the reads. This tends to indicate CoLoRMap mostly managed to correct the shortest reads of the dataset. The fact that hybrid correction methods did not manage to perform well on this dataset is consistent with previous observations indicating that they tend to suffer from performance drops when applied to large and complex genomes. However, it is still interesting to note that HG-CoLoR performed well on this dataset, managing to output a large number of high-quality corrected reads.

As for self-correction, it worth noting that ***FLAS*** and ***MECAT*** could not perform correction on these ultra-long reads, and that they had to be filtered out in order to allow the methods to run. In terms of quality of the results, ***LoRMA*** reached the lowest error rate, of only 2.2%, but reported reads with an average length of 183 bp, and thus aggressively trimmed the reads once again. Other self-correction tools globally performed worse than hybrid correction tools in terms of reduction of the error rate, the best performing tool being ***CONSENT***, and only reducing the error rate to 7%. It is however worth noting that the average length of the corrected reads was larger for self-correction than for hybrid correction methods, which indicates that self-correction methods trimmed a smaller proportion of the reads, and thus explains the differences in terms of error rates.

In terms of assembly, Jabba and especially ***LoRMA*** performed the worst, owing to the fact they split a large proportion of the reads. In the same fashion, CoLoRMap also yielded a low quality assembly, due to the fact its corrected reads reached a small average length. More precisely, the assembly yielded from Jabba corrected reads was composed of more than 2,000 contigs, and only covered 28.8% of the genome, the assembly yielded from CoLoRMap corrected reads was composed of 404 contigs, and only covered 4.5% of the genome, while ***LoRMA*** corrected reads yielded an assembly composed of only 4 contigs, which covered less than 0.1% of the genome. More generally, self-correction methods yielded higher quality assemblies, that, despite being composed of a larger number of contigs than assemblies obtained from reads corrected with hybrid methods, CoLoRMap excluded, covered largest proportion of the genome. Indeed, while most assemblies yielded from reads corrected by hybrid methods covered less than 50% of the genome, self-correction methods managed to produce corrected reads yielding assemblies covering around 88% of the genome. Once again, this can be explained by the performance drops hybrid methods suffer from when applied to large and complex datasets, and by the fact that self-correction methods benefit from the whole long-range information of the reads when performing correction.

In terms of runtime, most hybrid correction tools performed slower than self-correction methods, although Jabba was faster than all self-correction methods but ***FLAS*** and ***MECAT***, and LoRDEC was faster than ***Canu*** and ***LoRMA***. The fastest tool on this dataset was ***MECAT***, which managed to perform correction in under 2 hours. Conversely, the slowest tool was CoLoRMap, requiring more than 12 days to perform, while the slowest self-correction tool, ***Canu***, only required 14 hours. In terms on memory consumption, both hybrid and self-correction tools were comparable, with self-correction tools ***CONSENT*** and ***LoRMA*** requiring respectively 45 GB and 50 GB, and being comparable to hybrid correction tool HG-CoLoR, also requiring 50 GB. It is however worth noting that the most memory consuming method was the hybrid tool CoLoRMap, requiring up to 80 GB. Conversely, the least consuming hybrid correction tools were HALC and LoRDEC, both requiring a little less than 8 GB, while the least consuming self-correction tools were ***Canu*** and ***MECAT***, requiring respectively 10 GB and 11 GB.

The results further confirm conclusions drawn in Section 3.5.1, 3.6 and 3.7, and indicating that hybrid correction methods suffer from performance drops when the complexity of the studied organism grows. This can especially be seen on the assembly metrics of this dataset, where self-correction methods largely outperformed hybrid correction tools. Despite this fact, it is nonetheless worth noting that, in pure terms of reduction of the error rate, hybrid tool HG-CoLoR outperform all other methods. Moreover, it is also interesting to note that, while all hybrid tools could directly be applied to this ultra-long reads dataset, self-correction methods ***FLAS*** and ***MECAT*** required these ultra-long reads to be filtered out in order to run, although this did not impact their performances. As a result, a hybrid tool such a HG-CoLoR could be preferred when the aim is to lower the error rate as much as possible, even though it is much more time consuming than self-correction tools, which are not only faster, but should also be privileged when downstream analyses include assembly.

## 4 Conclusion

In this paper, we presented an in-depth description of the state-of-the-art on long-read error correction, tackling both hybrid and self-correction. For each of these two approaches, we thoroughly described the different existing methodologies. On the one hand, for hybrid correction, these methodologies include short reads alignment, contigs alignment, de Bruijn graphs usage, as well as Hidden Markov Models usage, although the latter is not as widespread as the three other ones. On the other hand, for self-correction, these methodologies are mainly restricted to multiple sequence alignment and de Bruijn graphs usage.

More precisely, this paper draws an exhaustive characterization of a total of thirty different error correction methods, relying on the different aforementioned methodologies, or on combinations of these methodologies. For each of these thirty tools, we precisely describe the way they perform correction, the manner in which they employ the aforementioned methodologies, the algorithmic specificities they rely on, as well as the specificities of the corrected reads they output, that is to say, whether they report corrected bases in a different case, or perform trimming or splitting of uncorrected reads regions. As a result, we present the most comprehensive survey on long-read error correction up to date.

Additionally, we also gathered a total of 19 different datasets, composed of both simulated and real data, both PacBio and ONT data, and displaying various error rates, sequencing depths, read lengths, and ranging from small bacterial to large mammal genomes, in order to precisely assess the efficiency of all the tools we described. This way, we not only provide a complete description of existing tools, but also the most comprehensive benchmark on long-read error correction up to date.

These experiments allowed us to study the precise behavior of the different tools and methodologies according to the characteristics of the long-reads datasets. For instance, they show that de Bruijn graphs based methods tend to split a large proportion of the reads, when they do not manage to find paths, but can also extend the reads beyond their initial extremities when non-branching paths can be followed. More practically, these experiments also allow us to provide a few guidelines regarding the best performing tools according to the datasets characteristics. More precisely, these results show that, when the error rate of the reads is high, above 30%, hybrid correction methods largely outperform self-correction methods, even when the sequencing depth is high. However, when the error rate is moderately high, around 18%, our experiments show that hybrid correction outperforms self-correction when the sequencing depth is low, but that self-correction tends to be more efficient as the sequencing depth grows. Finally, when the error rates are much lower, and reach 12% or less, our experiments show that self-correction performs best, even on low coverage datasets. Additionally, the complexity of the studied organisms was shown to have drastic impacts on the quality of the correction. More precisely, hybrid correction methods suffered from severe performance drops when applied to more complex and repeat-rich organisms, while the impact on self-correction methods was more balanced. Our experiments show that this especially impacts assembly results, and metrics reported on a *H. sapiens* dataset showed that all self-correction methods largely outperformed hybrid correction in terms of assembly quality. It is also worth noting that all our experiments were performed using default parameters. Fine tuning of the parameters for each tool might thus slightly improve their performances. However, our experiments and results nonetheless give a reliable trend as to which tool or type of tool performs best.

Although our work represents the most comprehensive survey and benchmark on long-read error correction up to date, further work shall focus on the study of additional datasets. In particular, the error rates of the datasets we studied here no longer properly reflect the error rates of both PacBio and ONT latest sequencing experiments, which now reach around 8% or less. However, since results on 12% error rates datasets indicate that self-correction is best suited, the further reduction of the error rates should only further confirm our conclusions. Another direction would also be to study the impact on error correction on other applications such as variant calling. Finally, we are also interested in studying the behavior of the different tools and methodologies on polyploid organisms, since there literature lacks of application of long-read error correction tools on such organisms.

## Acknowledgments

Part of this work was performed using computing resources of CRIANN (Normandy, France), project 2017020.

## Footnotes

1 http://www.novocraft.com/products/novoalign

2 https://github.com/biointec/brownie

3 https://hazelcast.com/

4 http://zombie.cb.k.u-tokyo.ac.jp/sprai/index.html

