## Supplementary Materials for "Long-read error correction: a survey and qualitative comparison"

#### Supplementary material

Pierre Morisse[1,*], Thierry Lecroq[2] and Arnaud Lefebvre[2]

[1]Univ Rennes, Inria, CNRS, IRISA, 35000, Rennes, France
[2]Normandie Univ, UNIROUEN, LITIS, 76000 Rouen, France

### 1 Software versions and command lines

#### 1.1 Hybrid correction tools

##### 1.1.1 CoLoRMap

We used CoLoRMap commit 24a9626. The following command lines were used for both PacBio and ONT data.

- `runCorr.sh longReads.fasta shortReads.fastq outputDirCorr \\ outputPrefixCorr 16`
- `runOEA.sh outputDirCorr/outputPrefixCorr_sp.fasta \\ shortReads.fastq outputDirOEA outputPrefixOEA 16`

##### 1.1.2 ECTools

ECTools requires to run a series of commands and external programs in order to run. We provide the versions of each program and the complete suite of command lines below.

**Versions**

- Minia 3 commit 0b52222.
- ECTools commit 031eb03.

**Command lines** The following command lines were used for both PacBio and ONT data.

- `mkdir outputDir`
- `cd outputDir`
- `minia -in shortReads.fasta`
- `mv shortReads.unitigs.fasta unitigs.fasta`
- `python ../partition.py 20 500 longReads.fasta`
- `for i in 0001..000N; do cd $i; for j in 1..500; do echo \\ "SGE_TASK_ID=$j TMPDIR=. ../correct.sh"; done | parallel \\ -j 16 ; cd ..; done` (where $N$ is the number of partitions)

##### 1.1.3 FMLRC

We used FMLRC commit b6c5d45. The following command lines were used for both PacBio and ONT data.

- `awk "NR % 2 == 0" shortReads.fasta | sort -T . | tr NT TN \\`
  `| ./ropebwt2/ropebwt2 -LR | tr NT TN \\`
  `| fmlrc-convert shortReads.npy`
- `fmlrc -p 16 shortReads.npy longReads.fasta \\`
  `correctedReads.fasta`

##### 1.1.4 HALC

HALC requires to run a series of commands and external programs in order to run. We provide the versions of each program and the complete suite of command lines below.

###### Versions

- Minia 3 commit 0b52222.
- HALC commit a106f78.

###### Command lines

The following command lines were used for both PacBio and ONT data. After running, results are stored in the `output` folder.

- `minia -in shortReads.fasta`
- `python runHALC.py -o shortReads -t 16 longReads.fasta \\`
  `shortReads.contigs.fa`

##### 1.1.5 Hercules

We used Hercules commit e0a7b39. The following command lines were used for both PacBio and ONT data.

- `mkdir outputDir`

- `hercules -1 -li longReads.fasta -si shortReads.fastq \\`
  `-o outputDir/`
- `./utils/runBowtieRmDup.sh \\`
  `outputDir/compressed_longReads.fasta \\`
  `outputDir/compressed_short.fasta .  16`
- `./utils/afteralignment.sh alignment.bam \\`
  `output_alignment.bam 16 32G`
- `hercules -2 -li longReads.fasta -ai output_alignment.bam \\`
  `-si outputDir/short.fasta -t 16 -o correctedReads.fasta`

###### 1.1.6 HG-CoLoR

We used HG-CoLoR commit 78d8ec6. The following command line was used for both PacBio and ONT data. After running, results are stored as `correctedReads.fasta`, `correctedReads.trim.fasta` and `correctedReads.split.fasta`.

- `HG-CoLoR -K 100 -k 40 -longreads longReads.fasta \\`
  `-shortreads shortReads.fastq -out correctedReads`

###### 1.1.7 Jabba

Jabba requires to run a series of command and other programs in order to run. We provide the versions of each program and the complete suite of command lines below. Although Jabba authors recommend correcting the reads with Karect, any tool can be used, and we used QuorUM, which is faster and provides similar results.

**Versions**

- QuorUM v1.1.1.
- Brownie commit abaf5ff.
- Jabba v1.0.0.

**Command lines** The following command lines were used for both PacBio and ONT data.

- mkdir correctedShortReads
- cd correctedShortReads
- quorum -q 22 shortReads.fastq
- mv quorum_corrected.fa quorum_corrected.fasta
- cd ..
- mkdir brownieOutput
- brownie graphCorrection -p brownieOutput -k 75 \\ correctedShortReads/quorum_corrected.fasta
- jabba -o jabbaOutputDir -k 75 -g brownieOutput/DBGraph.fasta \\ -fasta longReads.fasta

###### 1.1.8 LoRDEC

We used LoRDEC v0.6. The following command lines were used for both PacBio and ONT data.

**For bacterial genomes**

- `lordec-correct -T 16 -i longReads.fasta \\`
  `-2 shortReads.fasta -k 19 -o correctedReads.fasta`

**For other genomes**

- `lordec-correct -T 16 -i longReads.fasta \\`
  `-2 shortReads.fasta -k 23 -o correctedReads.fasta`

###### 1.1.9 LSC

We used LSC v2.0. The following command line was used for both PacBio and ONT data.

- `python runLSC.py -long_reads longReads.fasta -short_reads \\`
  `shortReads.fasta -threads 16 -specific_tempdir tmpDir \\`
  `-o outputDir`

##### 1.1.10 Nanocorr

We used Nanocorr commmit c11a885. The following command lines were used for both PacBio and ONT data.

- `python partition 100 500 longReads.fasta`
- ```
  for f in $(ls -d 0*); do
      cd $f
      for j in {0..500}; do
          echo "SGE_TASK_ID=$j TMPDIR=. ../nanocorr.py \\
          ../shortReads.fasta";
      done  | parallel -j 16
      cd ..
  done
  ```

### 1.1.11 NaS

We used NaS v1.0. The following command line was used for both PacBio and ONT data.

- ```
  NaS_v2/NaS_wrapped -fq1 shortReads_1.fastq \\
  -fq2 shortReads_2.fastq -nano longReads.fastq \\
  -out outputDir -mode fast -nb_proc 16
  ```

##### 1.1.12 Proovread

We used Proovread v2.14.1. The following command line was used for both PacBio and ONT data.

- ```
  proovread -t 16 -overwrite -long-reads longReads.fasta \\
  -short-reads shortReads.fasta -p outputDir
  ```

## 1.2 Self-correction tools

### 1.2.1 Canu

We used Canu v2.0. The following command lines were used.
  ```

##### For PacBio data

- `canu -correct -p ResultsPrefix -d ResultsDirectoy \\`
  `genomeSize=exactSizeOfTheGenome \\`
  `-pacbio-raw longReads.fasta -stopOnReadQuality=false \\`
  `-corOutCoverage=300 -useGrid=false`

##### For ONT data

- `canu -correct -p ResultsPrefix -d ResultsDirectory\\`
  `genomeSize=exactSizeOfTheGenome \\`
  `-nanopore-raw longReads.fasta -stopOnReadQuality=false \\`
  `-corOutCoverage=300 -useGrid=false`

##### 1.2.2 CONSENT

We used consent commit 39c5302. The following command lines were used.

##### For PacBio data

- `CONSENT-correct -in longReads.fasta \\`
  `-out correctedReads.fasta -type PB`

##### For ONT data

- `CONSENT-correct -in longReads.fasta \\`
  `-out correctedReads.fasta -type ONT`

##### 1.2.3 Daccord

Daccord requires to run a series of scripts and other programs in order to run. We provide the versions of each subprogram and the complete suite of command lines below. ConverToPacBio_q2a.py, fasta2DB, and DBsplit are provided within the DAZZ_DB suite.

##### Versions

- DAZZ_DB version July 17, 2013.
- DALIGNER version April 10, 2016.
- Daccord version v0.0.10.

**Command lines** The following command lines were used for both PacBio and ONT data.

- `python ConverToPacBio_q2a.py longReads.fasta` (This command creates, by default, a formatted file called "LR.fasta", which we use as an input for the following steps)
- `fasta2DB readsDb LR.fasta`
- `DBsplit -x14 readsDb.db`
- `daligner -T16 readsDb.db readsDb.db`
- 
```
daccord -t16 readsDb.readsDb.las readsDb.db \\
> correctedReads.fasta
```

###### 1.2.4 FLAS

We used FLAS commit 053c19b. The following command line was used for both PacBio and ONT data.

- `python runFLAS.py longReads.fasta`

###### 1.2.5 LoRMA

We used LoRMA v0.4. The following command line was used for both PacBio and ONT data.

- `lorma.sh -s -threads 16 longReads.fasta`

###### 1.2.6 MECAT

We used MECAT commit d04bfa8. The following command lines were used.

**For PacBio data**

- 
```
mecat2pw -j 0 -d longReads.fasta -w .  -t 16 \\
-o longReads.fasta.can
```
- 
```
mecat2cns -i 0 -t 16 longReads.fasta.can longReads.fasta \\
correctedReads.fasta
```

**For ONT data**

- `mecat2pw -j 0 -d longReads.fasta -o candidates.txt -w .  \\`
  `-t 28 -x 1`

- `mecat2cns -i 0 -t 28 -x 1 candidates.txt longReads.fasta \\`
  `correctedReads.fasta`
